# From bacteria to plants: a repurposing strategy in the pursuit for novel herbicides

**DOI:** 10.1101/2022.02.20.481218

**Authors:** Emily R. R. Mackie, Andrew S. Barrow, Marie-Claire Giel, Mark D. Hulett, Anthony R. Gendall, Santosh Panjikar, Tatiana P. Soares da Costa

## Abstract

Herbicide resistance represents one of the biggest threats to our natural environment and agricultural sector. Thus, new herbicides are urgently needed to tackle the rise in herbicide-resistant weeds. Here, we employed a novel strategy to repurpose a ‘failed’ antibiotic into a new and target-specific herbicidal compound. Specifically, we identified an inhibitor of bacterial dihydrodipicolinate reductase (DHDPR), an enzyme involved in lysine biosynthesis in plants and bacteria, that exhibited no antibacterial activity but severely attenuated germination of the plant *Arabidopsis thaliana*. We confirmed that the inhibitor targets plant DHDPR orthologues *in vitro*, and exhibits no toxic effects against human cell lines. A series of analogues were then synthesised with improved efficacy in germination assays and against soil-grown *A. thaliana* plants. We also showed that our lead compound is the first lysine biosynthesis inhibitor with herbicidal activity against a weed species, providing proof-of-concept that DHDPR inhibition may represent a much-needed new herbicide mode of action. Furthermore, this study exemplifies the untapped potential of repurposing ‘failed’ antibiotic scaffolds to fast-track the development of herbicide candidates targeting the respective plant enzymes to combat the global rise in herbicide-resistant weeds.

---

Herbicides play an integral role in modern agricultural practices as they enable the cost-effective management of weeds.^1^ However, herbicide options are dwindling due to the rapid emergence and spread of herbicide-resistant weed populations. Such weeds aggressively compete with crops for resources, resulting in decreased harvest yields and quality. The diminishing efficacy of current herbicides, coupled with the lack of new herbicides with novel modes of action over the last 30 years, has prompted serious concerns over sustainable agriculture.^2,3^ Consequently, there is an urgent need for the development of new herbicides, especially those with new modes of action.

The biosynthetic pathways that lead to the production of amino acids in plants have long been targeted for herbicide discovery, with great commercial success.^4^ The most widely used herbicide, glyphosate (the active ingredient in Roundup®), targets the production of aromatic amino acids through inhibition of the enzyme 5-enolpyruvyl-shikimate 3-phosphate synthase (EPSPS, EC 2.5.1.19).^4^ Similarly, herbicides that inhibit the biosynthesis of glutamine (e.g. glufosinate) and branched chain amino acids (e.g. chlorsulfuron) target a single enzyme within each pathway and have become indispensable to agricultural industries.^4^ Underpinning the success of these herbicides is the essentiality of amino acids for physiological processes, including protein synthesis, carbon and nitrogen metabolism, and the production of secondary metabolites.^5^ Given that plants can synthesise all amino acids, arresting their production represents an excellent herbicide development strategy. As such, we proposed that the unexplored diaminopimelate (DAP) pathway, which is responsible for lysine biosynthesis exclusively in plants, bacteria and algae, represents a potential herbicide target (Figure 1).^4,6^ Furthermore, we have recently identified the first lysine biosynthesis inhibitors with herbicidal activity against the model plant *Arabidopsis thaliana,* which target the first enzyme in the DAP pathway, dihydrodipicolinate synthase (DHDPS, EC 4.3.3.7) (Figure 1).^7–9^

**Figure 1.**
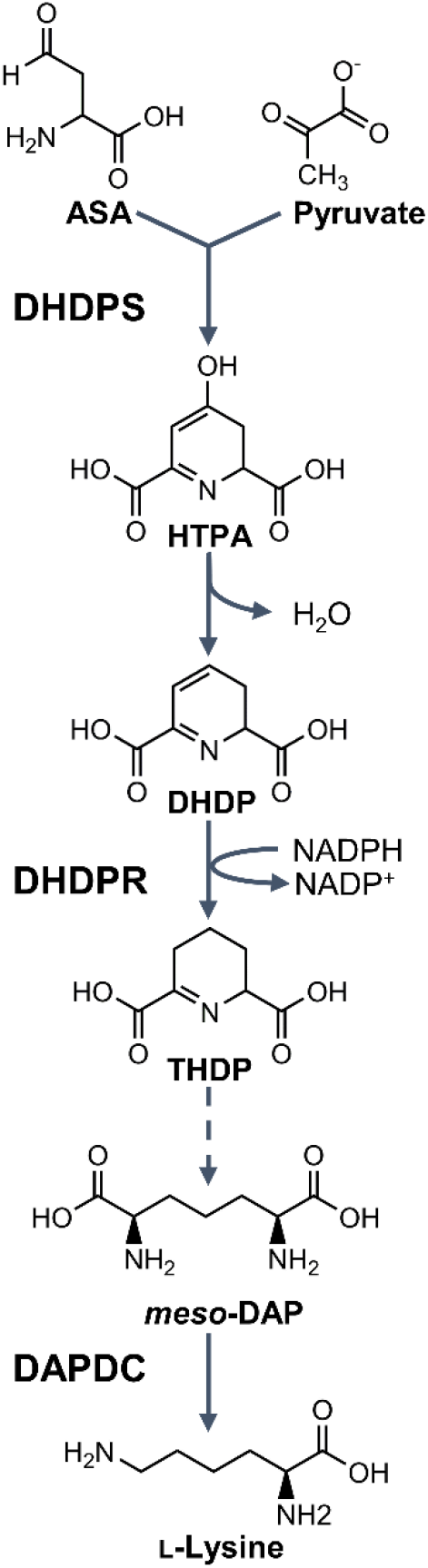
The diaminopimelate (DAP) pathway. The dihydrodipicolinate synthase (DHDPS)-catalysed condensation of aspartate semialdehyde (ASA) and pyruvate yields 4-hydroxy-2,3,4,5-tetrahydro-dipicolinic acid (HTPA), which is non-enzymatically dehydrated to form dihydrodipicolinate (DHDP). The NADPH-dependent reduction of DHDP to yield 2,3,4,5-tetrahydrodipicolinate (THDP) is catalysed by dihydrodipicolinate reductase (DHDPR). *meso*-DAP is eventually produced via one of four species-dependent sub-pathways, which is decarboxylated by DAP decarboxylase (DAPDC) to form L-lysine.

Despite the potential of targeting lysine biosynthesis production in plants as a strategy for herbicide development, little research has been done to date. Conversely, over the past 30 years many studies have focused on the development of antibiotics by inhibiting bacterial lysine biosynthesis enzymes.^10–16^ However, *in vitro* inhibitors of the bacterial enzymes are not effective against intact pathogenic bacteria, and hence, they have not progressed through the antibiotic development pipeline.^10,13^ These compounds are typically small molecules with MW <350 g·mol^-1^, the size of nearly all commercial herbicides to date. Moreover, plant enzymes in the DAP pathway are closely related to the bacterial orthologues and are essential for plant survival.^7,17,18^ Given the high degree of similarity between these enzymes from bacteria and plants, we explored the possibility that these ‘failed’ antibiotics could be repurposed into inhibitors of the respective plant enzymes. This strategy would circumvent the laborious screening of chemical libraries with unknown targets typically used in herbicide discovery, and therefore provide a fast-tracked method to develop urgently needed novel herbicide modes of action.

This study focuses on the second enzyme in the DAP pathway, dihydrodipicolinate reductase (DHDPR, EC 1.17.1.8) (Figure 1). Although there are no published plant DHDPR inhibitors, there are examples of inhibitors of the bacterial orthologues.^10,19^ The most well-characterised is 2,6-pyridine dicarboxylic acid (2,6-PDC), which has mid-micromolar potency against several bacterial DHDPR enzymes, including that from *Escherichia coli* (Ec), but with no antibacterial activity reported.^10,19^ Upon the recent publication of the first plant DHDPR structure, we postulated that 2,6-PDC may also bind to and inhibit plant DHDPR enzymes.^20^

Here, we sought to explore the evolutionary relationship between bacterial and plant DHDPR enzymes to repurpose 2,6-PDC as a potential herbicidal scaffold. To achieve this, we recombinantly produced two DHDPR enzymes from *A. thaliana,* AtDHDPR1 and AtDHDPR2. Subsequently, we characterised AtDHDPR1 functionally and structurally using enzyme kinetics assays and X-ray crystallography, and compared it to the previously characterised AtDHDPR2 and EcDHDPR enzymes. We confirmed that 2,6-PDC displays micromolar inhibition against AtDHDPR1 and AtDHDPR2 *in vitro* using enzyme kinetics assays, and is able to inhibit the germination of *A. thaliana* plants. To confirm its specificity, we employed antibacterial and cytotoxicity assays and determined that 2,6-PDC lacks activity against soil microbes and human cells. Finally, a series of analogues of 2,6-PDC were synthesised that had improved potency in germination assays, and when applied to soil-grown plants. Importantly, our lead inhibitor displayed herbicidal activity against the invasive weed species rigid ryegrass (*Lolium rigidum*).

## RESULTS

### Production of recombinant AtDHDPR proteins

The AtDHDPR1-encoding gene At2G44040 was identified using The Arabidopsis Information Resource (TAIR, https://www.arabidopsis.org/) and the resulting protein sequence uploaded to the ChloroP server for identification of the chloroplast transit peptide (cTP). ChloroP predicted a cTP length of 53 amino acids, however the final two amino acids were excluded based on the sequence of the previously characterised AtDHDPR2. Thus, the final construct was designed to exclude the first 51 amino acids and incorporate a custom fusion tag (Met-6×His-3C protease recognition site) for purification by immobilised metal affinity chromatography (IMAC) and tag removal (Supplementary Figure 1). A similar strategy was used to produce AtDHDPR2. The protein sequence resulting from the AtDHDPR2-encoding gene At3G59890 was used to predict a cTP length of 53 amino acids, which were excluded from the construct. Subsequent CD spectroscopy analysis revealed a similar secondary structure composition of 51% and 59% α/β structure for AtDHDPR1 and AtDHDPR2, respectively (Supplementary Figure S2). These results are in agreement with previous studies of bacterial and cyanobacterial orthologues, indicating correct protein folding.^21,22^

### Catalytic activity of AtDHDPR1

Having determined that AtDHDPR1 is folded similarly to AtDHDPR2, the kinetic properties of the enzyme were determined using a well-established DHDPS-DHDPR coupled assay.^23^ The best fits were obtained with a substrate inhibition model, consistent with inhibition by DHDP when using NADPH as the cofactor, which has been reported for AtDHDPR2 and other orthologues (Figure 2).^20,22,24^ AtDHDPR1 has a *k*_cat_ of 27 s^-1^, *K*_M_(DHDP) of 37 ± 6.5 μM and *K*_M_(NADPH) of 16 ± 2.6 μM. These kinetic constants are similar to the previously reported values for AtDHDPR2 of a *K*_M_(DHDP) of 57 μM and *K*_M_(NADPH) of 35 μM.^24^

**Figure 2.**
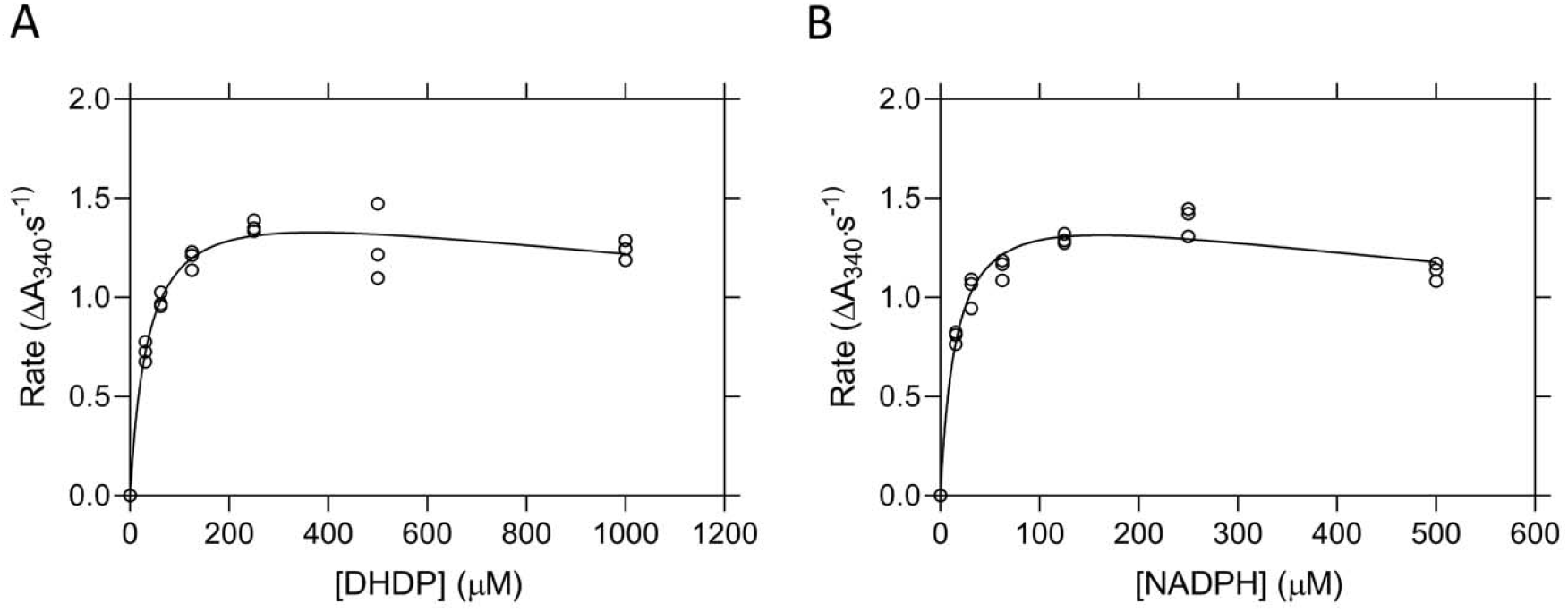
Kinetic analyses of *Arabidopsis thaliana* (At) DHDPR1. Initial rate (○) plotted as a function of (A) DHDP or (B) NADPH concentration. The nonlinear best fits (—) were obtained to a substrate inhibition model resulting in R^2^ values of 0.97 and 0.96, respectively. *n* = 3.

### Structural determination of AtDHDPR1 and comparison with AtDHDPR2 isoform

Given that plants generally possess two DHDPR isoforms, we investigated the similarity between paralogous plant DHDPR isoforms by determining the crystal structure of AtDHDPR1 and comparing it to the previously characterised AtDHDPR2 structure (PDB ID: 5UA0). The AtDHDPR1 structure has an N-terminal Rossmann-fold that is typically observed in DHDPR enzymes and a C-terminal oligomerisation domain that differs between species.^17^ AtDHDPR1 was crystallised as a monomer in the asymmetric unit, however based on symmetry operations was predicted to assemble as a weak tetramer, or ‘dimer of dimers,’ with a tight dimerisation interface. This is consistent with analytical ultracentrifugation analyses, which show that AtDHDPR1 exists in a dimer-tetramer equilibrium in solution (Supplementary Figure S3). As has been observed for AtDHDPR2, AtDHDPR1 was crystallised with the ‘latch and catch’ residues Met146, Gln145 and Thr189 within hydrogen bonding proximity (Figure 3).^17^ Interactions between these residues stabilise the enzyme’s closed conformation by pulling the N-terminal domain towards the C-terminal domain.^17^

**Figure 3.**
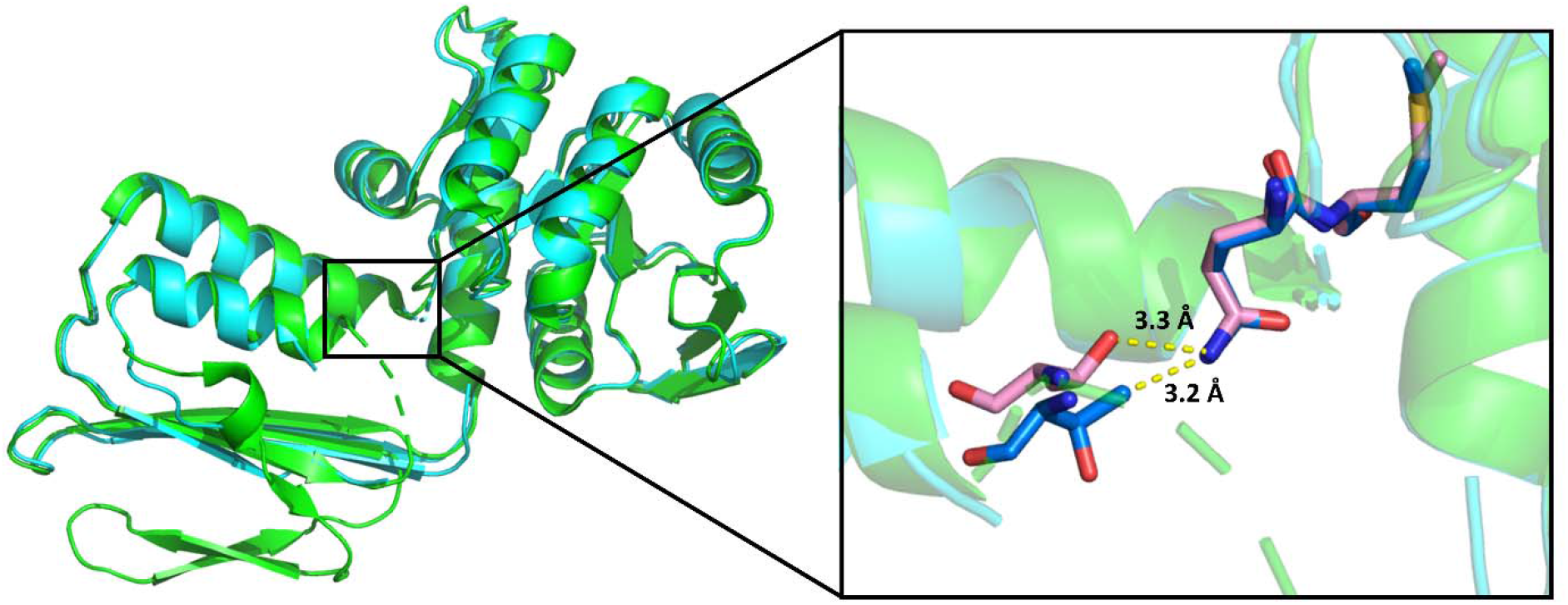
Structural alignment of AtDHDPR1 and AtDHDPR2. The monomeric crystal structure of AtDHDPR1 (cyan, PDB ID: 7T34) aligned with chain C of the dimeric AtDHDPR2 crystal structure (green, PDB ID: 5UA0). The inset depicts the proximity of the conserved ‘latch and catch’ residues of AtDHDPR1 (blue) and AtDHDPR2 (pink) allowing for hydrogen bonding, which stabilises the enzyme’s closed conformation.

Interestingly, the AtDHDPR1 structure lacked density for a nearly identical set of active site residues to those that could not be modelled in chain B of the AtDHDPR2 structure. The substrate binding loop within which these residues are contained is highly flexible, which may explain the disorder observed in this region. Indeed, such flexibility is supported by the inhibition of AtDHDPR1 by its substrate that was observed here, which has been reported to be a consequence of increased flexibility in this region in the plant enzymes.^17^ A structural alignment of AtDHDPR1 and AtDHDPR2 resulted in an RMSD of 0.5 Å over 1741 equivalent atoms, indicating a high degree of structural similarity (Figure 3).

### Sequence and structural homology of the 2,6-PDC binding pocket

The previously determined crystal structure of EcDHDPR in complex with 2,6-PDC (PDB ID: 1ARZ) showed a hydrogen bond network encompassing five active site residues that are conserved across bacterial species (Figure 4A).^25^ An alignment of the EcDHDPR amino acid sequence with that of both AtDHDPR isoforms revealed that four of the five residues involved in 2,6-PDC binding are conserved (Figure 4B). EcDHDPR and AtDHDPR1 resulted in an RMSD of 6.2 Å for 1193 equivalent carbon atoms, whereas an RMSD of 3.5 Å for 1276 equivalent carbon atoms was revealed when EcDHDPR was overlaid with AtDHDPR2. Inspection of the residues involved in 2,6-PDC binding indicates a structurally conserved binding pocket that may accommodate 2,6-PDC binding to AtDHDPR enzymes (Figure 4C).

**Figure 4.**
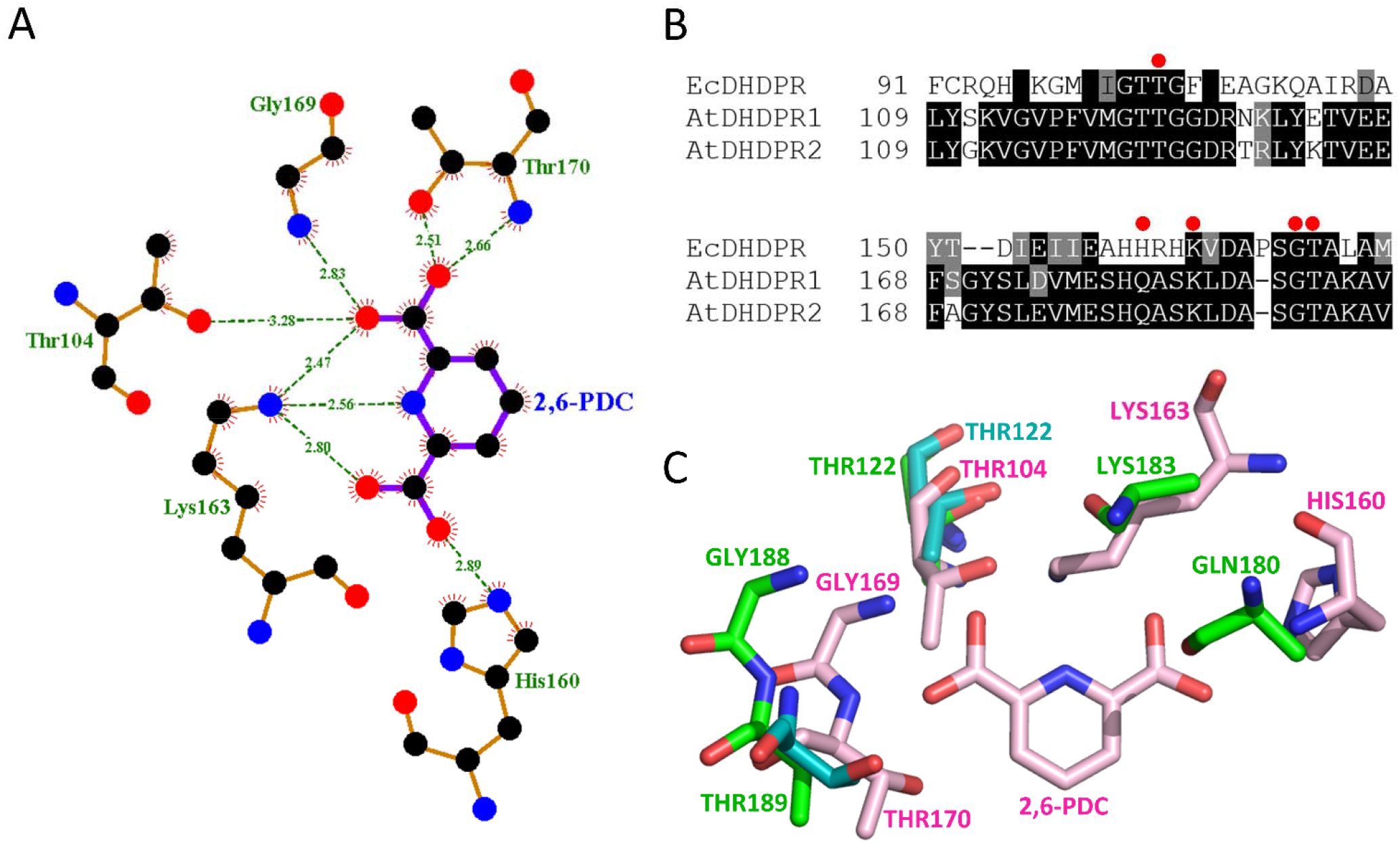
Homology of the 2,6-PDC binding pocket. (A) Schematic diagram of the 2,6-pyridine dicarboxylic acid (2,6-PDC) binding site in the three-dimensional structure of *Escherichia coli* DHDPR generated using LIGPLOT+. (B) Primary sequence alignment of DHDPR from *E. coli* (Ec) and *Arabidopsis thaliana* (At) generated using CLUSTALW. Residues forming hydrogen bonds with 2,6-PDC are indicated by red dots. (C) Overlay of the 2,6-PDC-bound EcDHDPR active site binding pocket (pink, PDB ID: 1ARZ) with that of AtDHDPR1 (cyan, PDB ID: 7T34) and AtDHDPR2 (green, PDB ID: 5UA0).

### Suitability of 2,6-PDC as a plant DHDPR inhibitor scaffold

The potency of 2,6-PDC against both AtDHDPR isoforms was assessed using the DHDPS-DHDPR coupled assay, with substrates fixed at their respective *K_M_* values. 2,6-PDC inhibited AtDHDPR1 with an *IC*_50_ of 140 μM and AtDHDPR2 with an *IC*_50_ of 470 μM. Therefore, 2,6-PDC represents the first example of a plant DHDPR inhibitor and a potential scaffold for the synthesis of more potent analogues. To assess whether 2,6-PDC had activity against plants, *A. thaliana* seeds were raised on media containing a concentration gradient of inhibitor (Figure 5A). The herbicidal activity of 2,6-PDC was visually evident from a near complete inhibition of growth above a concentration of 2 mM, and reduced growth at 1 mM. Quantitative analysis of the plant growth area at each concentration enabled the generation of a dose response curve from which an LD_50_ of 1.1 mM was determined (Figure 5B).^26^ Given that 2,6-PDC is an inhibitor of bacterial DHDPR enzymes, we employed antibacterial assays to examine its selectivity. The compound had no activity against bacterial strains commonly found in soil, with MIC values >5 mM (Supplementary Table S1). We further investigated the specificity of 2,6-PDC using a cell viability assay, which demonstrated that it lacks off-target toxicity against the human cell line, HepG2, with no significant difference in viability observed between control and treatment groups up to 5 mM (Supplementary Figure S4). The specificity for plants over common soil flora and human cells suggested that 2,6-PDC could be a suitable scaffold to pursue for the development of herbicidal DHDPR inhibitors.

**Figure 5.**
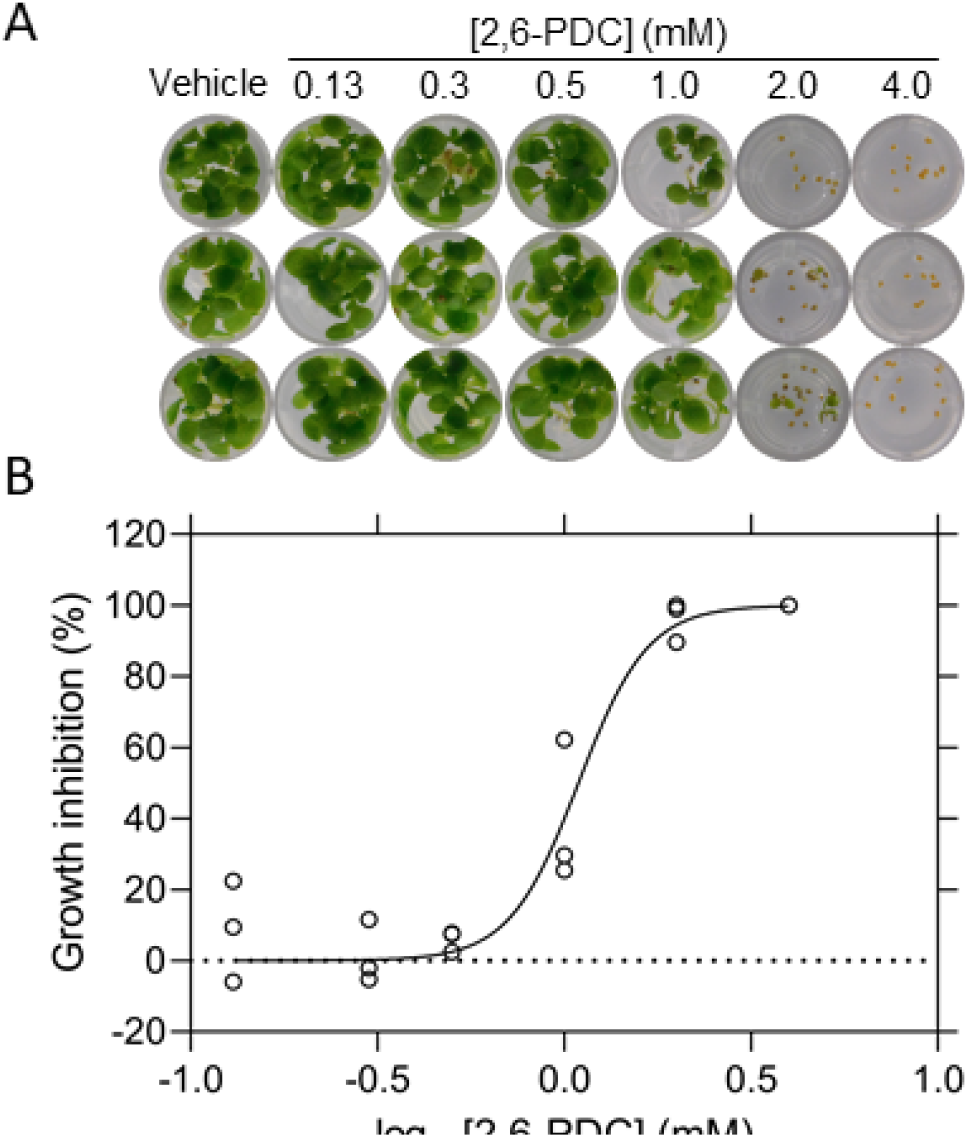
Growth of *A. thaliana* on media containing 2,6-PDC. (A) *A. thaliana* seeds raised on agar containing a concentration gradient of 2,6-PDC. (B) Dose-response curve for 2,6-PDC. The percentage of growth inhibition (○) relative to the DMSO control is plotted as a function of 2,6-PDC concentration. Data were fit to a variable slope model (—) resulting in an R^2^ value of 0.95. *n* = 3.

### Synthesis and structure-activity relationship of 2,6-PDC analogues

To afford insight into the chemical features important for 2,6-PDC potency, a series of analogues were synthesised. In total, 21 analogues were prepared, incorporating amide, ester and aldehyde functionality centred around a 2,6-disubstitued pyridine core (Supplementary Methods S1 and Supplementary Data S1). Analogues were screened for herbicidal activity at a concentration of 1 mM (~LD_50_ for parent) against *A. thaliana* seeds grown on agar (Table 2). The amide analogues (**1-7**) generally displayed reduced activity compared to 2,6-PDC. Of the linear chain esters (**8-13**), those with a carbon chain length of 3 and 4 (**10**, **11**) were more active than 2,6-PDC. Incorporation of a terminal halide (**14**, **15**) or alkynyl functionality (**16**, **17**) in the carbon chain also improved the activity, although was not as beneficial as the equivalent unsubstituted carbon chain length (**10**, **11**). However, modification of the shorter carbon chain length analogues (**8**, **9**) through the addition of the electron-withdrawing CF_3_ moiety (**18**) or a branched carbon chain (**19**) afforded improvements relative to analogues **8** and **9**. Conversely, the 3-methoxy substituted analogue **20** had slightly reduced activity compared to the unsubstituted equivalent (**11**). The benefits of modification of the carboxylic acid moiety to the corresponding aldehyde (**21**) were comparable to analogues **10** and **11**. Those exhibiting the best activity, that is, those that arrested growth upon radicle emergence or prevented seed germination entirely, had a carbon chain length of 2 if halide-substituted (**14**, **15**), or 3-4 if unsubstituted (**10**, **11**), with the exception of the aldehyde (**21**).

### Herbicidal efficacy of 2,6-PDC analogues

The most promising inhibitors identified from the agar assays described above were screened for herbicidal activity against soil-grown plants alongside 2,6-PDC (Figure 6). In contrast to the growth inhibition studies conducted on media (Table 2), the clear distinction between the effects of analogues on soil-grown *A. thaliana* allowed us to identify four lead compounds (**14**, **15**, **16**, **17**). The halide substituted analogues (**14**, **15**) largely prevented seed germination, and for those that did germinate, growth was greatly impaired (Figure 6). Interestingly, the terminal alkynyl functionality (**16**, **17**) was the most beneficial, more so than the saturated carbon chain of equivalent length (**10**, **11**) (Figure 6). Of the two alkynes, the shorter carbon chain (**16**) was more effective at preventing germination (Figure 6). To assess whether the herbicidal efficacy observed against *A. thaliana* could extend to other species, the invasive weed species *L. rigidum* was treated with **16** (Figure 7). Treatment with 1200 mg·L^-1^ (equivalent to 48 kg·ha^-1^) of **16** successfully inhibited germination and growth, albeit it with reduced potency than against *A. thaliana,* with treated plants having a significantly reduced fresh weight of shoots and roots compared to the vehicle control (Figure 7A-B). A significant reduction in shoot dry weight was also observed (Figure 7C). This compound represents the first example of a lysine biosynthesis pathway inhibitor with herbicidal activity against a weed species.

**Figure 6.**
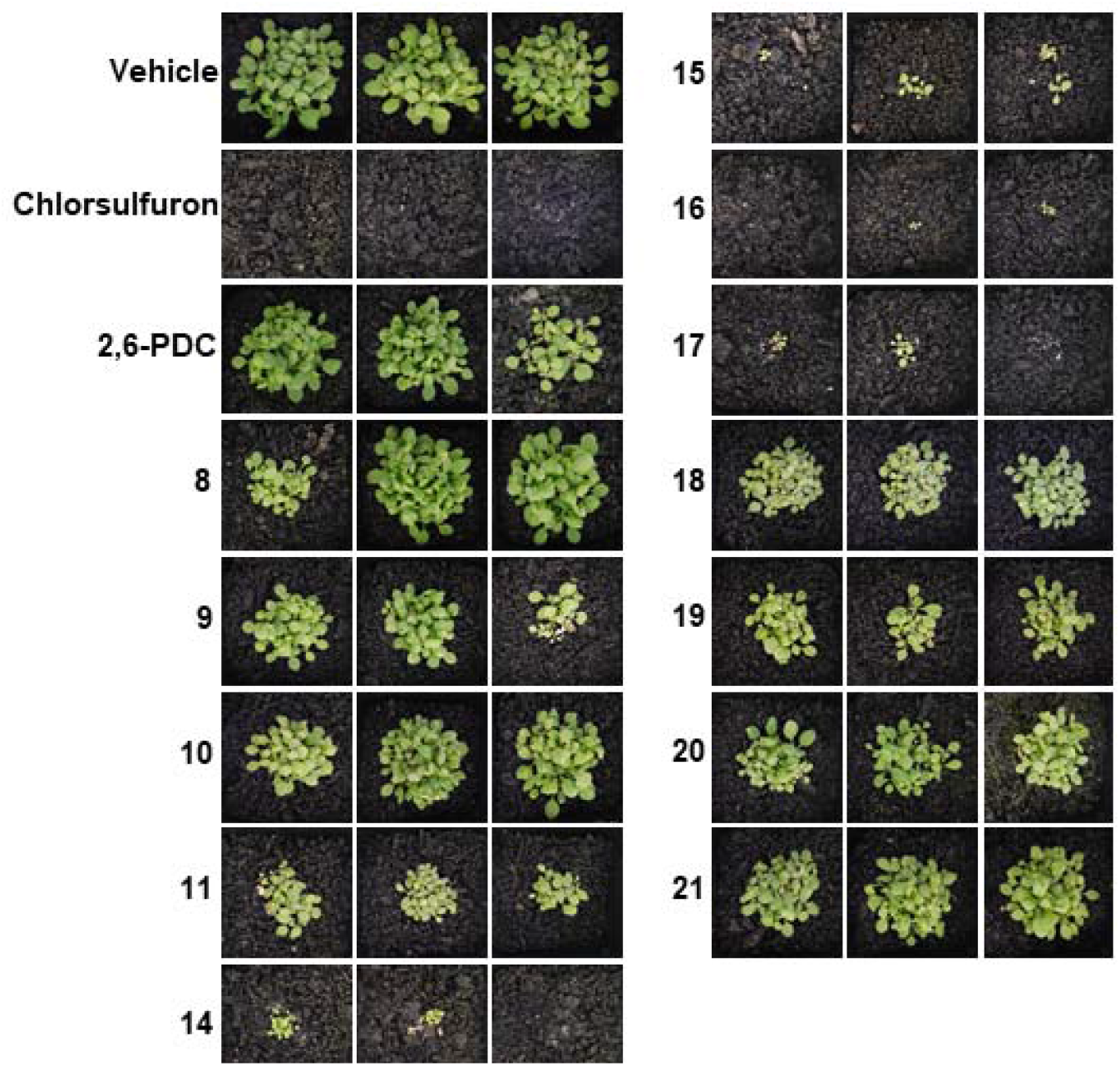
Herbicidal activity of 2,6-PDC analogues against *A. thaliana*. 14-day growth of *A. thaliana* treated with 1200 mg·L^-1^ of 2,6-PDC analogues. Images show the three replicates performed for each inhibitor.

**Figure 7.**
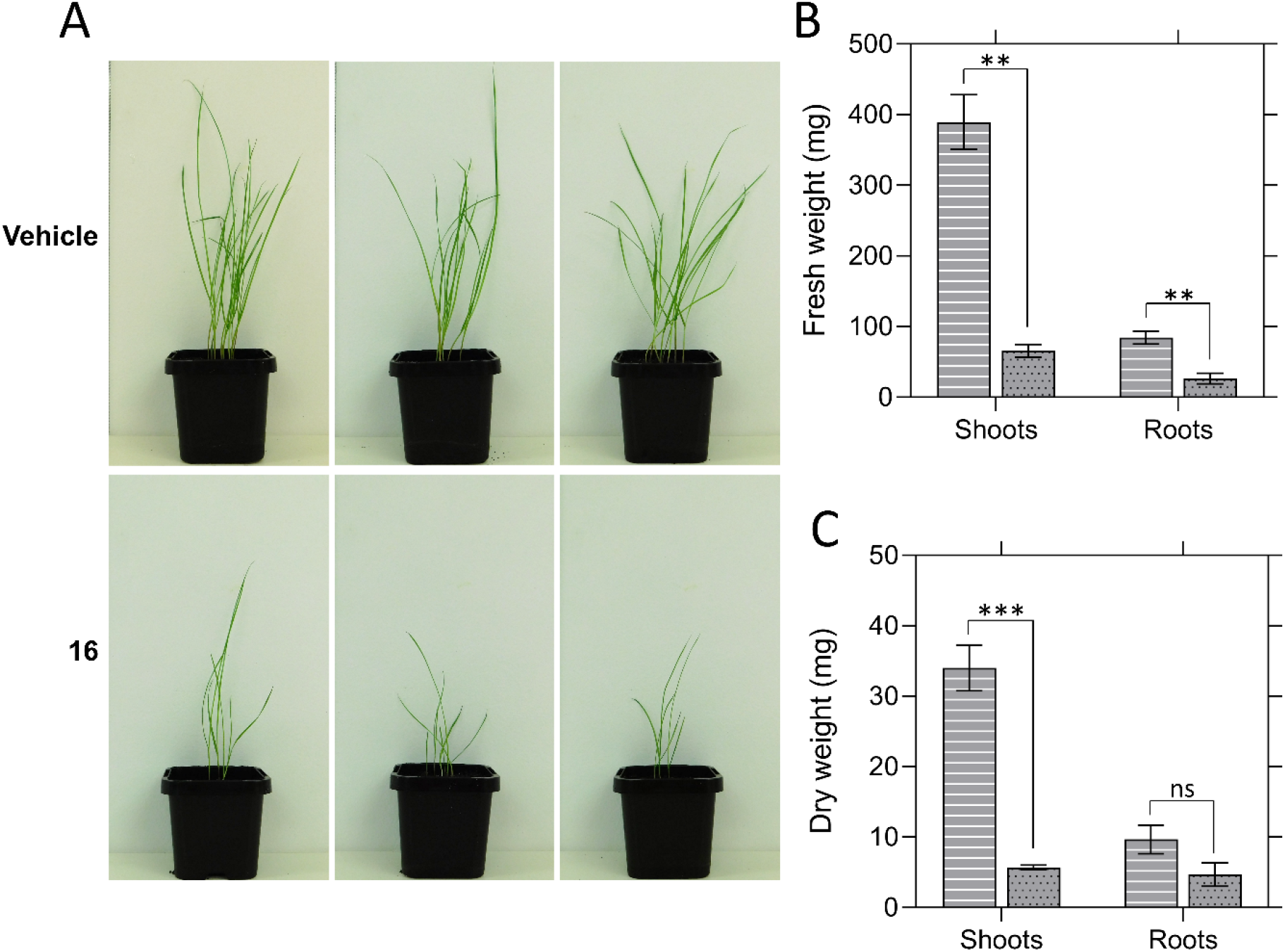
Growth of *Lolium rigidum* treated with 16. (A) 14-day growth of *L. rigidum* treated with three pre-emergence treatments of vehicle control (2% (v/v) DMSO, 0.01% Agral) or 1200 mg·L^-1^ of **16**. Images show the three replicates performed. (B) Fresh weight of *L. rigidum* shoots and roots following treatment of plants with vehicle control (lines) or 1200 mg·L^-1^ of **16** (dots). Shoots, *P* = 0.0013, roots, *P* = 0.0076, unpaired Student’s two-tailed *t*-test. (C) Dry weight of *L. rigidum* shoots and roots following treatment of plants with vehicle control (lines) or 1200 mg·L^-1^ of **16** (dots). Shoots, *P* = 0.0009, roots, *P* = 0.1295, unpaired Student’s two-tailed *t*-test. Data were normalised against the vehicle control. Data represents mean ± S.E.M. (*N* = 3). \*\**P* < 0.01, \*\*\**P* < 0.001.

## DISCUSSION

The inhibition of amino acid biosynthesis in plants has been a historically successful herbicide development strategy. However, examples of herbicidal lysine biosynthesis inhibitors had not been identified until our recent study reporting the first class of such inhibitors, which target the DHDPS enzyme.^7^ Subsequently, we set out to explore the next enzyme in the lysine biosynthesis pathway, DHDPR, as a potential herbicide target. Compared to amino acid biosynthesis enzymes targeted by commercial herbicides, the maximal expression levels of both AtDHDPR isoforms are comparable or lower, suggesting that achieving phytotoxicity with DHDPR inhibitors should not be hindered by high levels of target expression (Supplementary Table S2).^27^ Although no inhibitors of plant DHDPR enzymes have been reported previously, 2,6-PDC has been identified as an inhibitor of bacterial DHDPR orthologues, and thus, the present study aimed to assess the potential to repurpose the 2,6-PDC scaffold as a herbicide candidate.

Comparison of the primary sequences and crystal structures of bacterial and plant DHDPR orthologues revealed a high degree of conservation at the 2,6-PDC binding site, suggesting that this compound may also inhibit plant enzymes. Enzyme inhibition and plant germination assays supported our hypothesis. The presence of DHDPR in bacteria means that potential disruption of beneficial soil microbe communities needs to be addressed in the design of herbicidal DHDPR inhibitors. Despite 2,6-PDC being an *in vitro* inhibitor of bacterial DHDPR, the lack of antibacterial activity suggests that plant-specific inhibitors can be developed. Indeed, efflux and poor uptake of compounds by bacteria, which can impede the development of antibacterial agents, may conversely be an advantage in repurposing them as specific herbicides.^28–30^ Our findings that 2,6-PDC inhibited the germination of plants with specificity over bacterial and human cells prompted the subsequent synthesis of 21 analogues of 2,6-PDC, some of which had improved potency in a plant germination screen. Subsequent in-soil testing revealed that some of the analogues, which appeared promising in the screening phase, had reduced activity against soil-grown *A. thaliana*. This may be attributed to a complex range of factors influencing herbicidal activity. For example, compounds with good activity against the enzyme target may not necessarily have good soil binding properties, or resistance to clearance by the plant or soil flora. Whilst testing on plants in soil is important to assess these factors, initial screening of compounds in media provides an efficient strategy to rule out compounds lacking activity. Moreover, four of the analogues almost completely inhibited *A. thaliana* germination on soil, at a dose within one order of magnitude of conventional application rates of commercial herbicides such as asulam and atrazine.

Treatment of the invasive species *L. rigidum* with the most promising inhibitor **16** resulted in significant inhibition of its growth, suggesting that DHDPR inhibitors have the potential to be developed into herbicide candidates. However, the reduced potency of **16** against *L. rigidum* compared to *A. thaliana* instantiates the long road from lead identification to commercial formulation. Optimisation of the physicochemical properties of herbicide leads has the potential to improve soil persistence, delivery into the plant and leaf uptake for potential post-emergence application (Supplementary Table S3). For example, increasing the lipophilicity of these compounds would likely increase their translocation across the cuticle, cell wall and cell membrane. Furthermore, the high degree of conservation of DHDPR enzymes across plant species suggests that the specificity of these compounds for weeds is unlikely (Supplementary Figure S5).^31^ Directed evolution experiments would therefore be of interest to identify mutations which may be used to engineer crops resistant to DHDPR active site inhibitors. Identifying such mutations would also facilitate the monitoring of weed populations for the emergence of resistance so that early intervention strategies may be implemented.

Repurposing inhibitor scaffolds, as we have exemplified here, has the potential to fast-track herbicide discovery given that lead identification often involves costly high-throughput screening, or time-consuming rational design.^32^ Indeed, drug repurposing efforts have recently uncovered the herbicidal efficacy of the antibiotic ciprofloxacin, as well as antimalarial lead compounds.^33,34^ However, these drugs could not be used as herbicides due to the risk of accelerating the development of resistance to important medicines, without modifications to improve plant specificity. This drawback may be overcome by repurposing scaffolds that have not progressed through the drug development pipeline, such as 2,6-PDC. This study paves the way for future research into repurposing scaffolds previously identified as inhibitors of bacterial targets that have a high degree of similarity to enzymes in the plant kingdom. Given the rapidly increasing rate of herbicide resistance, such scaffolds could represent novel molecules for the development of much-needed new herbicide modes of action.

## METHODS

### Protein expression and purification

Synthetic codon-optimised genes encoding AtDHDPR1 (At2G44040) and AtDHDPR2 (At3G59890), excluding the chloroplast transit peptides, cloned into the pET11a expression vector were purchased from Bioneer (Daejeon, South Korea). Plasmids were transformed into *E. coli* BL21 (DE3) cells, which were subsequently treated with 1.0 mM IPTG to induce recombinant protein overexpression and cultured at 25 □ for 18 h. Cells were harvested by centrifugation, resuspended in lysis buffer (20 mM Tris-HCl, 20 mM imidazole, 500 mM NaCl, pH 8.0) and lysed by sonication. Following cell debris removal by centrifugation, soluble protein was applied to a His-Trap column and eluted over a stepwise gradient of imidazole (0-500 mM).^35^ Human rhinovirus 3C protease and 0.5 mM TCEP were added to protein-containing fractions and incubated at room temperature for 1 h for fusion tag cleavage. The protein mixture was applied to a His-Trap column for removal of the protease and the cleaved tag before dialysing into storage buffer (20 mM Tris-HCl, 150 mM NaCl, pH 8.0) and adding 0.5 mM TCEP.

### Circular dichroism spectroscopy

Spectra were collected using a CD spectrometer Model 420 (Aviv Biomedical) as described previously.^36,37^ AtDHDPR proteins in 20 mM NaH_2_PO_4_, 50 mM KF, pH 8.0 were diluted to 0.15 mg·mL^-1^. Wavelength scans were performed between 195 and 260 nm with a slit band width of 1.0 nm, step size of 0.5 nm and 5.0 s signal averaging time in a 1.0 mm quartz cuvette. The CDPro software package was used to fit the data to the SP22X reference set.^38^

### Enzyme kinetics and inhibition assays

The DHDPS-DHDPR coupled assay was used to measure DHDPR enzyme activity, using methods similar to those described previously.^22,23^ Briefly, reaction mixtures were incubated at 30 °C for 12 min before a second 60 s incubation following the addition of excess *E. coli* DHDPS (51 μg·mL^-1^) for generation of the DHDP substrate. Assays were then initiated by the addition of the relevant DHDPR isoform (2.6 μg·mL^-1^), and substrate turnover measured spectrophotometrically at 340 nm via the associated oxidation of the cofactor NADPH. For determination of kinetic parameters, data were fit to a substrate inhibition model (Equation 1).

For determination of *IC*_50_ values, DHDPR activity was measured in the presence of titrated concentrations of inhibitors in 1% (v/v) DMSO and data were fit to a variable slope model (Equation 2). Experiments were performed in technical triplicates.

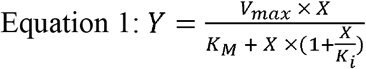

Where *Y* = initial rate, *V*_max_ = maximal enzyme velocity, *X* = concentration of substrate, *K*_M_ = Michaelis-Menten constant, *K*_i_ = dissociation constant for substrate binding.

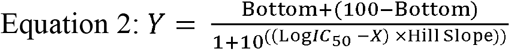

Where *Y* = response, Bottom = plateau in the same units as *Y*, Hill slope = slope factor.

### Crystallisation and structure determination

Protein crystallisation screening for AtDHDPR1 was initially performed at the CSIRO Collaborative Crystallisation Centre (CSIRO, Parkville, Melbourne, Australia) using the sitting drop vapour diffusion method and Shotgun crystal screen at 8 °C and 20 °C. Conditions were optimised in-house using the hanging drop vapour diffusion method and 4 μL drops comprised of 2 μL protein solution (10 mg·mL^-1^ AtDHDPR1, 2 mM NADPH) and 2 μL reservoir solution. Crystals used for data collection were obtained after 2 days at 20 °C using reservoir solutions containing 0.1 M bis-tris hydrochloride (pH 6.5), 0.245 M magnesium formate, 22% (w/v) PEG 3350. Crystals were transferred to cryo-protectant (0.1 M bis-tris hydrochloride (pH 6.5), 0.245 M magnesium formate, 22% (w/v) PEG 3350, 24% (v/v) glycerol) and flash-frozen in liquid nitrogen. X-ray diffraction data were collected on the MX2 beamline at the Australian Synchrotron.^39^ Data were processed using XDS and scaled using AIMLESS and structure solved by molecular replacement using Auto-Rickshaw employing the EcDHDPR structure (PDB ID: 1ARZ) as a search model.^40–43^ Model refinement and building was conducted in PHENIX and COOT respectively.^44,45^ Model quality was evaluated using MOLPROBITY.^46^ The structure has been deposited in the Protein Data Bank with code 7T34. Data collection and refinement statistics are presented in Table 1.

**Table 1.**
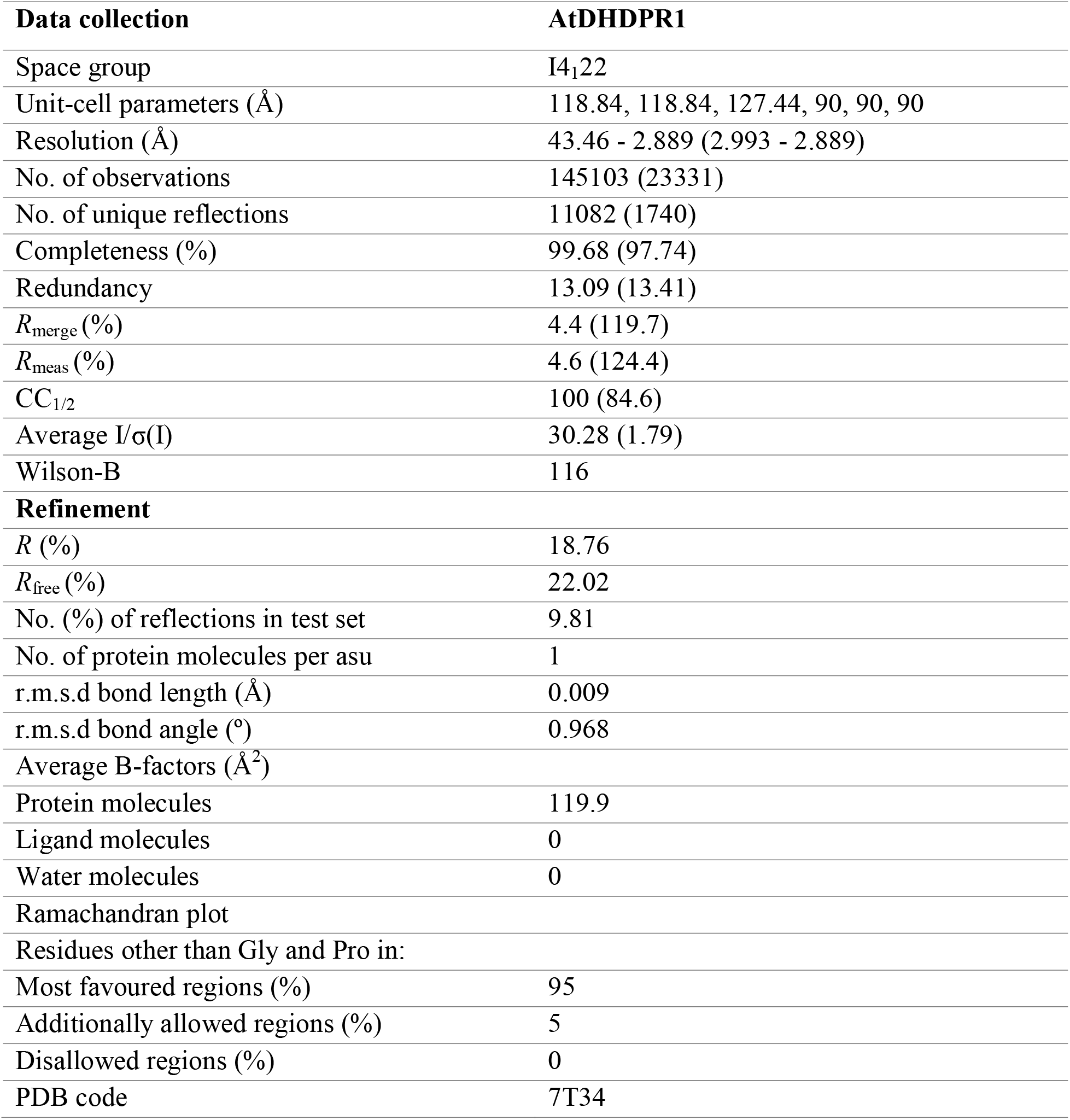
Summary of AtDHDPR1 crystallographic data collection, processing and refinement statistics.

**Table 2.**
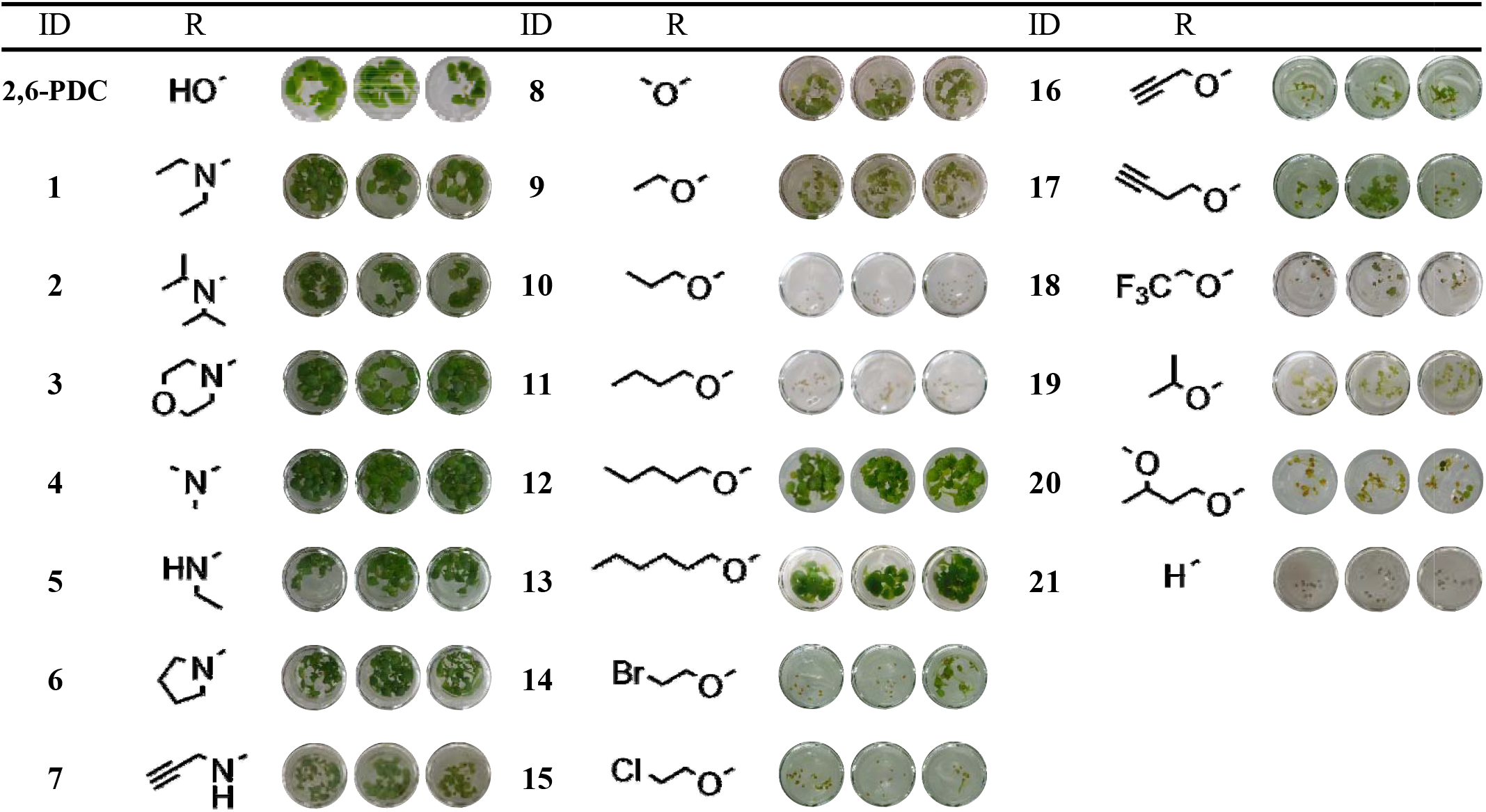
Growth of *A. thaliana* on media containing 2,6-PDC analogues at a concentration of 1 mM performed in triplicate.

### Analytical ultracentrifugation

Sedimentation velocity experiments were performed in a Beckman Coulter XL-A analytical ultracentrifuge at 25 °C using methods similar to those described previously.^47–50^ Briefly, 380 μL of protein storage buffer containing 0.5 mM TCEP, and 400 μL of protein at 0.9 mg·mL^-1^ were loaded into double sector cells with synthetic quartz windows. Centrifugation of cells was performed at 40,000 rpm using a 4-hole An50-Ti rotor. Data were collected continuously without averaging at 280 nm with a radial step size of 0.003 cm. SEDNTERP software was used to compute solvent density (1.007 g·mL^-1^), solvent viscosity (0.010259 cp) and estimated protein partial specific volume (0.738218 mL·g^-1^).^51^ SEDFIT was used to fit absorbance as a function of radial position to the Lamm equation to determine the continuous sedimentation coefficient distribution.^51,52^

### Growth inhibition assays on media

*A. thaliana* ecotype Columbia (Col-0) seeds were surface sterilised for 5 min in 80% (v/v) EtOH, followed by 15 min in 1% (v/v) NaClO and then thorough washing in sterile H_2_O. Seeds were resuspended in sterile 0.1% (w/v) plant tissue culture grade agar before stratification at 4 □ in the dark for 3 days. Seeds were sown on 0.25 mL of growth medium (0.8% (w/v) agar, 1% (w/v) sucrose, 0.44% (w/v) Murashige & Skoog salts with vitamins, 2.5 mM 2-(.*N*-morpholino)-ethanesulfonic acid (MES), pH 5.7) containing either DMSO (vehicle control) or inhibitor in 96-well microplates, which were then sealed with porous tape. Plates were transferred to a chamber at 22 □ under a 16 h light (100 μmol m^-2^ s^-1^)/8 h dark schedule for 7 days before photos were taken. Quantification of *A. thaliana* growth inhibition was performed as described previously, and the data were fit to a variable slope model (Equation 2) to determine the LD50.^26^ Experiments were performed in technical triplicates.

### Antibacterial assays

Minimum inhibitory concentration (MIC) values were determined using a broth microdilution assay in accordance with the guidelines issued by the Clinical Laboratory Standard Institute.^53,54^ Serial dilutions of 2,6-PDC were prepared in 96-well microplates using tryptic soy broth as the diluent.^55,56^ Plates were inoculated with 1 × 10^5^ colony forming units per mL of bacteria and incubated at 25 □ for 20 h. Growth was assessed by measuring the absorbance at 600 nm and the lowest concentration of inhibitor with no observable growth determined to be the MIC value.^55,56^ Experiments were performed in biological triplicates.

### Cell culture and cytotoxicity assays

The human hepatocellular carcinoma (HepG2) cell line was grown and maintained in a humidified incubator at 37 □ with 5% CO_2_ in high (HepG2) glucose Dulbecco’s Modified Eagle’s Medium (DMEM, Gibco, Waltham, USA, 11885084) with 10% (v/v) fetal bovine serum (FBS, Gibco, 10099141) and 50 U/mL penicillin/50 μg·mL^-1^ streptomycin. Cytotoxicity assays were performed using similar methods to those previously described.^7,57^ Specifically, 5,000 viable cells/well were seeded into 96-well plates and incubated at 37 □ for 24 h. Cells were subsequently treated with varying concentrations of inhibitor, 1% (v/v) DMSO or the cytotoxic defensin protein NaD1 at 30 μM and incubated at 37 □.^57,58^ After 48 h, cells were incubated with 0.5 mg·mL^-1^ [3-(4,5-dimethylthiazolyl)-2,5-diphenyl-tetrazolium bromide] in DMEM without FBS at 37 □ for 3 h. All liquid was removed from wells and formazan crystals dissolved in DMSO before measuring the absorbance at 570 nm. The percentage viability remaining reported is relative to the 1% (v/v) DMSO vehicle control. Four technical replicates were performed for each treatment condition.

### Herbicidal activity analyses

The herbicidal efficacy of AtDHDPR inhibitors in soil was assessed using methods similar to those reported previously.^34,59^ Pre-wet seed-raising soil (pH 5.5) (Biogro, Dandenong South, VIC, Australia) supplemented with 0.22% (w/w) Nutricote N12 Micro 140 day-controlled release fertiliser (Yates, Sydney, NSW, Australia) was used for all experiments. For experiments conducted with *A. thaliana,* approximately 40 ecotype Columbia (Col-0) seeds were sown in pots onto the soil surface following surface sterilisation and stratification as described for germination assays on media. For experiments conducted with *L. rigidum,* 10 seeds were sown at a depth of 0.5 cm into pots of pre-wet soil, following stratification at 4 □ for 21 days in the dark. Compounds dissolved in DMSO were diluted to working concentrations in H_2_O containing 0.01% (v/v) Agral (Syngenta, North Ryde, NSW, Australia) to a final DMSO concentration of 2% (v/v). Treatments were given by pipetting 1.0 mL (*A. thaliana)* or 2.0 mL (*L. rigidum)* of test compound, vehicle control or positive control (chlorosulfuron PESTANAL (Sigma-Aldrich, North Ryde, NSW, Australia)) directly onto seeds upon sowing and on each of the subsequent two days. Plants were grown in a chamber at 22 □ under a 16 h light (100 μmol m^-2^ s^-1^)/8 h dark schedule for 14 days before photos were taken. Roots and shoots were separated prior to drying at 70 □ for 72 h. Experiments were performed in biological triplicates.

## Supplementary Information

Supplementary Figure S1. Expression and purification of recombinant AtDHDPR enzymes.

Supplementary Figure S2. Secondary structure of AtDHDPR isoforms.

Supplementary Figure S3. Sedimentation velocity analysis by analytical ultracentrifugation of AtDHDPR1.

Supplementary Figure S4. Viability of human cells treated with 2,6-PDC.

Supplementary Figure S5. Sequence alignment of plant DHDPR enzymes.

Supplementary Table S1. Minimum inhibitory concentration (MIC) values of 2,6-PDC against soil bacteria.

Supplementary Table S2. Maximal expression levels of *A. thaliana* DHDPR isoforms and commercial herbicide targets determined by RNA-sequencing.

Supplementary Table S3. Physicochemical properties of lead compounds.

Supplementary Methods S1. Synthesis of compounds.

Supplementary Data S1. NMR spectra of compounds.

## ACKNOWLEDGEMENTS

T.P.S.C. acknowledges the Australian Research Council for funding support through a DECRA Fellowship (DE190100806). Work in A.R.G.’s laboratory is supported by the Australian Research Council Research Hub for Medicinal Agriculture (IH180100006). E.R.R.M. acknowledges the Grains Research and Development Corporation (9176977) for support through a PhD scholarship and operational funding and La Trobe University for support through a Research Training Program scholarship. We thank Professor Ashley Franks (La Trobe University, Australia) for supplying bacterial isolates and Professor John Moses (La Trobe University, Australia) for providing infrastructure. We acknowledge the La Trobe University Comprehensive Proteomics Platform for providing infrastructure support. We acknowledge the use of the MX2 beamline at the Australian Synchrotron, part of ANSTO and employed the Australian Cancer Research Foundation (ACRF) detector.

## AUTHOR CONTRIBUTIONS

E.R.R.M. performed the biology experiments, analysed data and co-wrote the manuscript. A.S.B. synthesised compounds, analysed data and co-wrote the manuscript. M.-C.G. synthesised compounds and analysed data. M.D.H. and A.R.G. provided reagents and materials and revised the manuscript. S.P. analysed data and revised the manuscript. T.P.S.C. designed the research and co-wrote the manuscript.

## COMPETING INTERESTS

The authors declare no competing interests.

## Notes

### Competing Interest Statement

The authors have declared no competing interest.

