## Supplementary Information for "From bacteria to plants: a repurposing strategy in the pursuit for novel herbicides"

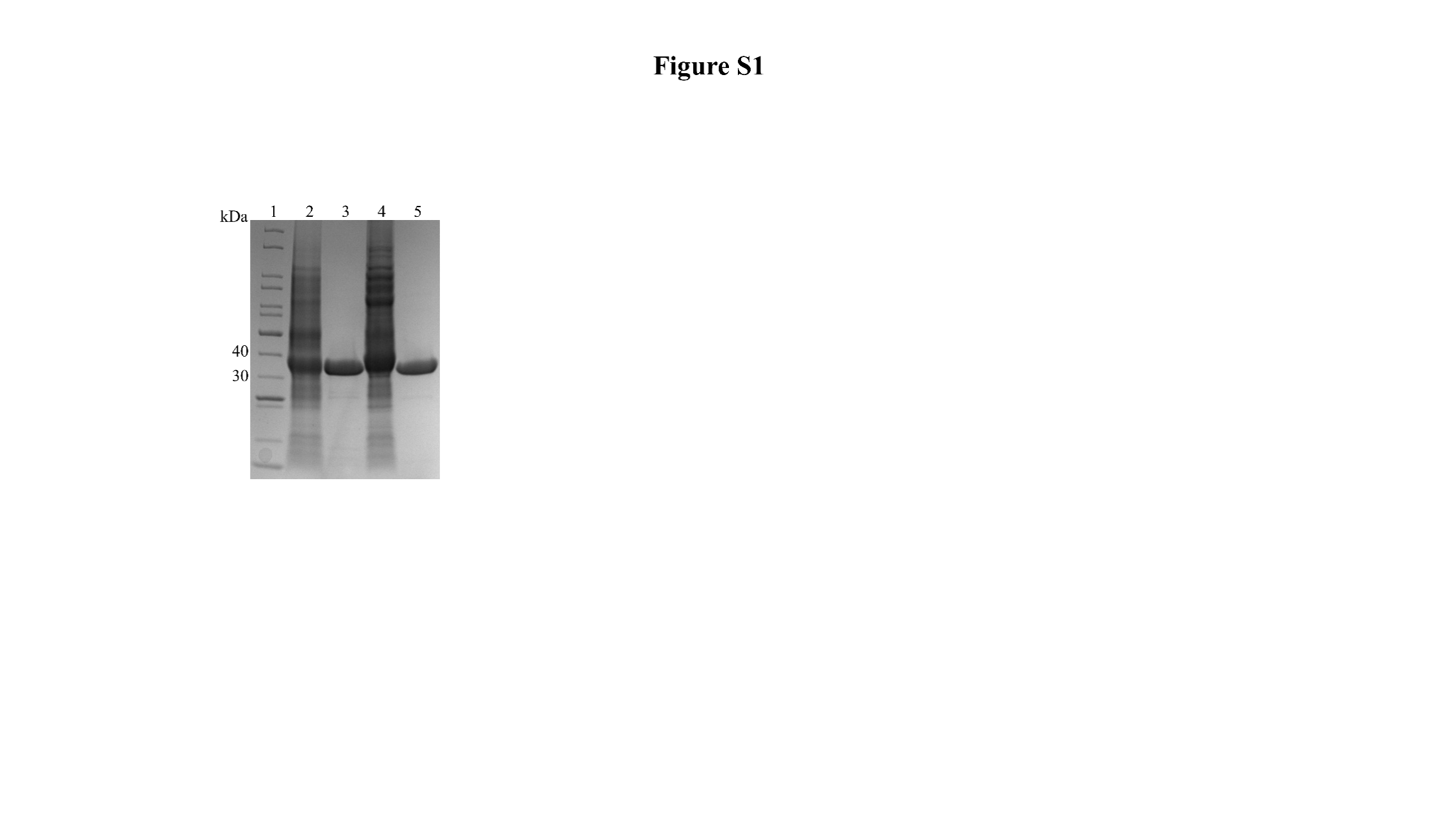
**SUPPLEMENTARY FIGURES**

**Supplementary Figure S1. Expression and purification of recombinant AtDHDPR enzymes.** Lane 1: molecular weight markers (kDa); lane 2: soluble extract of *E. coli* cultures expressing the AtDHDPR1 construct; lane 3: purified recombinant AtDHDPR1; lane 4: soluble extract of *E. coli* cultures expressing the AtDHDPR2 construct; lane 5: purified recombinant AtDHDPR2.

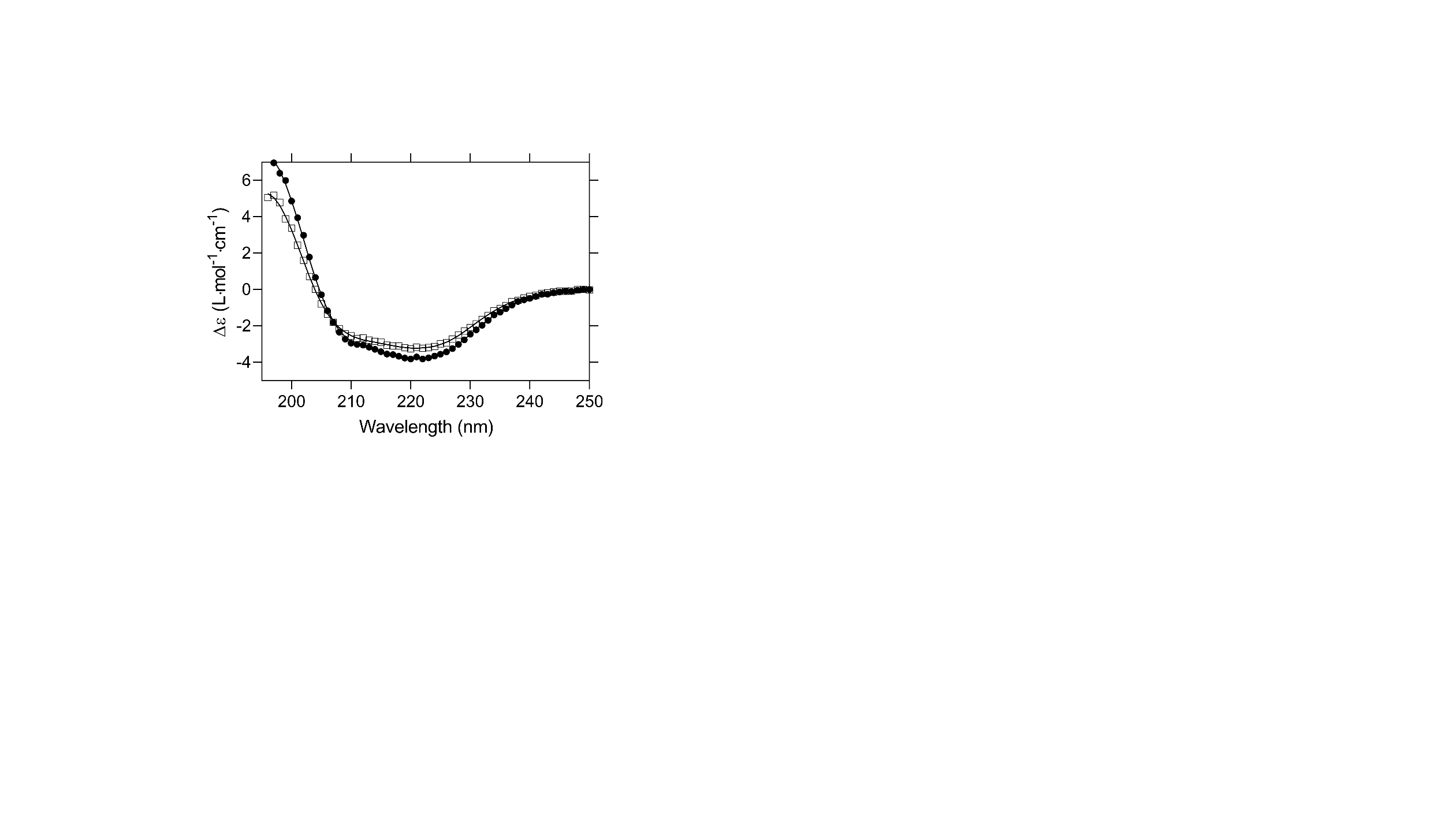

**Supplementary Figure S2. Secondary structure of AtDHDPR isoforms.** Spectra were collected at a protein concentration of 0.2 mg·mL^-1^ over wavelengths spanning 195-250 nm with a step size of 1.0 nm. The CONTINLL algorithm from the CDPro software package was used to fit the experimental data for AtDHDPR1 (□) and AtDHDPR2 (●) to the SP22X reference set (—). The fit predicted AtDHDPR1 to be comprised of 30% α-helix, 21% β-strand, 12% turn and 37% unordered with a RMSD of 0.061, and AtDHDPR2 to be comprised of 37% α-helix, 22% β-strand, 8% turn and 33% unordered with a RMSD of 0.067.

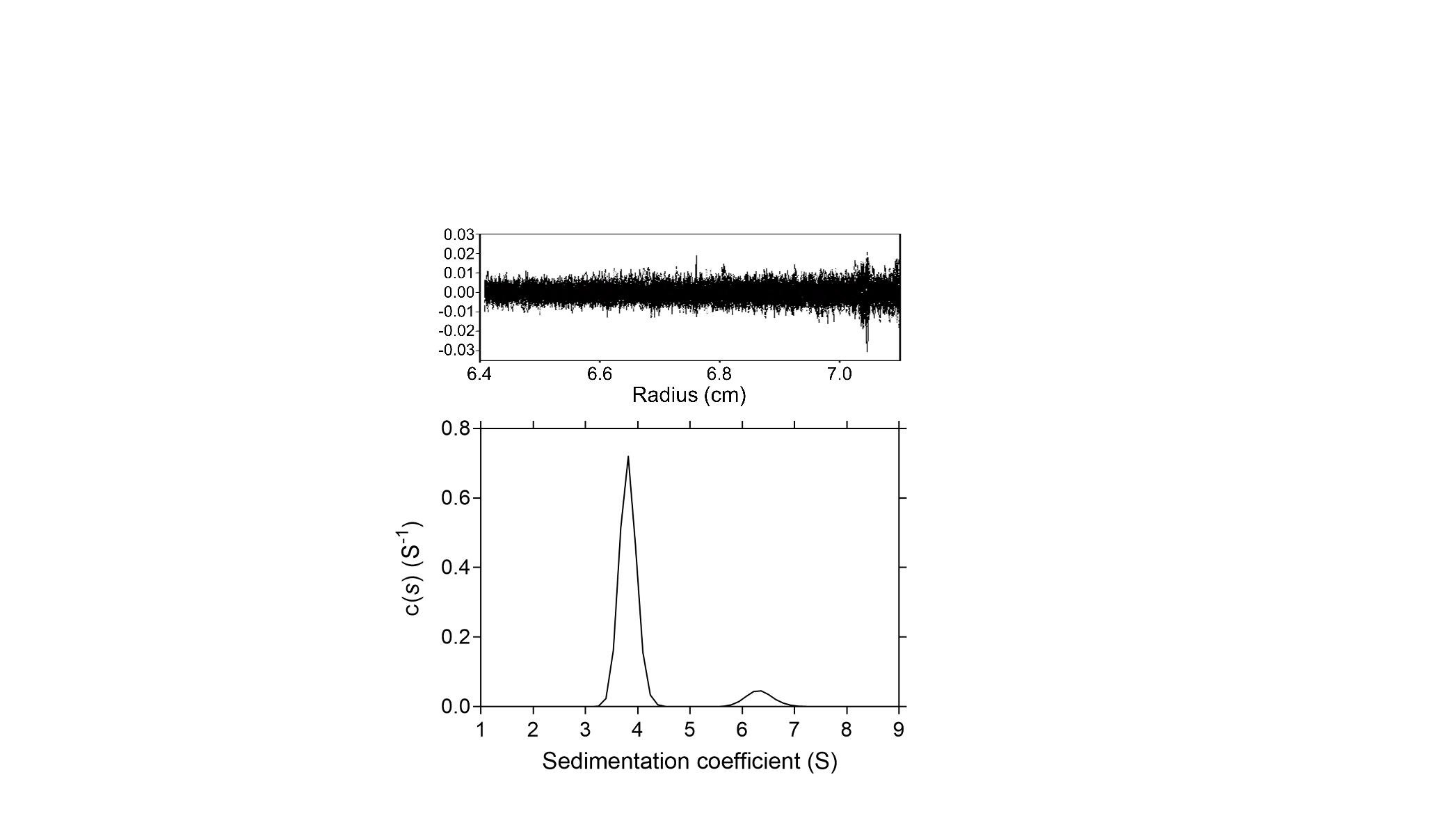

**Supplementary Figure S3. Sedimentation velocity analysis by analytical ultracentrifugation of AtDHDPR1.** Continuous sedimentation coefficient distribution analysis of AtDHDPR1 at a concentration of 0.9 mg·mL^-1^ resulted in peaks at ~4 S and ~6.5 S, which are consistent with a dimeric species and tetrameric species, respectively. Residuals resulting from nonlinear regression best fits are shown in the top panel.

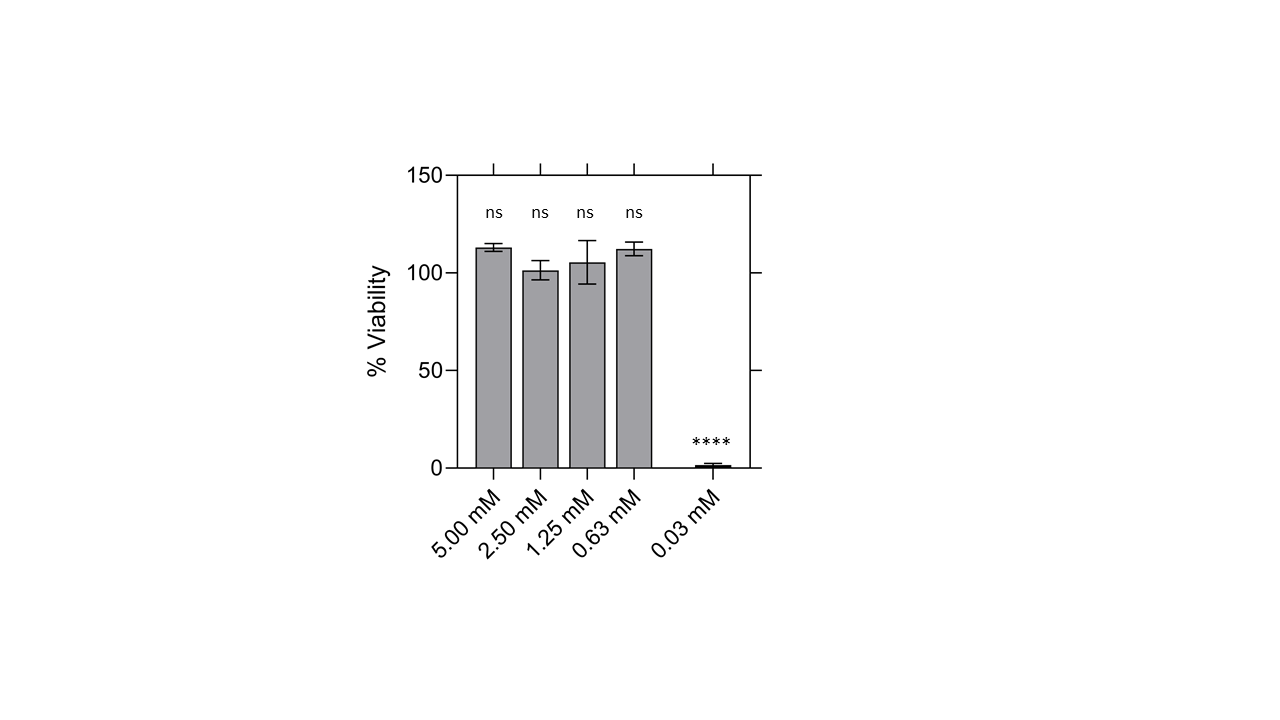

**Supplementary Figure S4. Viability of human cells treated with 2,6-PDC.** Percentage viability of the HepG2 human cell line after treatment with varying concentrations of 2,6-PDC (grey) or a concentration of the positive control defensin (black) determined using the MTT assay. Data were normalised against a vehicle control (1% (v/v) DMSO). Data represents mean ± S.E.M. (*n* = 4). One-way ANOVA multiple comparisons test, ns = not significant, *****P* < 0.0001.

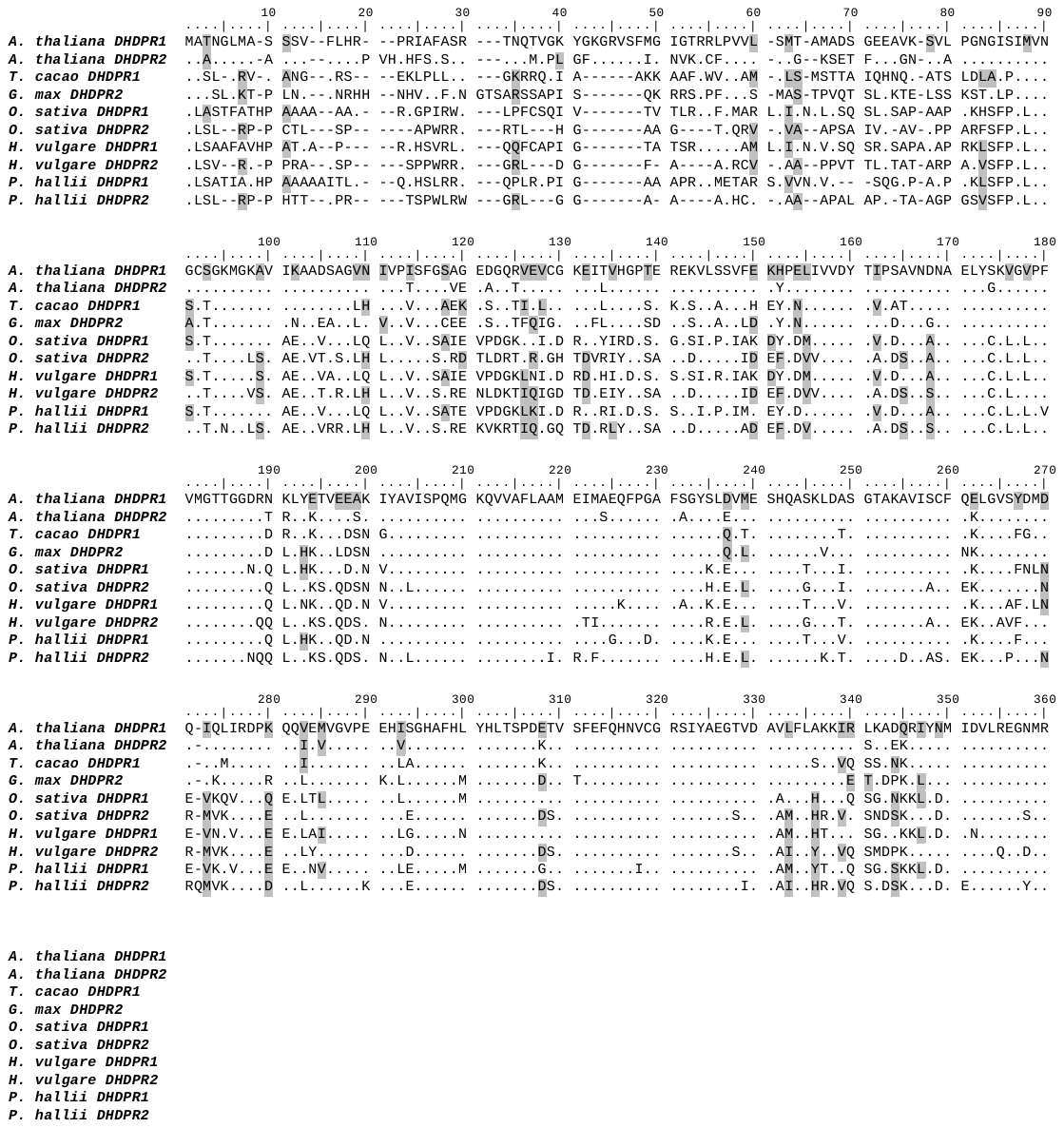

**Supplementary Figure S5. Sequence alignment of plant DHDPR enzymes.** *A. thaliana* DHDPR1 (UniProt ID: O80574) and DHDPR2 (UniProt ID: Q8LB01), *Theobroma cacao* DHDPR1 (UniProt ID: A0A061DK14), *Glycine max* DHDPR2 (UniProt ID: D2DKE9), *Oryza sativa* DHDPR1 (UniProt ID: Q67W29) and DHDPR2 (UniProt ID: Q10P67), *Hordeum vulgare* DHDPR1 (UniProt ID: F2D3R8) and DHDPR2 (UniProt ID: F2DWQ1) and *Panicum hallii* DHDPR1 (UniProt ID: A0A2T7F6F2) and DHDPR2 (UniProt ID: A0A2T7CF59). Residues are numbered in reference to AtDHDPR1. Identical residues are shown as dots (•), similar (≥50%) residues are shaded in grey, gaps are shown as dashes (-). Sequences were aligned using T-Coffee and edited using BioEdit (v 7.0.5.3).

**SUPPLEMENTARY TABLES**

**Supplementary Table S1. Minimum inhibitory concentration (MIC) values of 2,6-PDC against soil bacteria.**

| **Species** | **MIC (mM)** |
| --- | --- |
| *Enterobacter ludwigii* | >5.0 |
| *Cedecea davisae* | >5.0 |
| *Enterobacter cancerogenus* | >5.0 |

**Supplementary Table S2. Maximal expression levels of *A. thaliana* DHDPR isoforms and commercial herbicide targets determined by RNA-sequencing.**

| **Target** | **Commercial Herbicide Mode of Action** | **Maximum reads per gene** |
| --- | --- | --- |
| Dihydrodipicolinate reductase 1  *At2G44040.1* | - | 45 |
| Dihydrodipicolinate reductase 2  *At3G59890.1* | - | 18 |
| Acetolactate synthase  *At3G48560* | Branched chain amino acid biosynthesis inhibition | 251 |
| 5-enolpyruvylshikimate-3-phosphate synthase  *At1G48860* | Aromatic amino acid biosynthesis inhibition | 40 |
| 4-hydroxyphenylpyruvate dioxygenase  *At1G06570.1* | Tyrosine catabolism inhibition | 146 |

|  | **14** | **15** | **16** | **17** |
| --- | --- | --- | --- | --- |
| Molecular mass (g·mol^-1^) | 381.0 | 292.1 | 243.2 | 271.3 |
| clogP | 1.453 | 1.174 | 0.190 | 1.148 |
| clogS | -3.413 | -2.898 | -1.972 | -2.256 |
| tPSA | 64.96 | 64.96 | 64.96 | 64.96 |
| H-bond acceptors | 3 | 3 | 3 | 3 |
| H-bond donors | 0 | 0 | 0 | 0 |
| Rotatable bonds | 8 | 8 | 8 | 10 |

**Supplementary Table S3. Physicochemical properties of lead c**

**Supplementary Methods S1. Synthesis of compounds.**

**General Methods**

Commercial solvents and reagents were used as supplied. The petroleum ether used refers to the fraction with 40-60 °C boiling point. Unless otherwise stated, all reactions were monitored by TLC on Polygram® SIL/G25 plates and visualized using UV light (254 nm). ^1^H, ^13^C and ^19^F NMR spectra were recorded on either a Bruker Ascend™ 400 (400 MHz) or a Ultrashield™ 500 PLUS (500 MHz) instrument as dilute solutions in the deuterated solvent. All chemical shifts (δ) are reported in parts per million (ppm) with ^1^H and ^13^C NMR referenced to solvent signals [^1^H NMR: CDCl_3_ (7.27); ^13^C NMR: CDCl_3_ (77.16)]. Coupling constants (*J*) are reported in Hertz (Hz) and recorded after averaging. The multiplicity of the ^1^H NMR signals are designated by one of the following abbreviations: s=singlet, d=doublet, t=triplet, q=quartet, hept=heptet, m=multiplet, br=broad signal. Infra-red spectra as solutions in CHCl_3_ or with KBr discs, with the peaks recorded as ν_max_ (cm^-1^). HRMS were obtained using an Agilent 6530 accurate-mass Q-TOF LC/MS in electrospray ionisation (ESI) mode. Melting point data was collected using a Gallenkamp melting point apparatus.

**Synthesis of amides**

**General Procedure**

To a solution of pyridine-2,6-dicarbonyl dichloride (51.0 mg, 250 µmol, 1.00 eq) in CH_2_Cl_2_ (4 mL) was added the required amine (4.00 eq) at 0 °C. The solution was warmed room temperature and stirred for 16 h then diluted with CH_2_Cl_2_ (10 mL) and washed with brine (10 mL). The organic phase was dried over anhydrous MgSO_4_, filtered and concentrated under reduced pressure. The crude product was purified by silica gel column chromatography to yield the product.

***N^2^,N^2^,N^6^,N^6^*-tetraethylpyridine-2,6-dicarboxamide**^1^ **(1)**

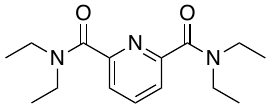

Following general procedure, the crude product was purified by silica gel column chromatography (20% EtOAc in petrol) to yield the title compound (**1**) as yellow solid (42.0 mg, 61%). m.p. 74 °C (lit: 82 °C).

**^1^H NMR** (500 MHz, CDCl_3_) δ 7.86 (t, *J* = 7.8 Hz, 1H), 7.61 (d, *J* = 7.8 Hz, 2H), 3.55 (q, *J* = 7.1 Hz, 4H), 3.33 (q, *J* = 7.1 Hz, 4H), 1.25 (t, *J* = 7.1 Hz, 6H), 1.13 (t, *J* = 7.1 Hz, 6H); **^13^C NMR** (126 MHz, CDCl_3_) δ 168.2, 153.7, 138.0, 123.7, 43.4, 40.3, 14.3, 12.9; **IR** ν_max_ (cm^-1^): 2988, 1628, 1485, 1217, 752; **HRMS** (ESI) calculated for C_15_H_24_N_3_O_2_ [M+H]^+^ 278.1863, found 278.1864.

***N^2^,N^2^,N^6^,N^6^*-tetraisopropylpyridine-2,6-dicarboxamide (2)**

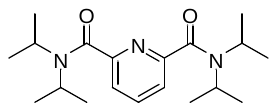

Following general procedure, the crude product was purified by silica gel column chromatography (20% EtOAc in petrol) to yield the title compound (**2**) as a white solid (67.0 mg, 80%). m.p. 161-162 °C (lit: 167 °C).

**^1^H NMR** (500 MHz, CDCl_3_) δ 7.83 (t, *J* = 7.8 Hz, 1H), 7.52 (d, *J* = 7.8 Hz, 2H), 3.89 (hept, *J* = 6.7 Hz, 2H), 3.53 (hept, *J* = 6.8 Hz, 2H), 1.53 (d, *J* = 6.8 Hz, 12H), 1.16 (d, *J* = 6.7 Hz, 12H). **^13^C NMR** (126 MHz, CDCl_3_) δ 168.5, 154.8, 138.2, 122.9, 51.1, 46.3, 20.8, 20.6; **IR** ν_max_ (cm^-1^): 2968, 1634, 1456, 1339, 1040, 772; **HRMS** (ESI) calculated for C_19_H_32_N_3_O_2_ [M+H]^+^ 334.2489, found 334.2498.

**Pyridine-2,6-diylbis(morpholinomethanone)**^2^ **(3)**

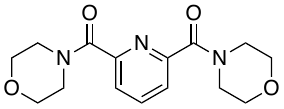

Following general procedure B, the crude product was purified by silica gel column chromatography (30% EtOAc in petrol) to yield the title compound (**3**) as a yellow solid (21.0 mg, 28%). m.p. 120-122 °C.

**^1^H NMR** (500 MHz, CDCl_3_) δ 7.95 (d, *J* = 7.8 Hz, 1H), 7.74 (d, *J* = 7.8 Hz, 2H), 3.87 – 3.78 (m, 8H), 3.67 (dd, *J* = 5.6, 3.7 Hz, 4H), 3.60 (dd, *J* = 5.6, 3.7 Hz, 4H); **^13^C NMR** (126 MHz, CDCl_3_) δ 166.8, 152.4, 138.6, 125.2, 67.1, 67.0, 47.9, 43.0; **IR** ν_max_ (cm^-1^): 2922, 2857, 1636, 1115, 752; **HRMS** (ESI) calculated for C_15_H_20_N_3_O_4_ [M+H]^+^ 306.1448, found 306.1443.

***N^2^,N^2^,N^6^,N^6^*-tetramethylpyridine-2,6-dicarboxamide**^3^ **(4)**

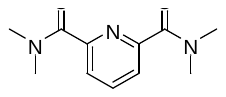

Following general procedure, the crude product was purified by silica gel column chromatography (20% EtOAc in petrol) to yield the title compound (**4**) as a white solid (23.0 mg, 42%). m.p. 144-145°C (lit: 144.5-148 °C).

**^1^H NMR** (500 MHz, CDCl_3_) δ 7.89 (t, *J* = 7.8 Hz, 1H), 7.66 (d, *J* = 7.8 Hz, 2H), 3.14 (s, 6H), 3.05 (s, 6H). **^13^C NMR** (126 MHz, CDCl_3_) δ 168.4, 153.3, 138.2, 124.2, 39.2, 35.9; **IR** ν_max_ (cm^-1^): 2928, 1634, 1506, 1391, 1103, 1082, 835; **HRMS** (ESI) calculated for C_11_H_16_N_3_O_2_ [M+H]^+^ 222.1237, found 222.1237.

***N^2^*,*N^6^*-diethylpyridine-2,6-dicarboxamide**^4^ **(5)**

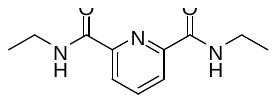

Following general procedure, the crude product was purified by silica gel column chromatography (25% EtOAc in petrol) to yield the title compound (**5**) as a white solid (30.0 mg, 87%). m.p. 178-179 °C (lit: 184.2-184.6 °C).

**^1^H NMR** (500 MHz, CDCl_3_) δ 8.35 (d, *J* = 7.8 Hz, 2H), 8.00 (t, *J* = 7.8 Hz, 1H), 7.92 (s, 2H), 3.52 (m, 4H), 1.26 (m, 6H). **^13^C NMR** (126 MHz, CDCl_3_) δ 163.6, 149.1, 139.1, 125.0, 34.6, 15.1; **IR** ν_max_ (cm^-1^): 3308, 2972, 1655, 1533, 1447; **HRMS** (ESI) calculated for C_11_H_16_N_3_O_2_ [M+H]^+^ 222.1237, found 222.1241.

**Pyridine-2,6-diylbis(pyrrolidin-1-ylmethanone)**^5^ **(6)**

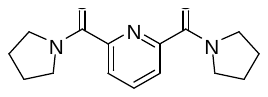

Following general procedure, the crude product was purified by silica gel column chromatography (20% EtOAc in petrol) to yield the title compound (**6**) as a yellow solid (45.0 mg, 66%). m.p. 78-80 °C.

**^1^H NMR** (500 MHz, CDCl_3_) δ 7.92 – 7.82 (m, 3H), 3.66 (m, 8H), 1.97 – 1.86 (m, 8H). **^13^C NMR** (126 MHz, CDCl_3_) δ 166.2, 153.1, 137.8, 124.9, 49.2, 47.0, 26.7, 24.1; **IR** ν_max_ (cm^-1^): 2978, 1622, 1456, 1412, 1217, 752; **HRMS** (ESI) calculated for C_15_H_20_N_3_O_2_ [M+H]^+^ 274.1550, found 274.1558.

***N^2^,N^6^*-Di(prop-2-yn-1-yl)pyridine-2,6-dicarboxamide**^6^ **(7)**

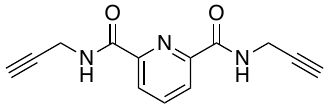

To a solution of pyridine-2,6-dicarbonyl dichloride (51.0 mg, 250 µmol, 1.00 eq) and Et_3_N (70.0 µL, 500 µmol, 2.00 eq) was added propargylamine (32.0 µL, 500 µmol, 2.00 eq). The solution was stirred at room temperature for 16 h then diluted with CH_2_Cl_2_ (10 mL) and washed with brine (10 mL). The organic phase was dried over anhydrous MgSO_4_, filtered and concentrated under reduced pressure. The crude residue was purified by silica gel column chromatography (35% EtOAc in petrol) to yield the title compound (**7**) as a white solid (37.0 mg, 62%). m.p. 62-64 °C (lit: 63-64 °C).

**^1^H NMR** (500 MHz, CDCl_3_) δ 8.40 (d, *J* = 7.8 Hz, 2H), 8.06 (t, *J* = 7.8 Hz, 1H), 7.93 (s, 2H), 4.34 (dd, *J* = 5.7, 2.5 Hz, 4H), 2.33 – 2.28 (m, 2H); **^13^C NMR** (126 MHz, CDCl_3_) δ 163.3, 148.6, 139.3, 125.7, 79.4, 72.0, 29.4; **IR** ν_max_ (cm^-1^): 3294, 1661, 1522, 1445; **HRMS** (ESI) calculated for C_13_H_11_N_3_NaO_2_ [M+Na]^+^ 264.0743, found 264.0747.

**Synthesis of Esters**

**Dimethyl pyridine-2,6-dicarboxylate**^7^ **(8)**

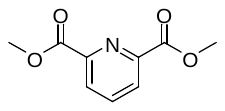

To a solution of pyridine-2,6-dicarbonyl dichloride (408 mg, 2.00 mmol) in CH_2_Cl_2_ (10 mL) was added MeOH (243 µL, 6.00 mmol) followed by Et_3_N (1.12 mL, 8.00 mmol). The solution was stirred at room temperature for 2 h, diluted with CH_2_Cl_2_ (50 mL) and washed with water (50 mL). The organic phase was dried over anhydrous MgSO_4_, filtered and concentrated under reduced pressure. The crude residue was purified by silica gel column chromatography (0-40% EtOAc in petrol) to yield the title compound (**8**) as an off-white solid (281 mg, 72%). m.p. 121-122 °C.

**^1^H NMR** (500 MHz, CDCl_3_) δ 8.32 (d, *J* = 7.8 Hz, 2H), 8.03 (t, *J* = 7.8 Hz, 1H), 4.03 (s, 6H). **^13^C NMR** (126 MHz, CDCl_3_) δ 165.2, 148.4, 138.5, 128.2, 53.3; **IR** ν_max_ (cm^-1^): 1751, 1715, 1450, 1323, 1290, 1252, 1144, 739; **HRMS** (ESI) calculated for C_9_H_10_NO_4_ [M+H]^+^ 196.0604, found 196.0613.

**Diethyl pyridine-2,6-dicarboxylate**^8^ **(9)**

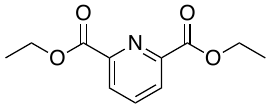

To a solution of pyridine-2,6-dicarbonyl dichloride (408 mg, 2.00 mmol) in CH_2_Cl_2_ (10 mL) was added EtOH (350 µL, 6.00 mmol) followed by Et_3_N (1.12 mL, 8.00 mmol). The solution was stirred at room temperature for 1 h, diluted with CH_2_Cl_2_ (50 mL) and washed with water (50 mL). The organic phase was dried over anhydrous MgSO_4_, filtered and concentrated under reduced pressure. The crude residue was purified by silica gel column chromatography (0-30% EtOAc in petrol) to yield the title compound (**9**) as an off-white solid (273 mg, 61%). m.p. 43-45 °C (lit: 28 °C).

**^1^H NMR** (500 MHz, CDCl_3_) δ 8.27 (d, *J* = 7.8 Hz, 2H), 7.99 (t, *J* = 7.8 Hz, 1H), 4.48 (q, *J* = 7.2 Hz, 4H), 1.45 (t, *J* = 7.1 Hz, 6H); **^13^C NMR** (126 MHz, CDCl_3_) δ 164.7, 148.7, 138.3, 127.9, 62.4, 14.3. **IR** ν_max_ (cm^-1^): 1744, 1719, 1369, 1321, 1242, 1138, 1024, 752; **HRMS** (ESI) calculated for C_11_H_14_NO_4_ [M+H]^+^ 224.0917, found 224.0919.

**Dipropyl pyridine-2,6-dicarboxylate**^9^ **(10)**

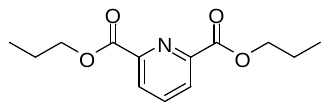

To a solution of pyridine-2,6-dicarbonyl dichloride (240 mg, 1.18 mmol) in CH_2_Cl_2_ (10 mL) was added n-propanol (264 µL, 3.53 mmol) followed by Et_3_N (658 µL, 4.72 mmol). The solution was stirred at room temperature for 2 h, diluted with CH_2_Cl_2_ (50 mL) and washed with water (50 mL). The organic phase was dried over anhydrous MgSO_4_, filtered and concentrated under reduced pressure. The crude residue was purified by silica gel column chromatography (0-30% EtOAc in petrol) to yield the title compound (**10**) as a pale yellow oil (162 mg, 55%).

**^1^H NMR** (400 MHz, CDCl_3_) δ 8.27 (d, *J* = 7.8 Hz, 2H), 7.99 (t, *J* = 7.8 Hz, 1H), 4.38 (t, *J* = 6.9 Hz, 4H), 1.95 – 1.72 (m, 4H), 1.05 (t, *J* = 7.4 Hz, 6H). **^13^C NMR** (101 MHz, CDCl_3_) δ 164.8, 148.8, 138.3, 127.8, 67.9, 22.1, 10.5; **IR** ν_max_ (cm^-1^): 2968, 1748, 1719, 1323, 1240, 1165, 1140, 752; **HRMS** (ESI) calculated for C_13_H_18_NO_4_ [M+H]^+^ 252.1230, found 252.1236.

**Dibutyl pyridine-2,6-dicarboxylate**^10^ **(11)**

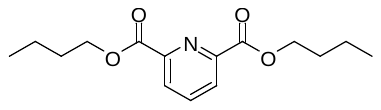

To a solution of pyridine-2,6-dicarbonyl dichloride (408 mg, 2.00 mmol) in CH_2_Cl_2_ (10 mL) was added n-butanol (550 µL, 6.00 mmol) followed by Et_3_N (1.12 mL, 8.00 mmol). The solution was stirred at room temperature for 2 h, diluted with CH_2_Cl_2_ (50 mL) and washed with water (50 mL). The organic phase was dried over anhydrous MgSO_4_, filtered and concentrated under reduced pressure. The crude residue was purified by silica gel column chromatography (0-30% EtOAc in petrol) to yield the title compound (**11**) as an off-white solid (405 mg, 75%). m.p. 62-64 °C (lit: 63-64 °C).

**^1^H NMR** (400 MHz, CDCl_3_) δ 8.26 (d, *J* = 7.8 Hz, 2H), 7.99 (t, *J* = 7.8 Hz, 1H), 4.43 (t, *J* = 6.8 Hz, 4H), 1.87 – 1.75 (m, 4H), 1.54 – 1.43 (m, 4H), 0.99 (t, *J* = 7.4 Hz, 6H). **^13^C NMR** (101 MHz, CDCl_3_) δ 164.8, 148.9, 138.3, 127.8, 66.2, 30.8, 19.3, 13.9; **IR** ν_max_ (cm^-1^): 2955, 1740, 1578, 1290, 1248, 766; **HRMS** (ESI) calculated for C_15_H_22_NO_4_ [M+H]^+^ 280.1543, found 280.1548.

**Dipentyl pyridine-2,6-dicarboxylate (12)**

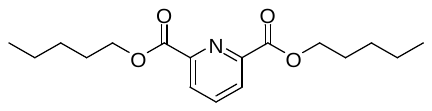

To a solution of pyridine-2,6-dicarbonyl dichloride (550 mg, 2.70 mmol) in CH_2_Cl_2_ (20 mL) was added n-pentanol (877 µL, 8.10 mmol) followed by Et_3_N (1.51 mL, 10.8 mmol). The solution was stirred at room temperature for 2 h, diluted with CH_2_Cl_2_ (50 mL) and washed with water (50 mL). The organic phase was dried over anhydrous MgSO_4_, filtered and concentrated under reduced pressure. The crude residue was purified by silica gel column chromatography (0-30% EtOAc in petrol) to yield the title compound (**12**) as an off-white solid (340 mg, 41%). m.p. 39-41 °C.

**^1^H NMR** (500 MHz, CDCl_3_) δ 8.27 (d, *J* = 7.8 Hz, 2H), 8.00 (t, *J* = 7.8 Hz, 2H), 4.42 (t, *J* = 6.9 Hz, 4H), 1.84 (dd, *J* = 8.1, 6.9 Hz, 4H), 1.49 – 1.34 (m, 8H), 0.93 (t, *J* = 7.1 Hz, 6H); **^13^C NMR** (126 MHz, CDCl_3_) δ 164.8, 148.9, 138.2, 127.8, 66.5, 28.4, 28.2, 22.5, 14.1; **IR** ν_max_ (cm^-1^): 2957, 1741, 1734, 1719, 1325, 1240, 1144; **HRMS** (ESI) calculated for C_17_H_26_NO_4_ [M+H]^+^ 308.1856, found 308.1866.

**Dihexyl pyridine-2,6-dicarboxylate (13)**

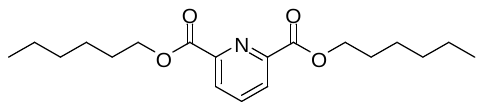

To a solution of pyridine-2,6-dicarbonyl dichloride (550 mg, 2.70 mmol) in CH_2_Cl_2_ (20 mL) was added n-hexanol (1.02 mL, 8.10 mmol) followed by Et_3_N (1.51 mL, 10.8 mmol). The solution was stirred at room temperature for 2 h, diluted with CH_2_Cl_2_ (50 mL) and washed with water (50 mL). The organic phase was dried over anhydrous MgSO_4_, filtered and concentrated under reduced pressure. The crude residue was purified by silica gel column chromatography (0-30% EtOAc in petrol) to yield the title compound (**13**) as a low melting colourless solid (608 mg, 73%).

**^1^H NMR** (400 MHz, CDCl_3_) δ 8.25 (d, *J* = 7.8 Hz, 2H), 7.99 (t, *J* = 7.8 Hz, 1H), 4.40 (t, *J* = 7.0 Hz, 4H), 1.88 – 1.76 (m, 4H), 1.49 – 1.39 (m, 4H), 1.39 – 1.26 (m, 8H), 0.93 – 0.85 (m, 6H); **^13^C NMR** (101 MHz, CDCl_3_) δ 164.8, 148.8, 138.2, 127.8, 66.5, 31.6, 28.6, 25.7, 22.6, 14.1; **IR** ν_max_ (cm^-1^): 2955, 1740, 1576, 1290, 1250, 764; **HRMS** (ESI) calculated for C_19_H_30_NO_4_ [M+H]^+^ 336.2169, found 336.2162.

**Bis(2-bromoethyl) pyridine-2,6-dicarboxylate**^11^ **(14)**

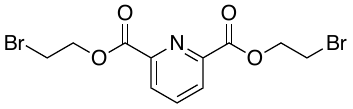

To a solution of pyridine-2,6-dicarbonyl dichloride (550 mg, 2.70 mmol) in CH_2_Cl_2_ (20 mL) was added 2-bromoethanol (574 µL, 8.10 mmol) followed by Et_3_N (1.51 mL, 10.8 mmol). The solution was stirred at room temperature for 2 h, diluted with CH_2_Cl_2_ (50 mL) and washed with water (50 mL). The organic phase was dried over anhydrous MgSO_4_, filtered and concentrated under reduced pressure. The crude residue was purified by silica gel column chromatography (10-50% EtOAc in petrol) to yield the title compound (**14**) as an off-white solid (532 mg, 52%). m.p. 104-106 °C.

**^1^H NMR** (400 MHz, CDCl_3_) δ 8.33 (d, *J* = 7.8 Hz, 2H), 8.05 (t, *J* = 7.8 Hz, 1H), 4.73 (t, *J* = 6.5 Hz, 4H), 3.70 (t, *J* = 6.5 Hz, 4H); **^13^C NMR** (101 MHz, CDCl_3_) δ 164.0, 148.2, 138.6, 128.5, 65.3, 28.1; **IR** ν_max_ (cm^-1^): 2924, 1748, 1724, 1379, 1317, 1242, 1142, 750; **HRMS** (ESI) calculated for C_11_H_11_Br_2_NNaO_4_ [M+Na]^+^ 401.8947, found 401.8944.

**Bis(2-chloroethyl) pyridine-2,6-dicarboxylate**^12^ **(15)**

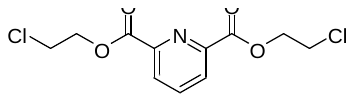

To a solution of pyridine-2,6-dicarbonyl dichloride (408 mg, 2.00 mmol) in CH_2_Cl_2_ (20 mL) was added 2-chloroethanol (402 µL, 6.00 mmol) followed by Et_3_N (1.12 mL, 8.00 mmol). The solution was stirred at room temperature for 2 h, diluted with CH_2_Cl_2_ (50 mL) and washed with water (50 mL). The organic phase was dried over anhydrous MgSO_4_, filtered and concentrated under reduced pressure. The crude residue was purified by silica gel column chromatography (0-30% EtOAc in petrol) to yield the title compound (**15**) as an off-white solid (342 mg, 58%). m.p. 94-96 °C.

**^1^H NMR** (400 MHz, CDCl_3_) δ 8.33 (d, *J* = 7.8 Hz, 2H), 8.05 (t, *J* = 7.8 Hz, 1H), 4.68 (t, *J* = 6.0 Hz, 4H), 3.87 (t, *J* = 6.0 Hz, 4H). **^13^C NMR** (101 MHz, CDCl_3_) δ 164.1, 148.2, 138.6, 128.5, 65.5, 41.2; **IR** ν_max_ (cm^-1^): 2957, 1749, 1719, 1319, 1236, 1142, 750; **HRMS** (ESI) calculated for C_11_H_12_Cl_2_NO_4_ [M+H]^+^ 292.0138, found 292.0132.

**Di(prop-2-yn-1-yl) pyridine-2,6-dicarboxylate**^13^ **(16)**

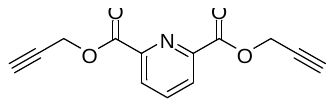

To a solution of pyridine-2,6-dicarbonyl dichloride (550 mg, 2.70 mmol) in CH_2_Cl_2_ (20 mL) was added propargyl alcohol (466 µL, 8.10 mmol) followed by Et_3_N (1.50 mL, 10.8 mmol). The solution was stirred at room temperature for 2 h, diluted with CH_2_Cl_2_ (50 mL) and washed with water (50 mL). The organic phase was dried over anhydrous MgSO_4_, filtered and concentrated under reduced pressure to yield the title compound (**16**) as an off-white solid (562 mg, 86%). m.p. 123-124 °C (lit: 124-125 °C).

**^1^H NMR** (500 MHz, CDCl_3_) δ 8.34 (d, *J* = 7.8 Hz, 2H), 8.05 (t, *J* = 7.8 Hz 1H), 5.02 (d, *J* = 2.5 Hz, 4H), 2.54 (t, *J* = 2.5 Hz, 2H). **^13^C NMR** (126 MHz, CDCl_3_) δ 163.8, 148.0, 138.6, 128.6, 77.2, 75.8, 53.7; **IR** ν_max_ (cm^-1^): 3258, 1732, 1315, 1134, 1121, 1078; **HRMS** (ESI) calculated for C_13_H_10_NO_4_ [M+H]^+^ 244.0604, found 244.0609.

**Di(but-3-yn-1-yl) pyridine-2,6-dicarboxylate (17)**

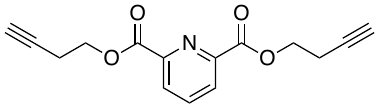

To a solution of pyridine-2,6-dicarbonyl dichloride (408 mg, 2.00 mmol) in CH_2_Cl_2_ (20 mL) was added 3-butyn-1-ol (454 µL, 6.00 mmol) followed by Et_3_N (1.12 mL, 8.00 mmol). The solution was stirred at room temperature for 1 h, diluted with CH_2_Cl_2_ (50 mL) and washed with water (50 mL). The organic phase was dried over anhydrous MgSO_4_, filtered and concentrated under reduced pressure. The crude residue was purified by silica gel column chromatography (0-30% EtOAc in petrol) to yield the title compound (**17**) as an off-white solid (415 mg, 77%). m.p. 86-88 °C.

**^1^H NMR** (500 MHz, CDCl_3_) δ 8.29 (d, *J* = 7.8 Hz, 2H), 8.02 (t, *J* = 7.8 Hz, 1H), 4.54 (t, *J* = 7.2 Hz, 4H), 2.74 (td, *J* = 7.1, 2.7 Hz, 4H), 2.04 (t, *J* = 2.7 Hz, 2H); **^13^C NMR** (126 MHz, CDCl_3_) δ 164.3, 148.4, 138.4, 128.2, 79.7, 70.5, 63.8, 19.1; **IR** ν_max_ (cm^-1^): 3289, 2959, 1748, 1717, 1325, 1240, 1144, 1084, 752; **HRMS** (ESI) calculated for C_15_H_14_NO_4_ [M+H]^+^ 272.0917, found 272.0926.

**Bis(2,2,2-trifluoroethyl) pyridine-2,6-dicarboxylate**^12^ **(18)**

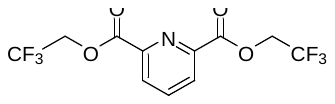

To a solution of pyridine-2,6-dicarbonyl dichloride (408 mg, 2.00 mmol) in CH_2_Cl_2_ (20 mL) was added 2,2,2-trifluoroethanol (432 µL, 6.00 mmol) followed by Et_3_N (1.12 mL, 8.00 mmol). The solution was stirred at room temperature for 2 h, diluted with CH_2_Cl_2_ (50 mL) and washed with water (50 mL). The organic phase was dried over anhydrous MgSO_4_, filtered and concentrated under reduced pressure. The crude residue was purified by silica gel column chromatography (0-30% EtOAc in petrol) to yield the title compound (**18**) as an off-white solid (457 mg, 69%). m.p. 106-108 °C.

**^1^H NMR** (500 MHz, CDCl_3_) δ 8.35 (d, *J* = 7.8 Hz, 2H), 8.13 – 8.06 (appt t, *J* = 7.8 Hz, 1H), 4.81 (q, *J* = 8.3 Hz, 4H). **^13^C NMR** (126 MHz, CDCl_3_) δ 162.8, 147.4, 138.9, 129.1, 122.9 (q, *J* = 277.5 Hz), 61.7 (q, *J* = 37.1 Hz); **IR** ν_max_ (cm^-1^): 3061, 1757, 1576, 1260, 1153, 962, 760; **HRMS** (ESI) calculated for C_11_H_7_F_6_NNaO_4_ [M+Na]^+^ 354.0171, found 354.0176.

**Diisopropyl pyridine-2,6-dicarboxylate (19)**

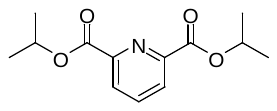

To a solution of pyridine-2,6-dicarbonyl dichloride (408 mg, 2.00 mmol) in CH_2_Cl_2_ (20 mL) was added isopropanol (459 µL, 6.00 mmol) followed by Et_3_N (1.12 mL, 8.00 mmol). The solution was stirred at room temperature for 2 h, diluted with CH_2_Cl_2_ (50 mL) and washed with water (50 mL). The organic phase was dried over anhydrous MgSO_4_, filtered and concentrated under reduced pressure. The crude residue was purified by silica gel column chromatography (0-30% EtOAc in petrol) to yield the title compound (**19**) as an off white solid (292 mg, 58%). m.p. 64-65 °C.

**^1^H NMR** (400 MHz, CDCl_3_) δ 8.24 (d, *J* = 7.8 Hz, 2H), 7.97 (t, *J* = 7.8 Hz, 1H), 5.33 (hept, *J* = 6.3 Hz, 2H), 1.43 (d, *J* = 6.3 Hz, 12H). **^13^C NMR** (101 MHz, CDCl_3_) δ 164.2, 149.2, 138.1, 127.7, 70.2, 22.0; **IR** ν_max_ (cm^-1^): 2926, 1748, 1734, 1717, 1506, 1244, 1105, 754; **HRMS** (ESI) calculated for C_13_H_16_NO_4_ [M+H]^+^ 252.1230, found 252.1235.

**Bis(3-methoxybutyl) pyridine-2,6-dicarboxylate (20)**

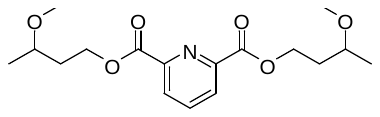

To a solution of pyridine-2,6-dicarbonyl dichloride (500 mg, 2.45 mmol) in CH_2_Cl_2_ (20 mL) was added 3-methoxy-1-butanol (825 µL, 7.35 mmol) followed by Et_3_N (1.37 mL, 9.80 mmol). The solution was stirred at room temperature for 1 h, diluted with CH_2_Cl_2_ (50 mL) and washed with water (50 mL). The organic phase was dried over anhydrous MgSO_4_, filtered and concentrated under reduced pressure. The crude residue was purified by silica gel column chromatography (0-30% EtOAc in petrol) to yield the title compound (**20**) as a low melting colourless solid (319 mg, 38%).

**^1^H NMR** (400 MHz, CDCl_3_) δ 8.27 (d, *J* = 7.8 Hz, 2H), 8.00 (dd, *J* = 8.1, 7.5 Hz, 1H), 4.53 (t, *J* = 6.8 Hz, 4H), 3.53 (h, *J* = 6.2 Hz, 2H), 3.35 (s, 6H), 2.06 – 1.92 (m, 4H), 1.23 (d, *J* = 6.1 Hz, 6H); **^13^C NMR** (101 MHz, CDCl_3_) δ 164.8, 148.8, 138.3, 127.9, 73.9, 63.4, 56.3, 35.6, 19.3; **IR** ν_max_ (cm^-1^): 2970, 2928, 1748, 1719, 1325, 1242, 1146, 1082, 752; **HRMS** (ESI) calculated for C_17_H_26_NO_6_ [M+H]^+^ 340.1755, found 340.1759.

**Synthesis of Pyridine-2,6-dicarbaldehyde**^14^ **(21)**

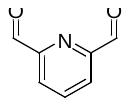

To a solution of 2,6-pyridinedimethanol (417 mg, 3.00 mmol, 1.00 eq) in 1,4-dioxane (10.0 mL) was added SeO_2_ (333 mg, 3.00 mmol, 1.00 eq). The resultant mixture was stirred at room temperature for 16 h then concentrated under a stream of nitrogen. The crude product was dissolved in CH_2_Cl_2_, filtered and concentrated under reduced pressure. Purification by silica gel column chromatography (0-30% EtOAc/Petrol) yielded the title compound (**21**) as a white solid (240 mg, 59%). m.p. 120-122 °C (lit: 124 °C).

**^1^H NMR** (500 MHz, CDCl_3_) δ 10.18 (d, *J* = 0.7 Hz, 2H), 8.19 (dd, *J* = 7.7, 0.7 Hz, 2H), 8.11 – 8.06 (m, 1H). **^13^C NMR** (126 MHz, CDCl_3_) δ 192.5, 153.2, 138.5, 125.5; **IR** ν_max_ (cm^-1^): 2860, 1717, 1348, 1261, 1086, 804; **HRMS** (ESI) calculated for C_7_H_5_NNaO_2_ [M+Na]^+^ 158.0212, found 158.0205.

**Supplementary Data S1. NMR spectra of compounds.**

**Compound 1**

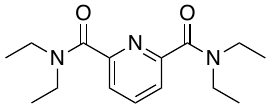

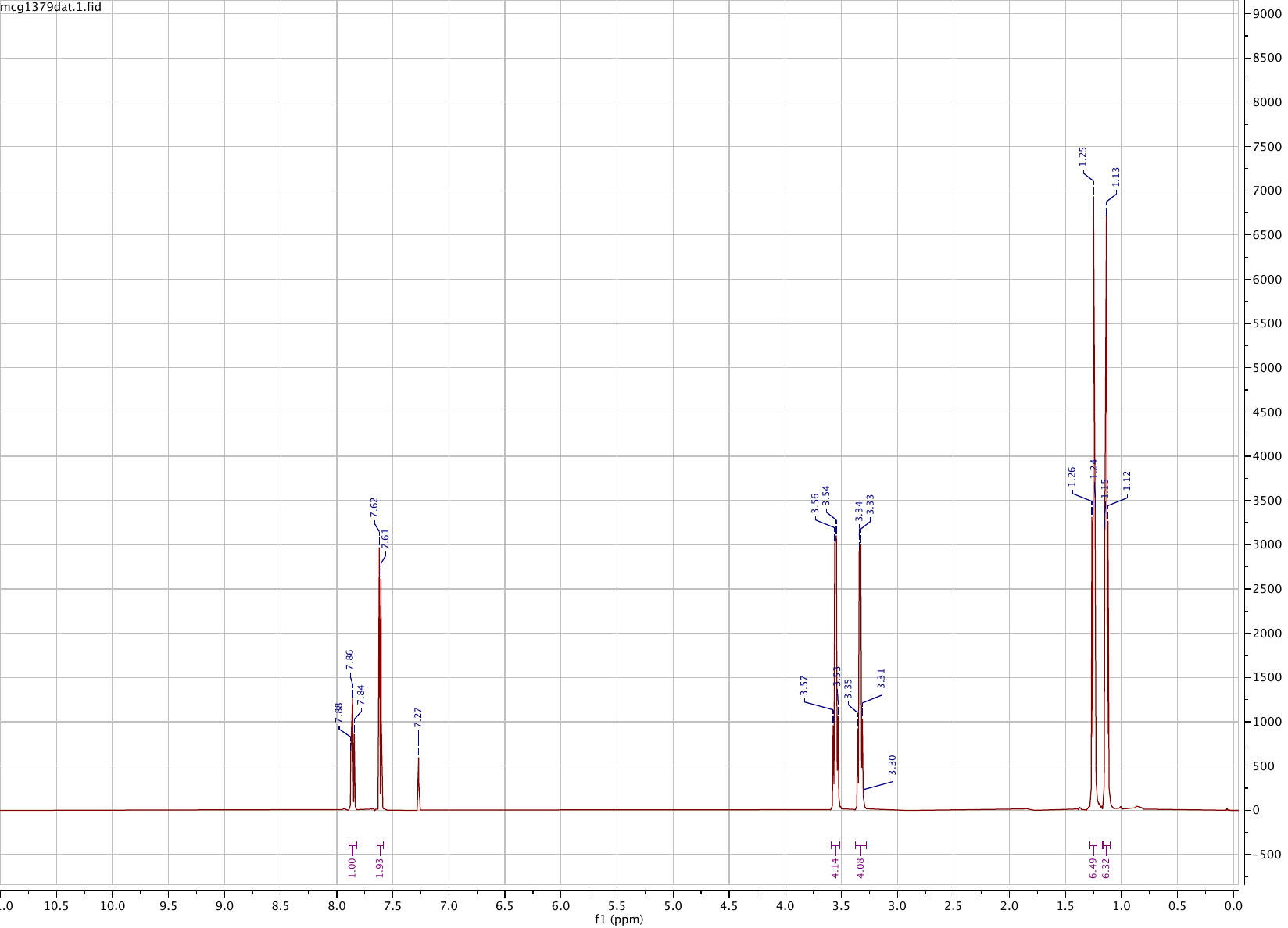

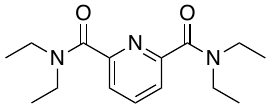

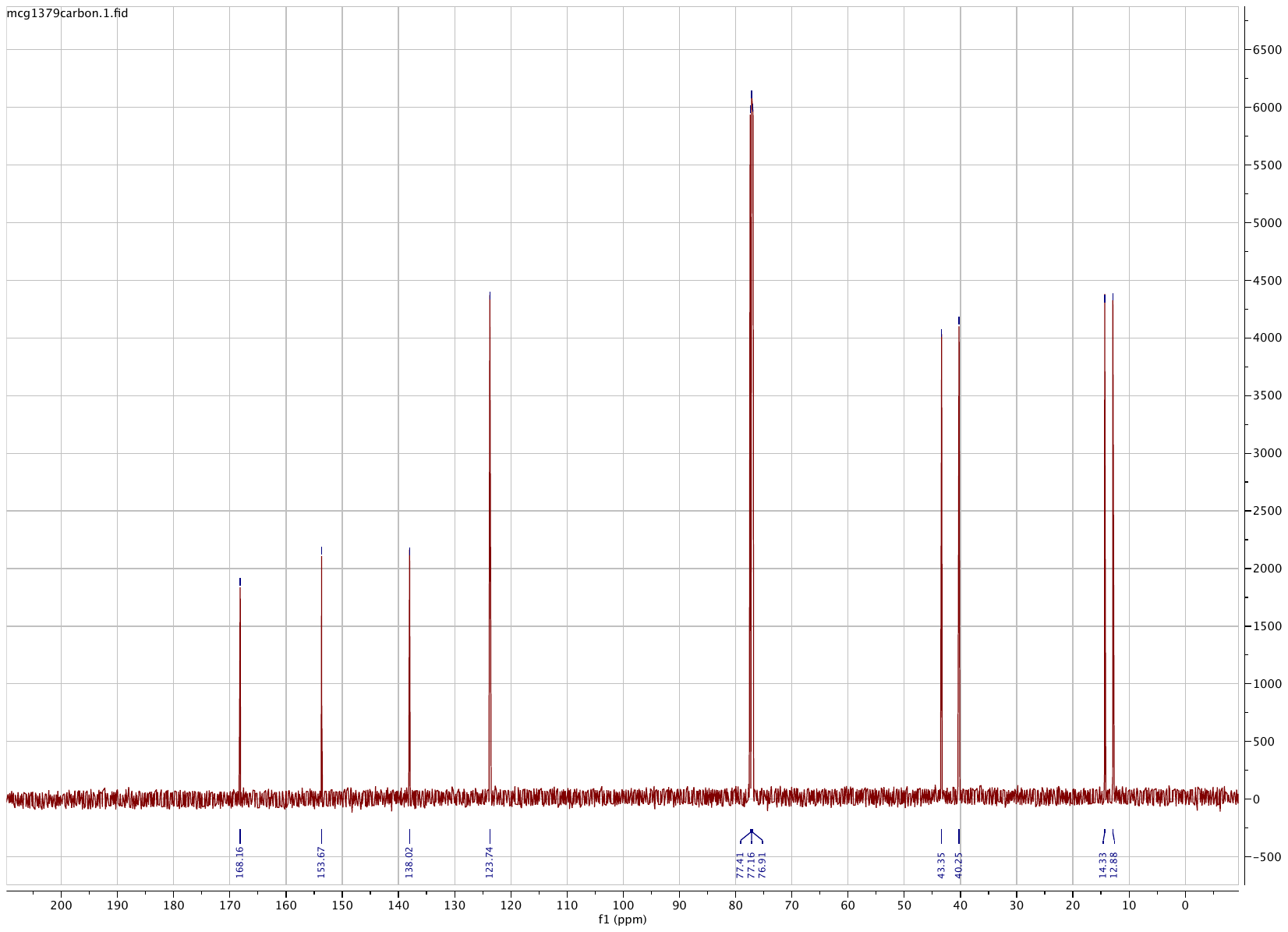

**Compound 2**

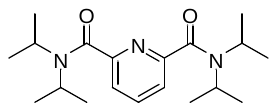

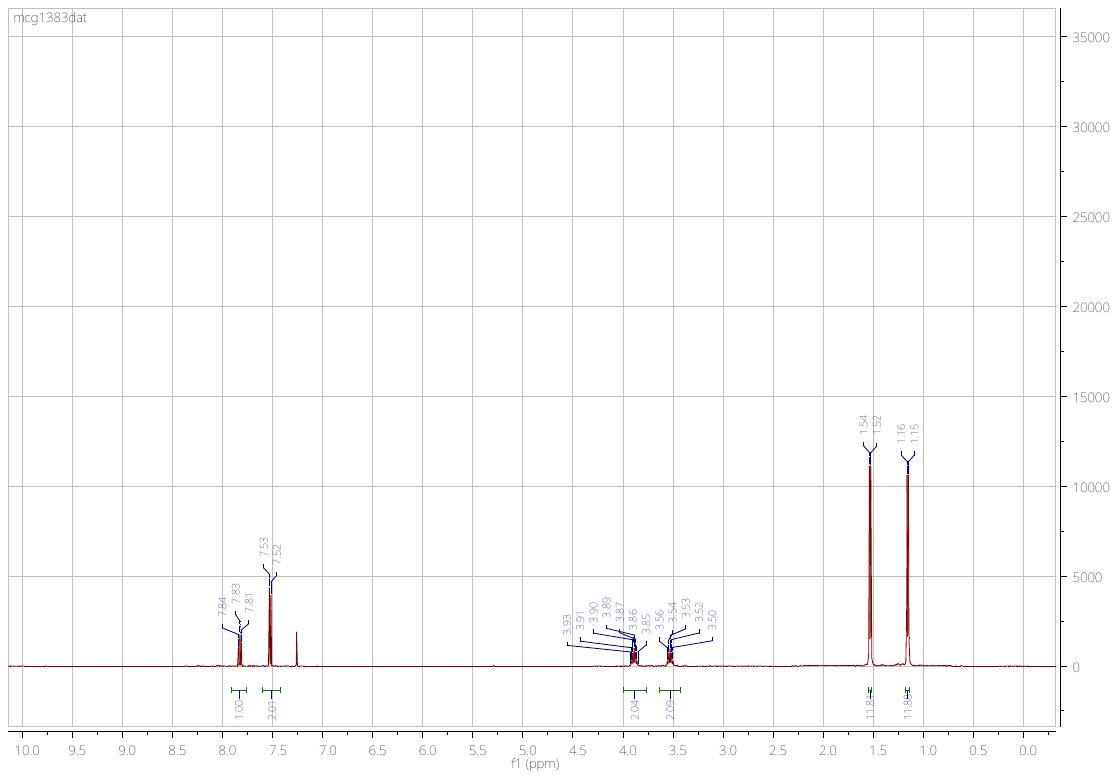

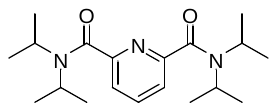

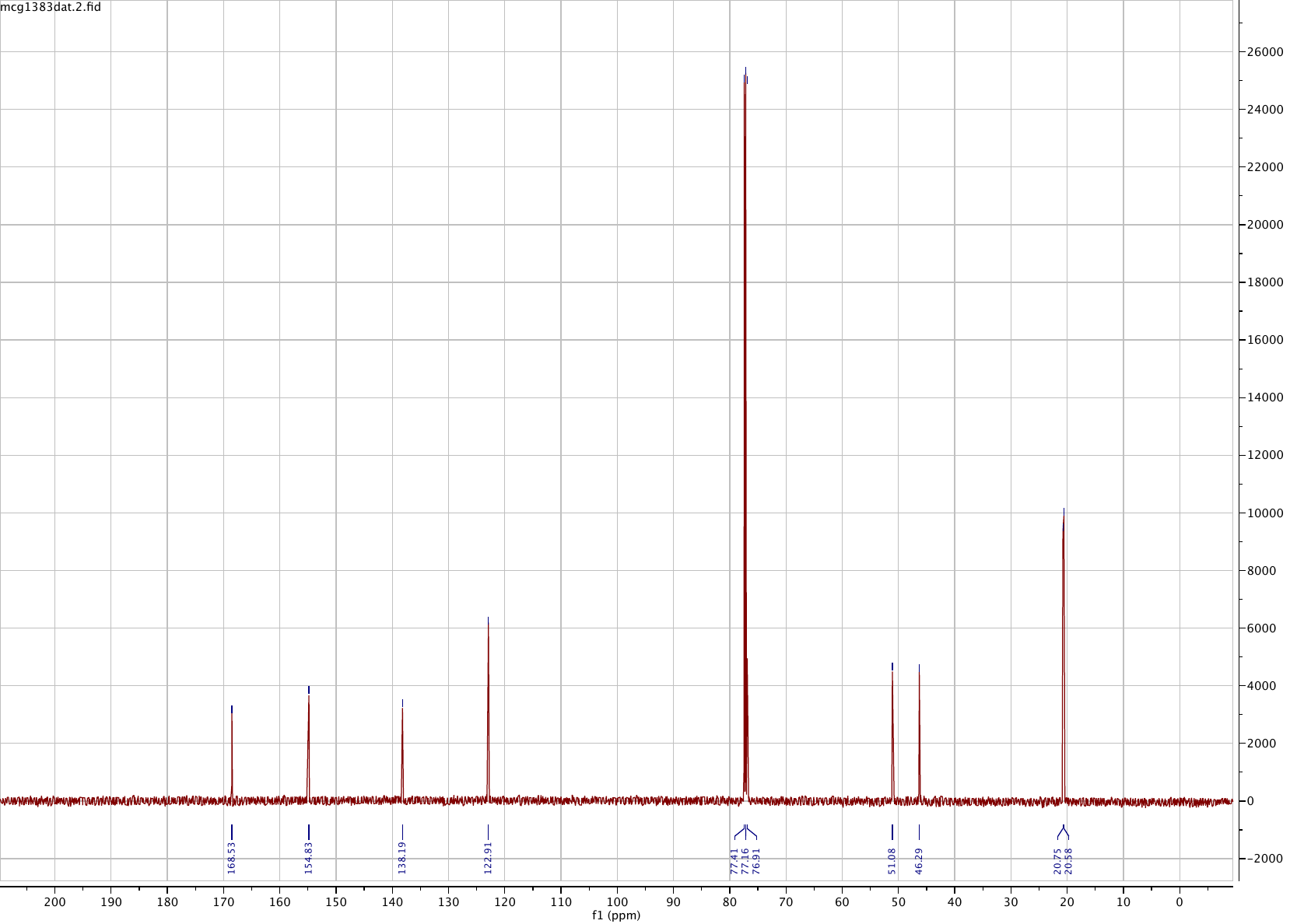

**Compound 3**

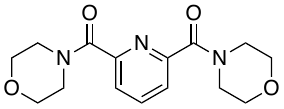

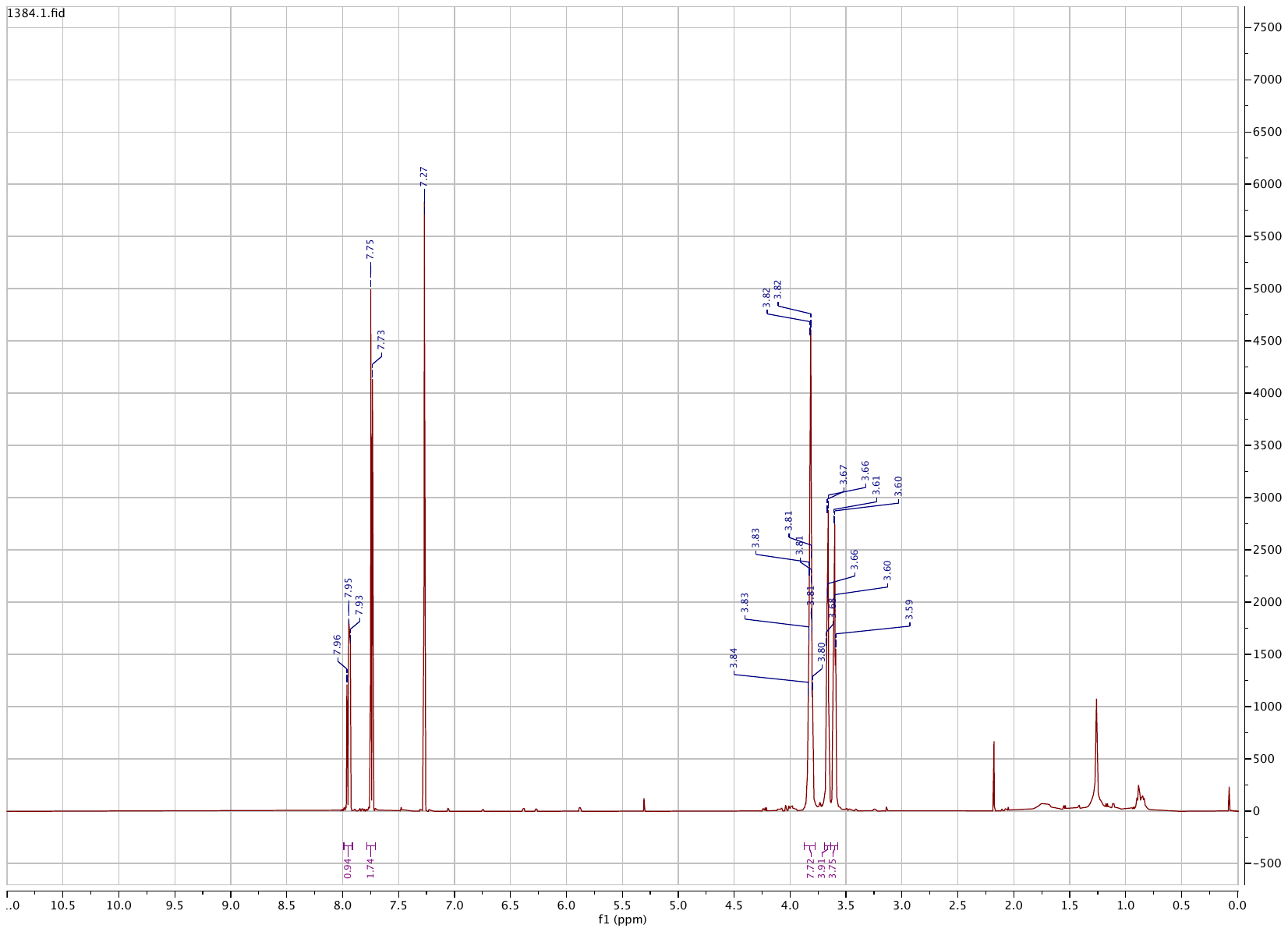

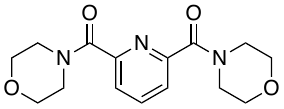

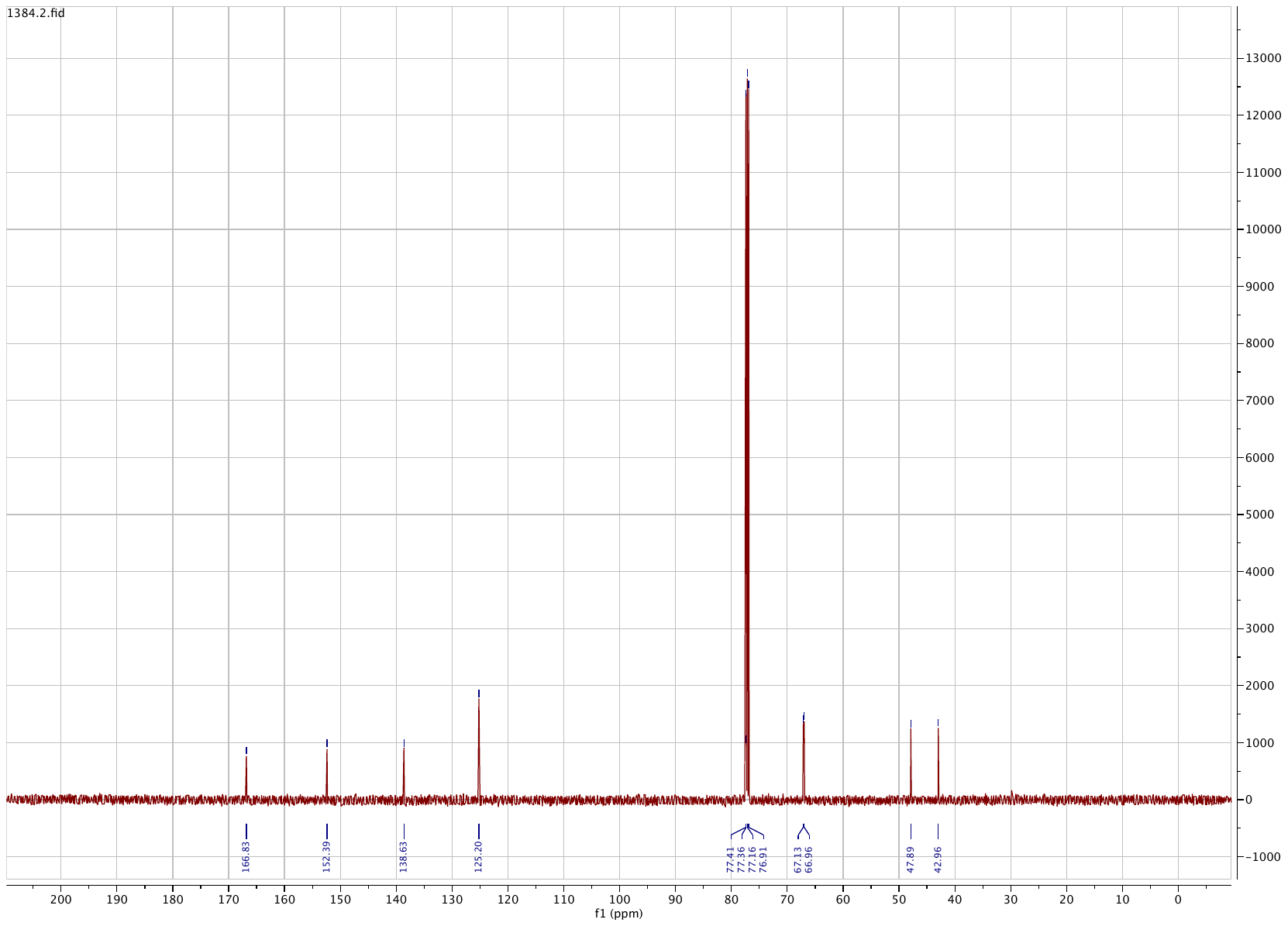

**Compound 4**

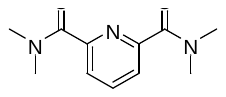

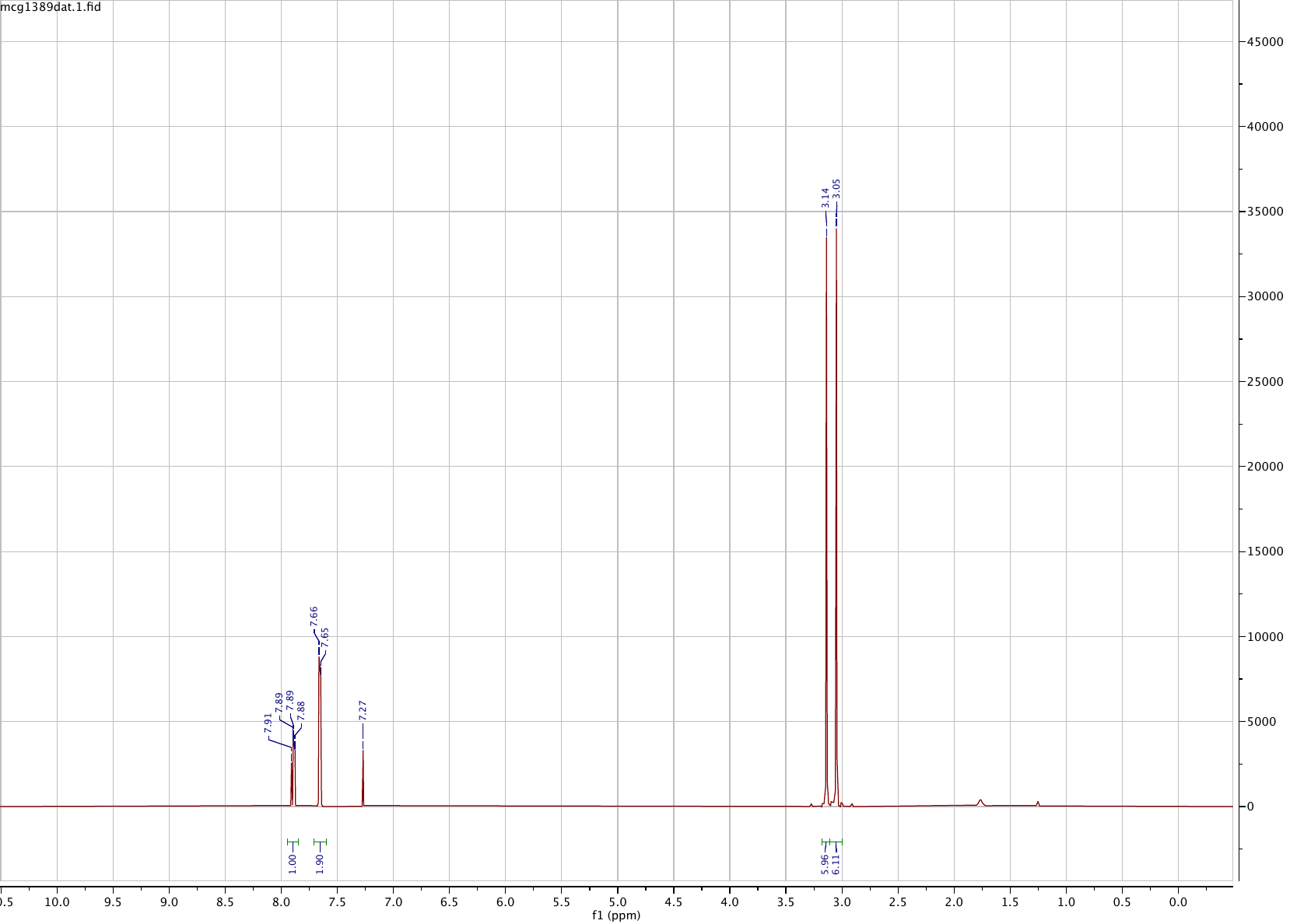

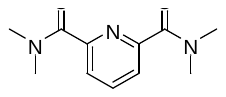

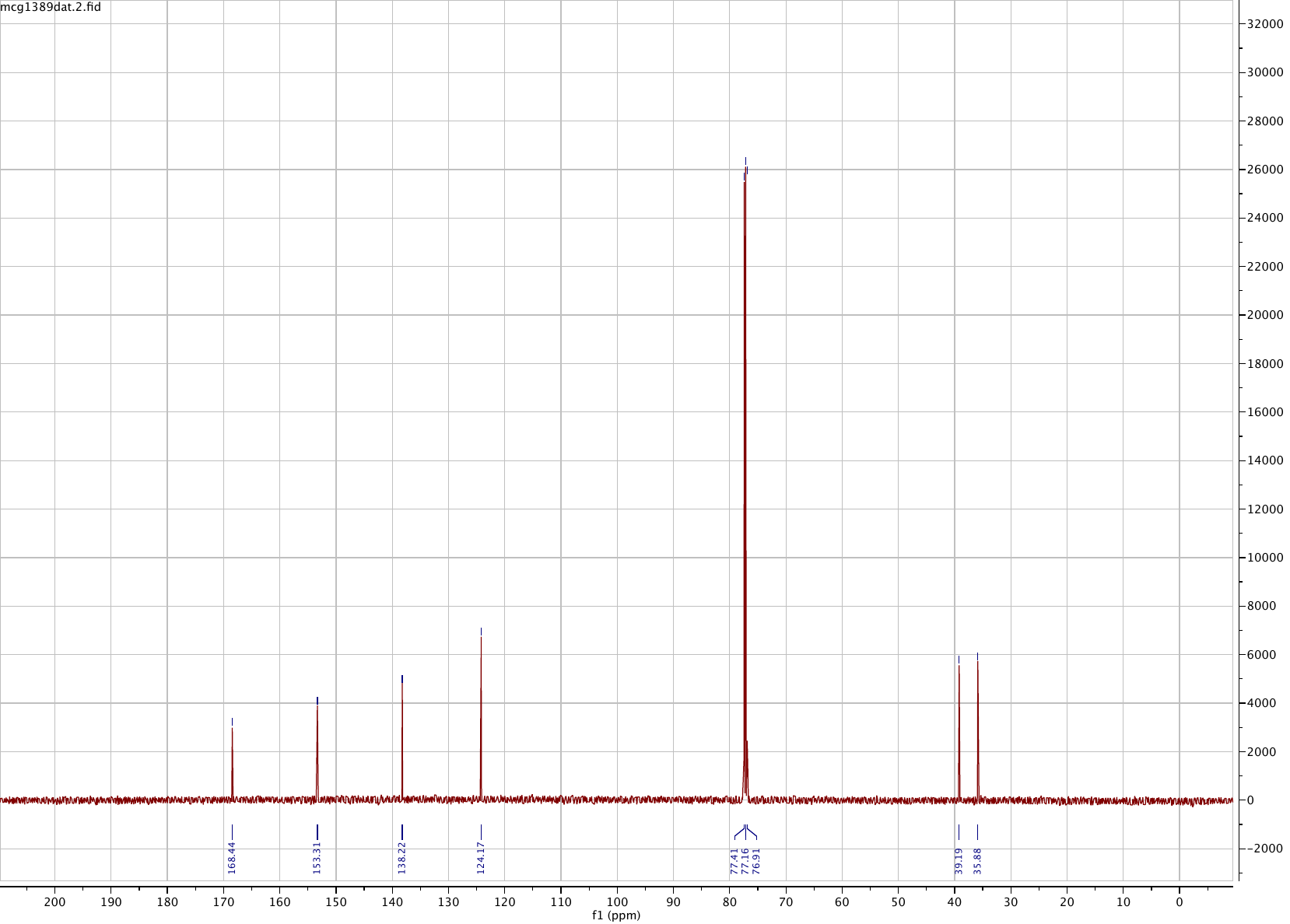

**Compound 5**

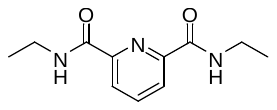

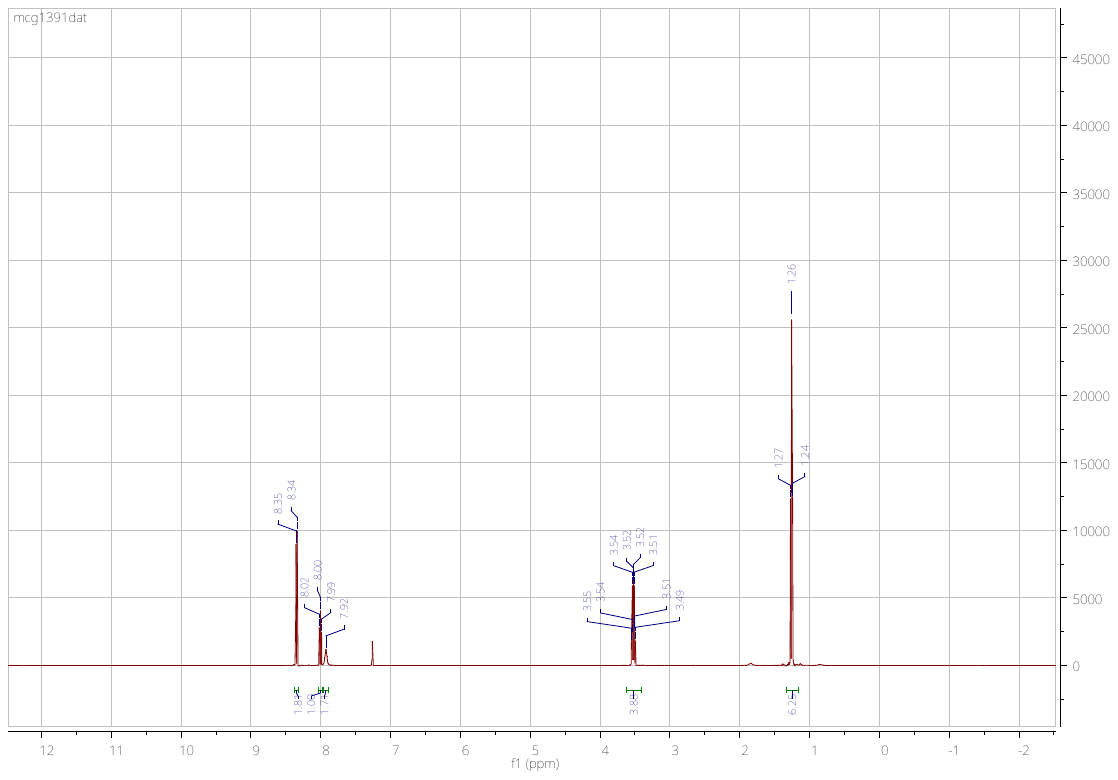

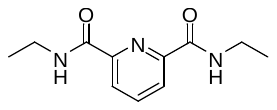

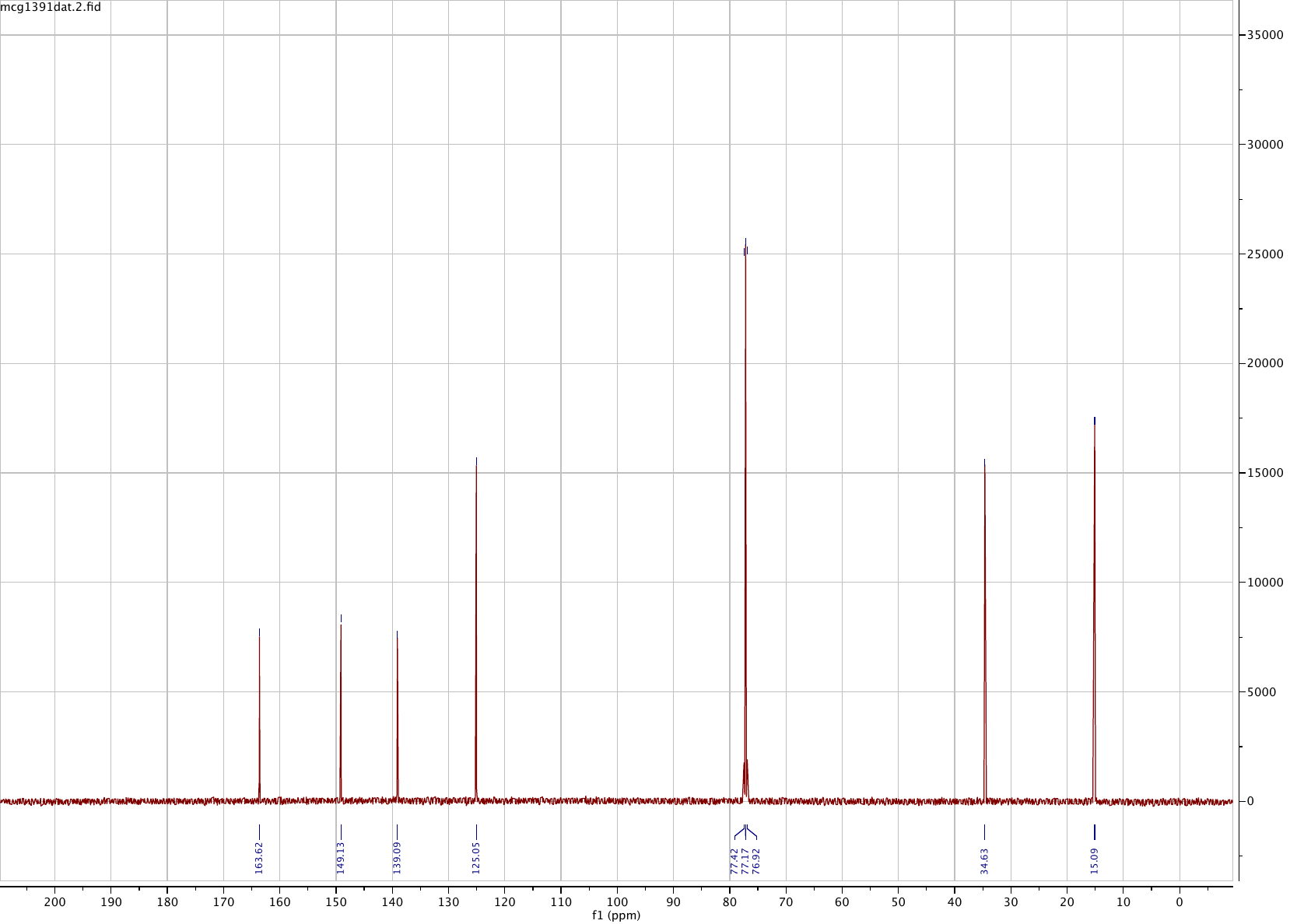

**Compound 6**

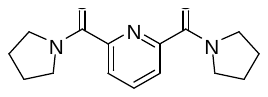

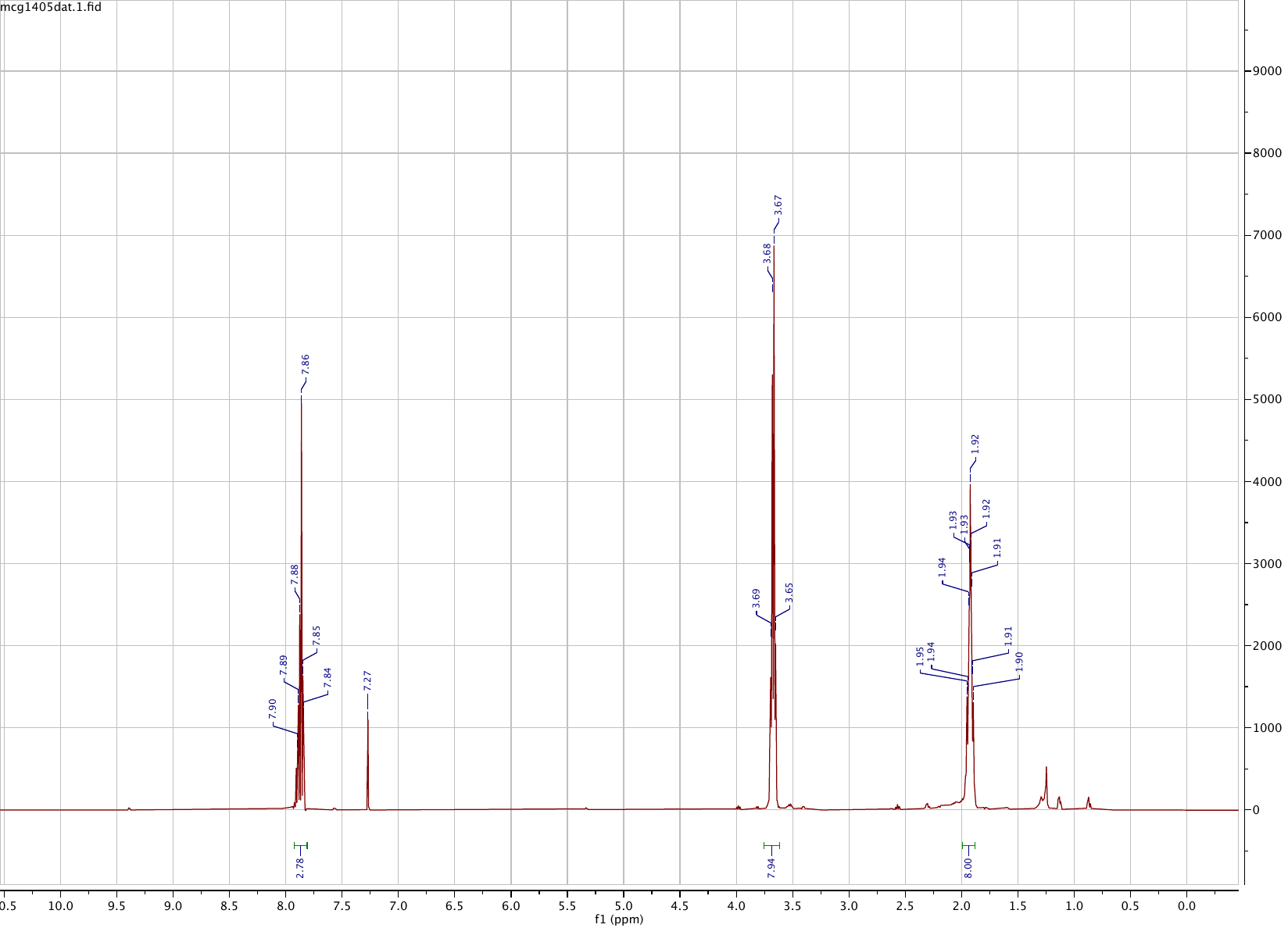

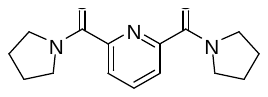

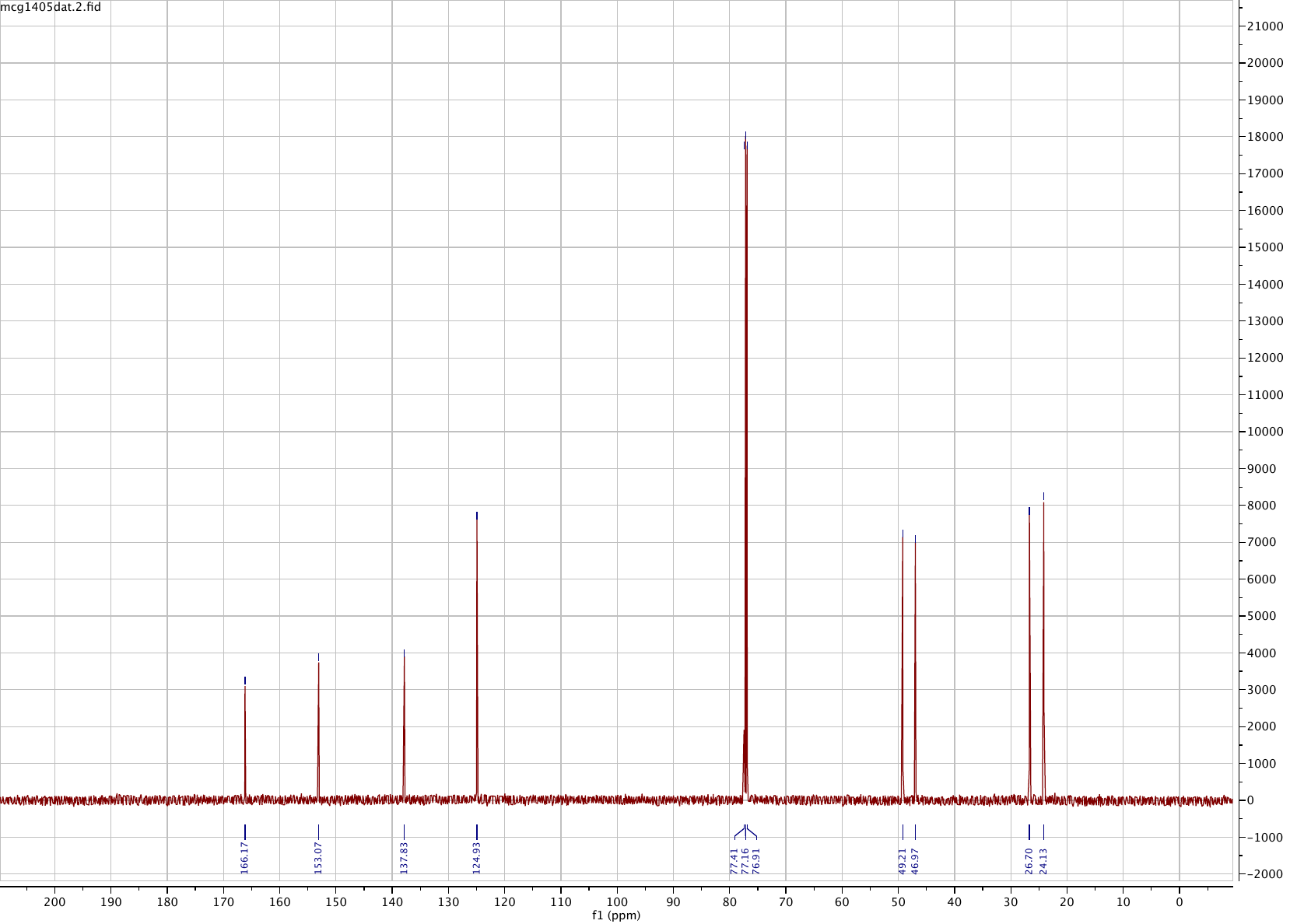

**Compound 7**

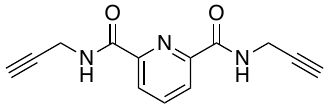

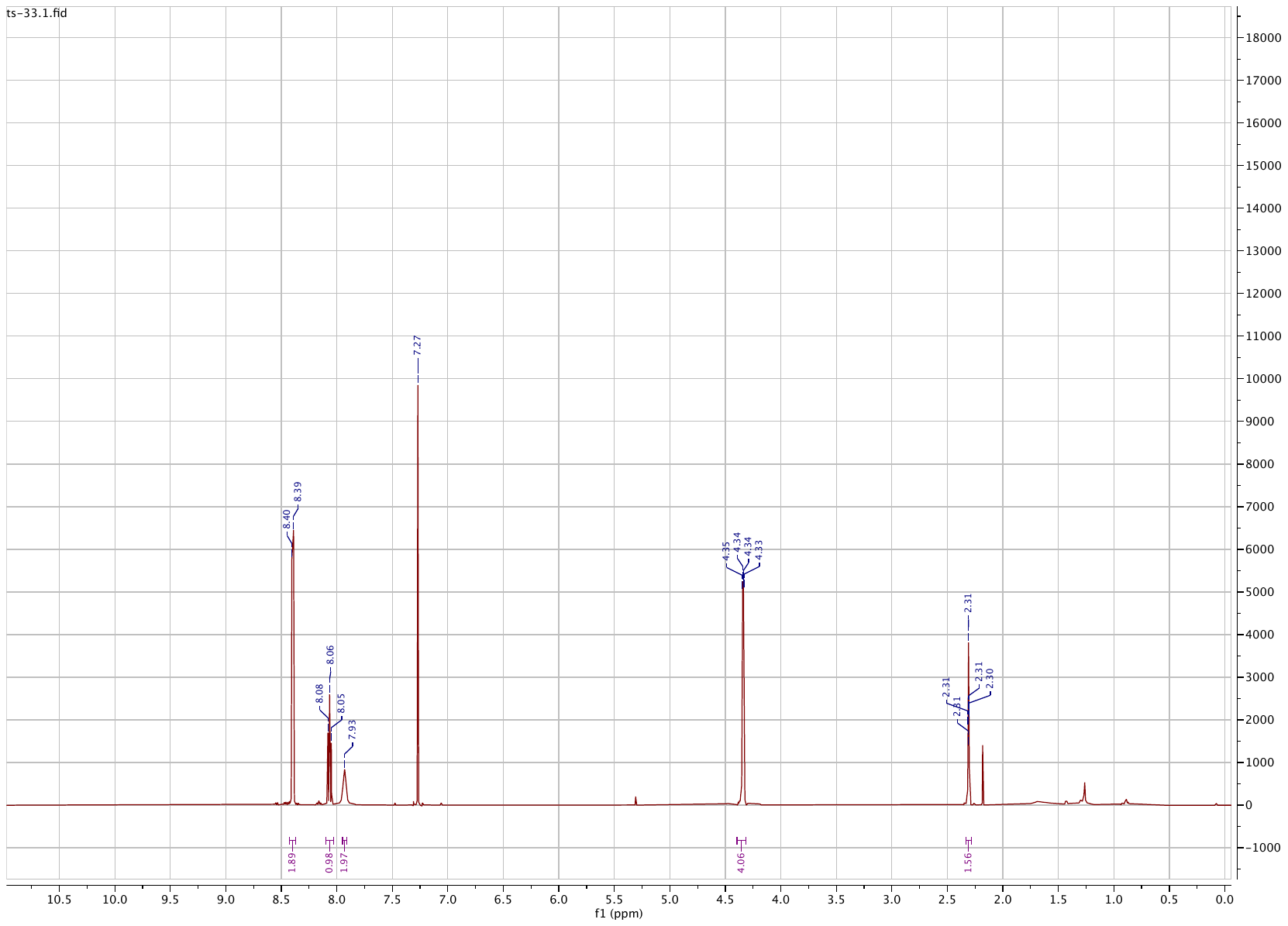

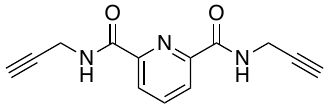

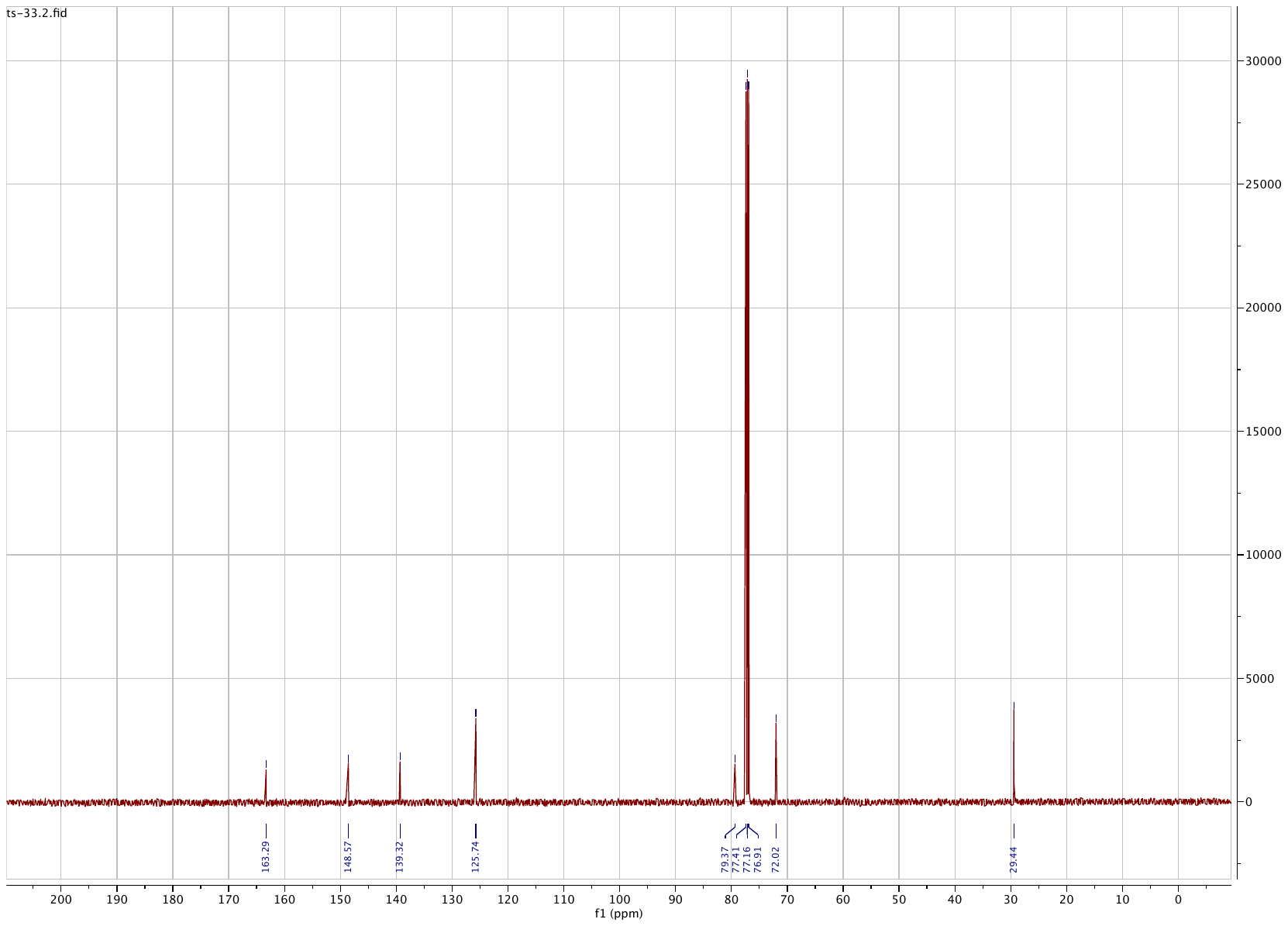

**Compound 8**

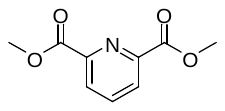

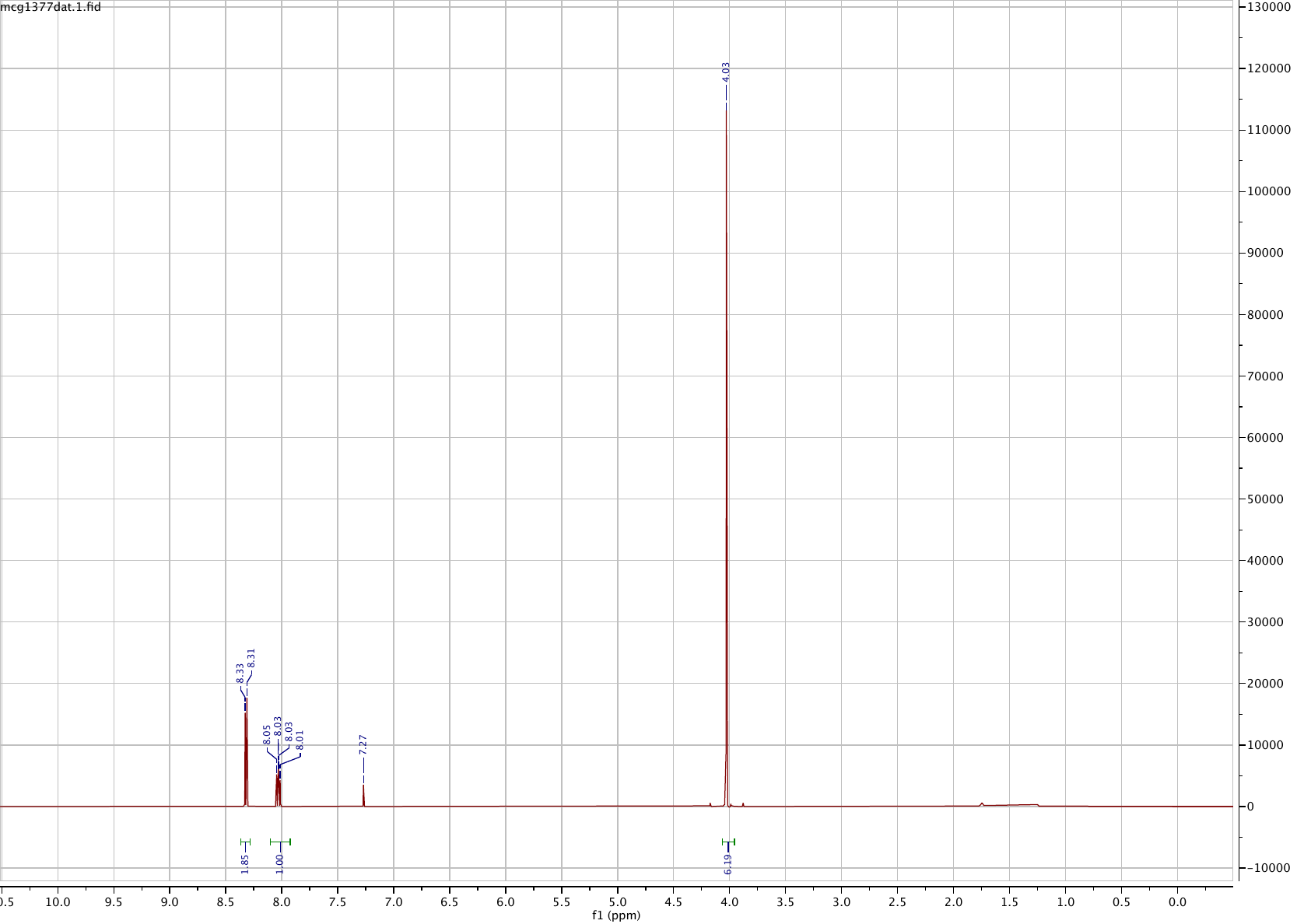

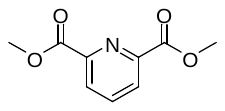

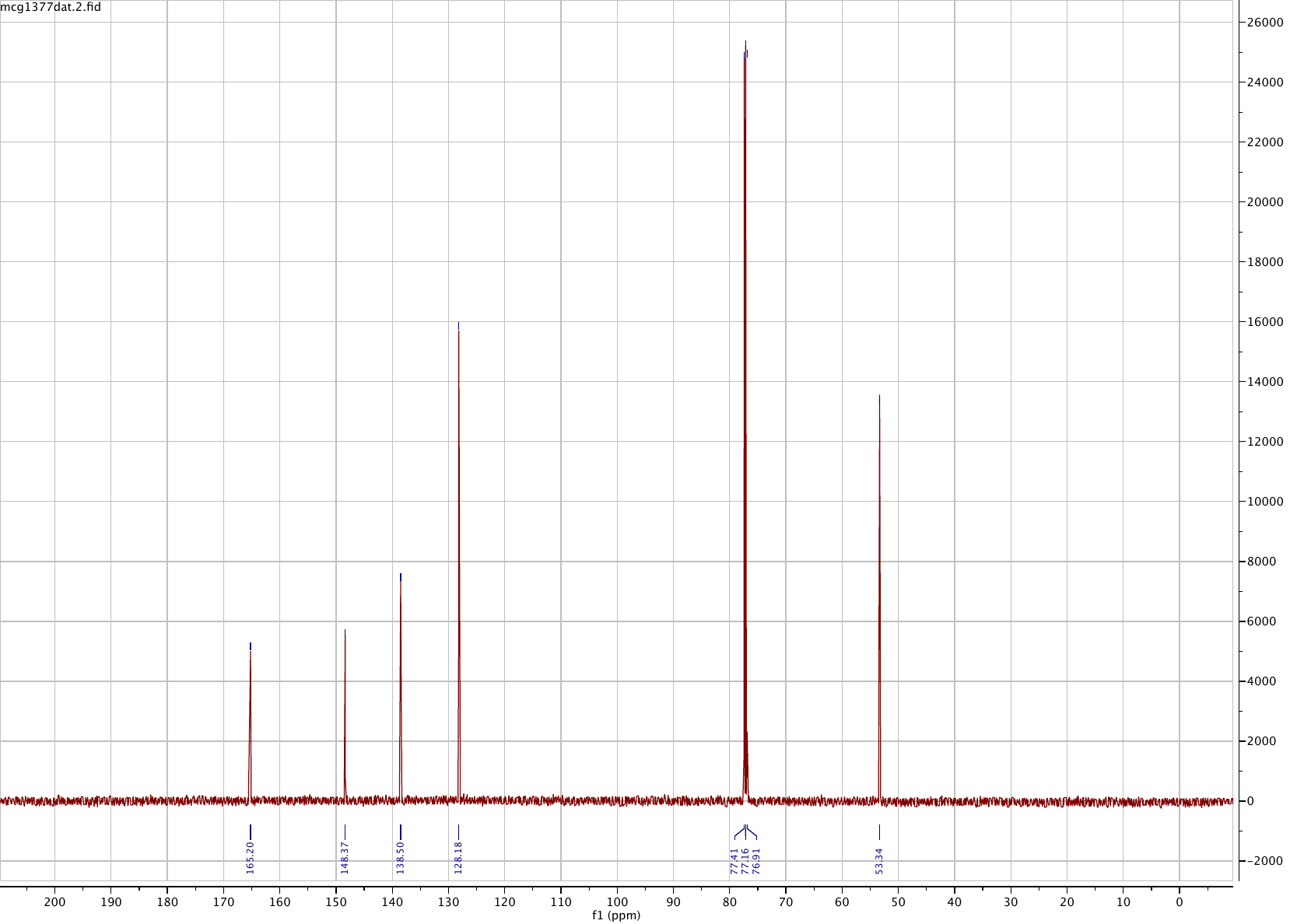

**Compound 9**

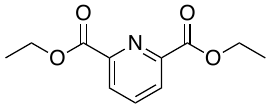

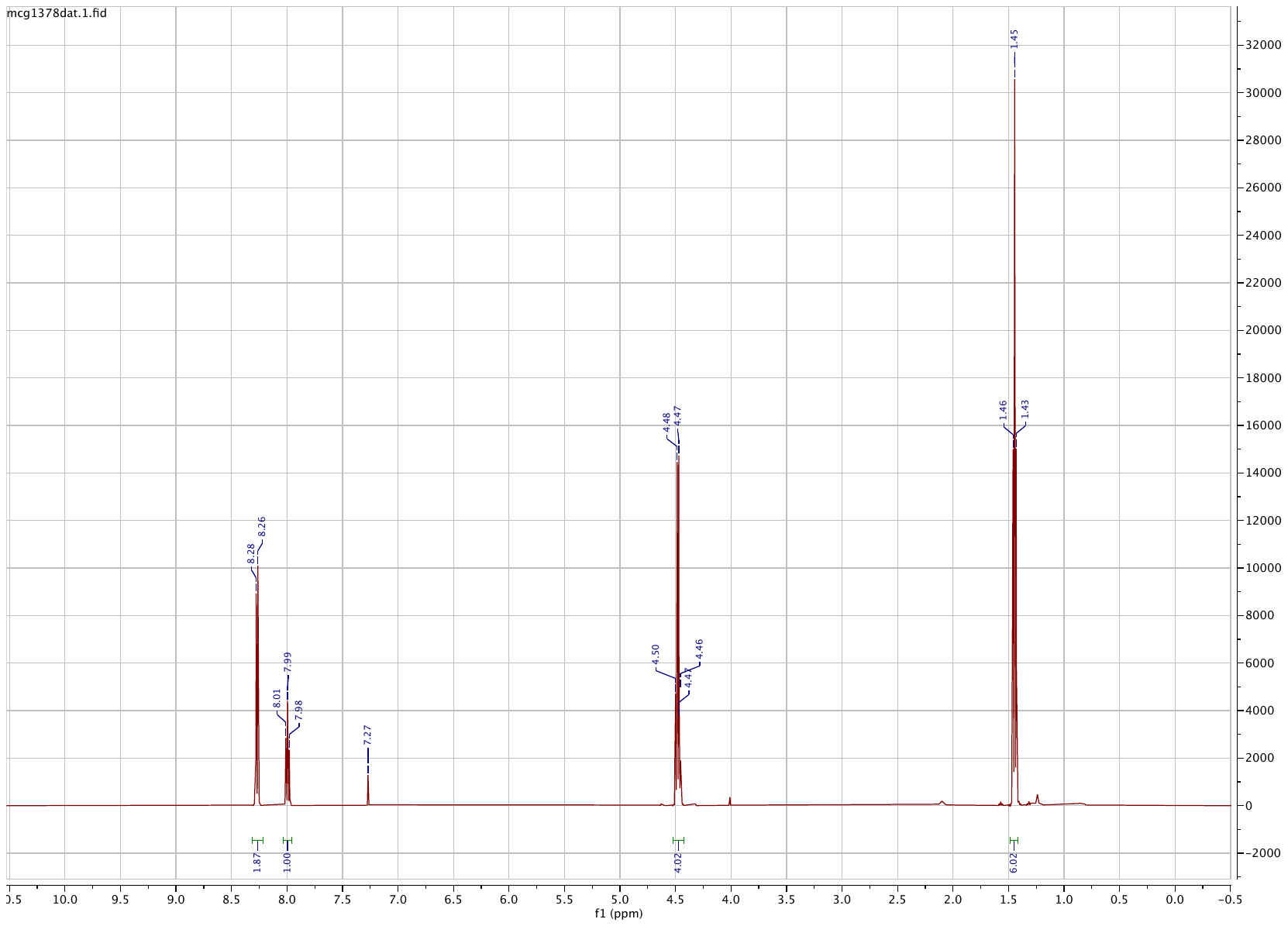

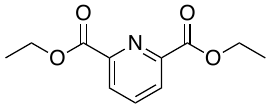

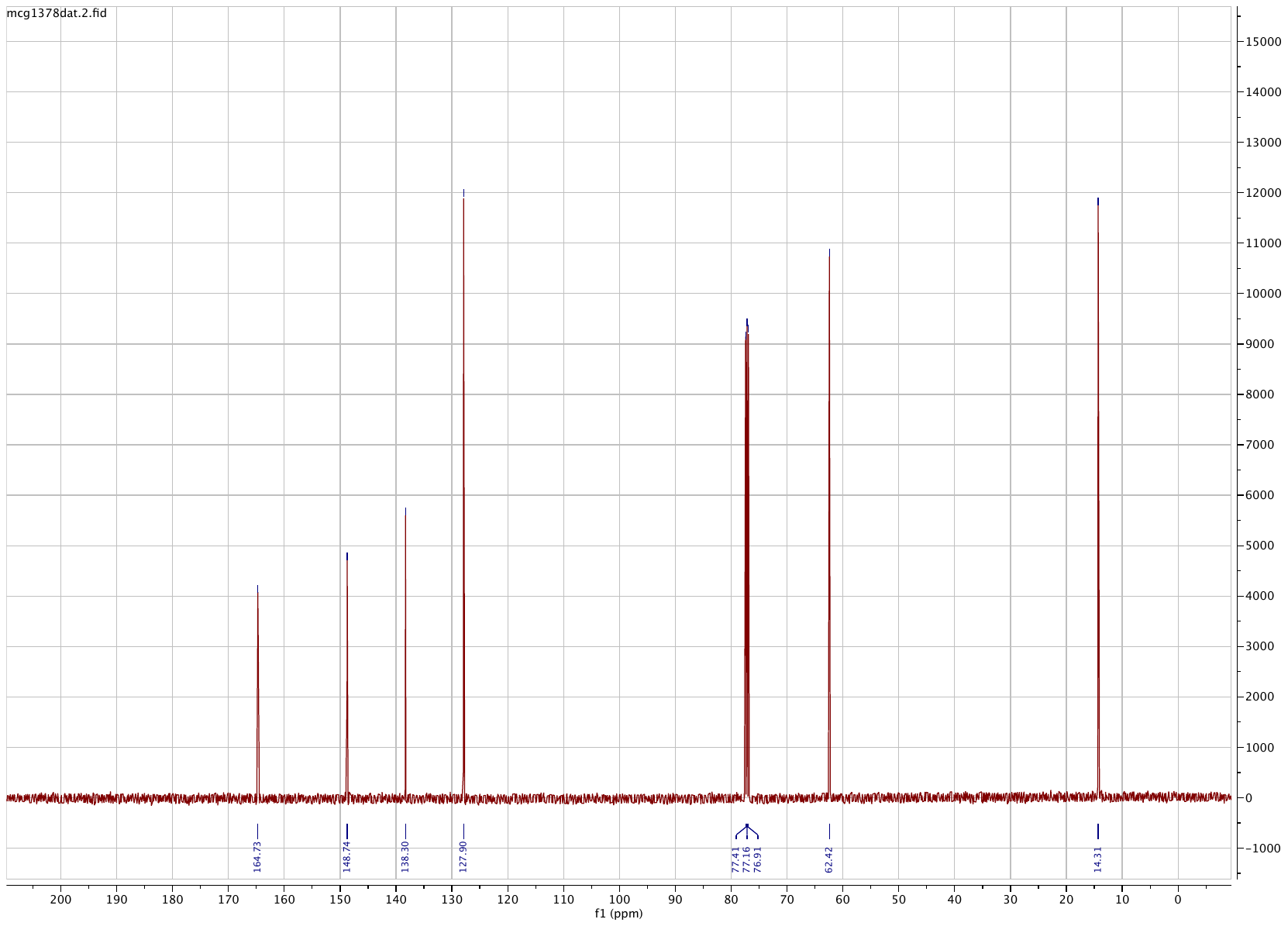

**Compound 10**

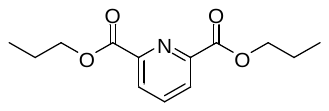

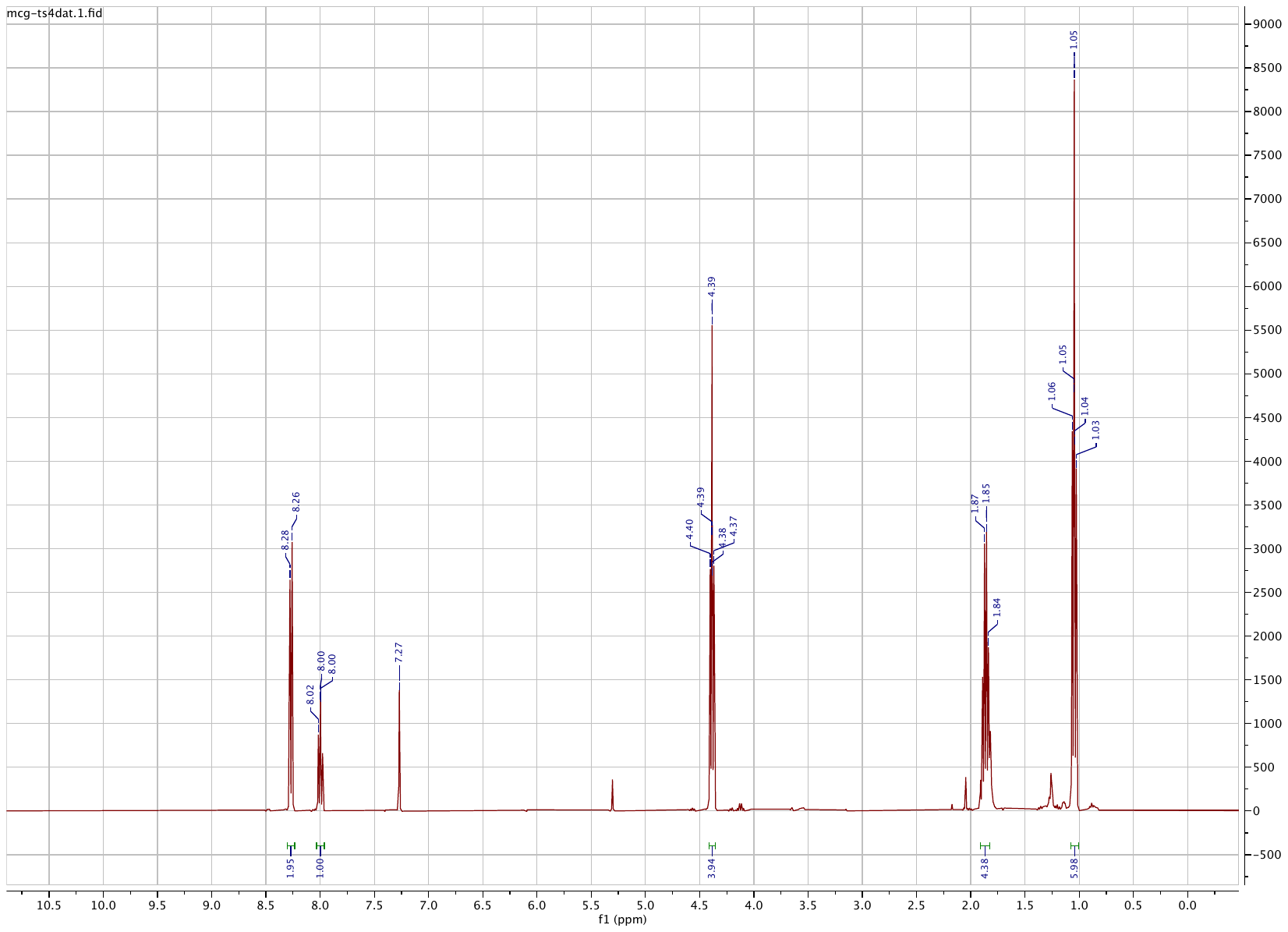

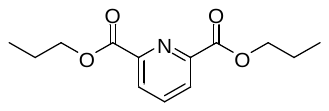

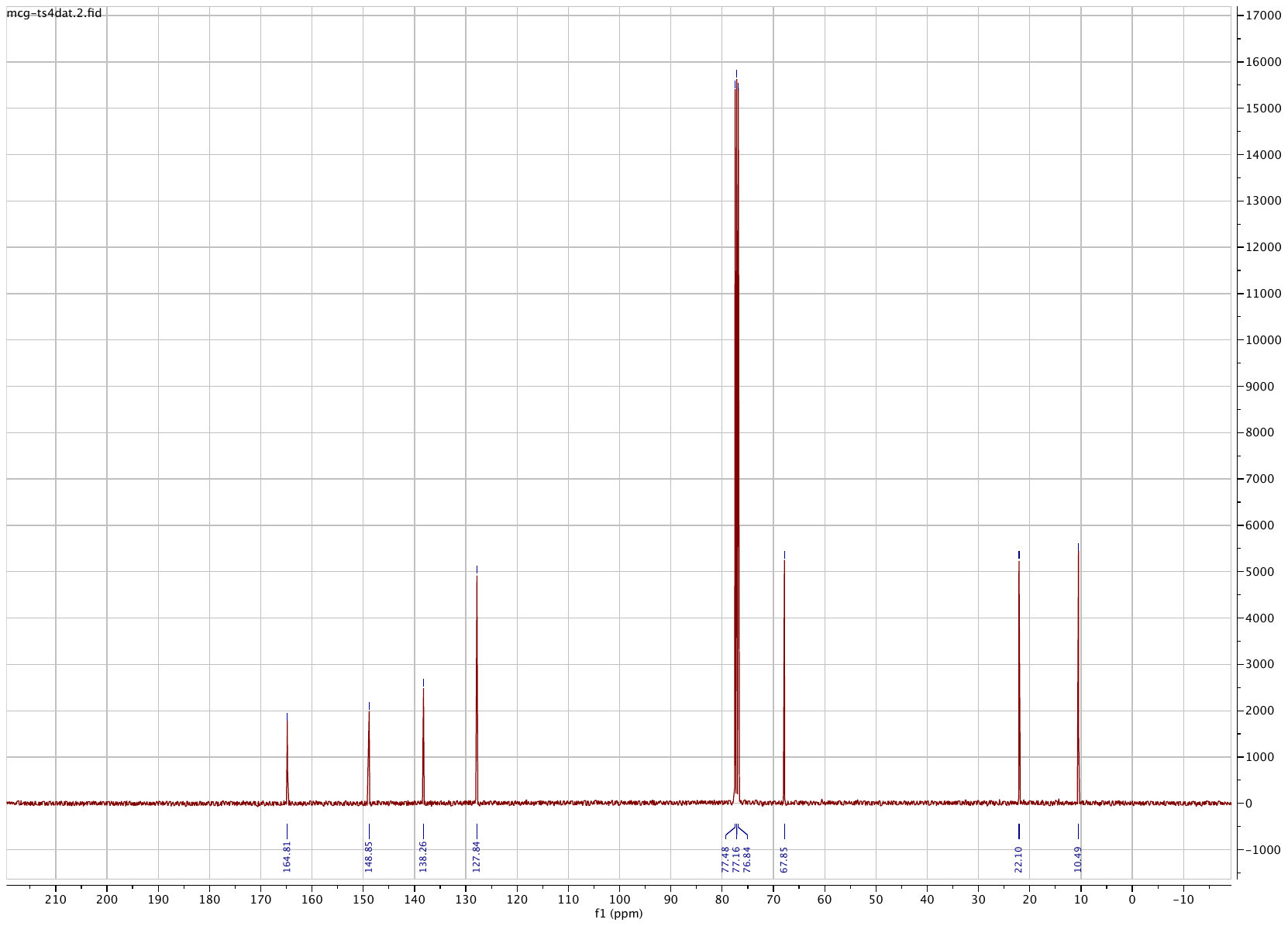

**Compound 11**

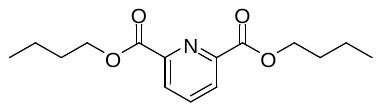

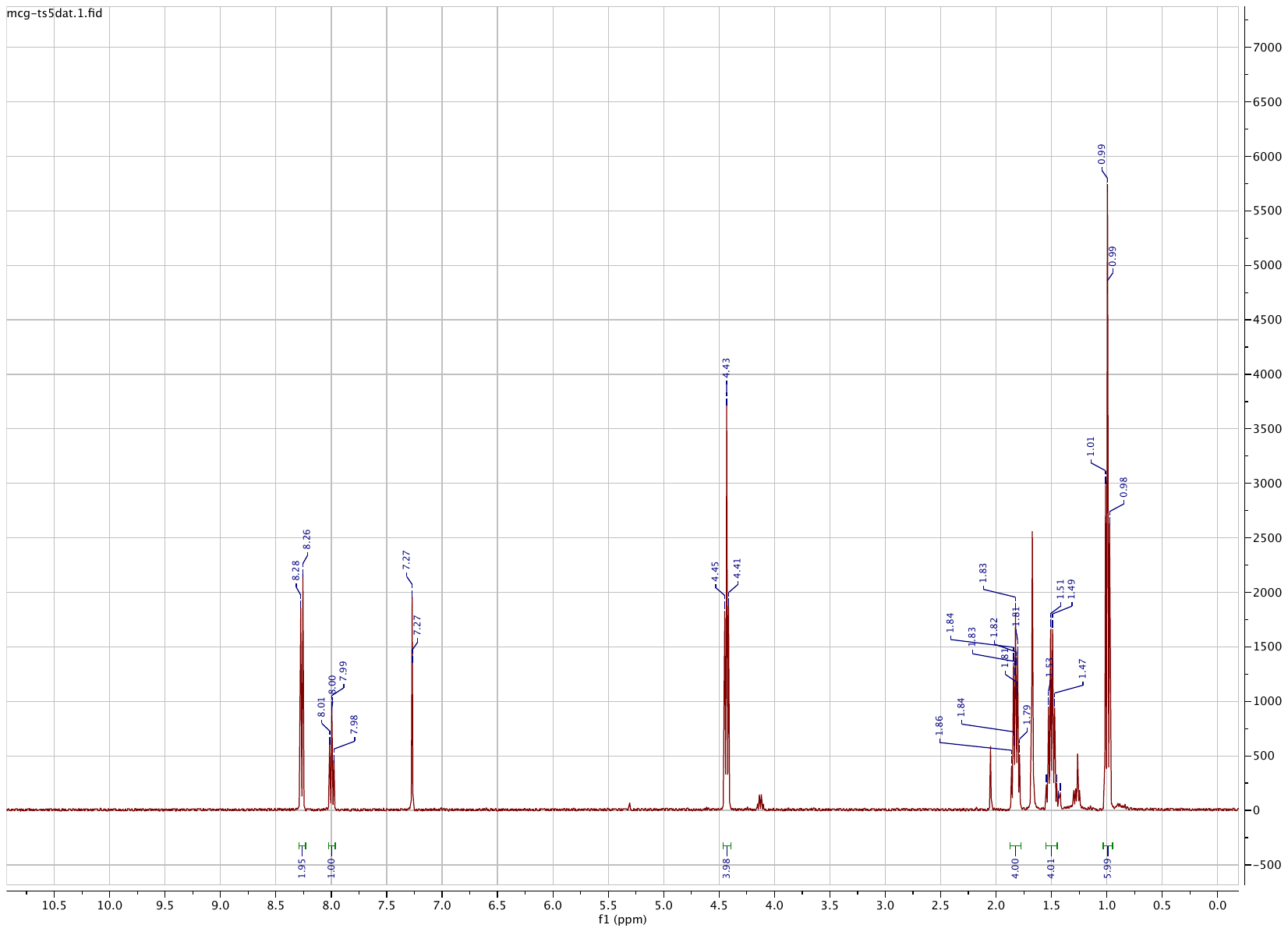

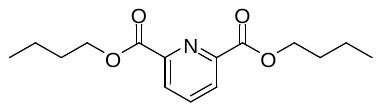

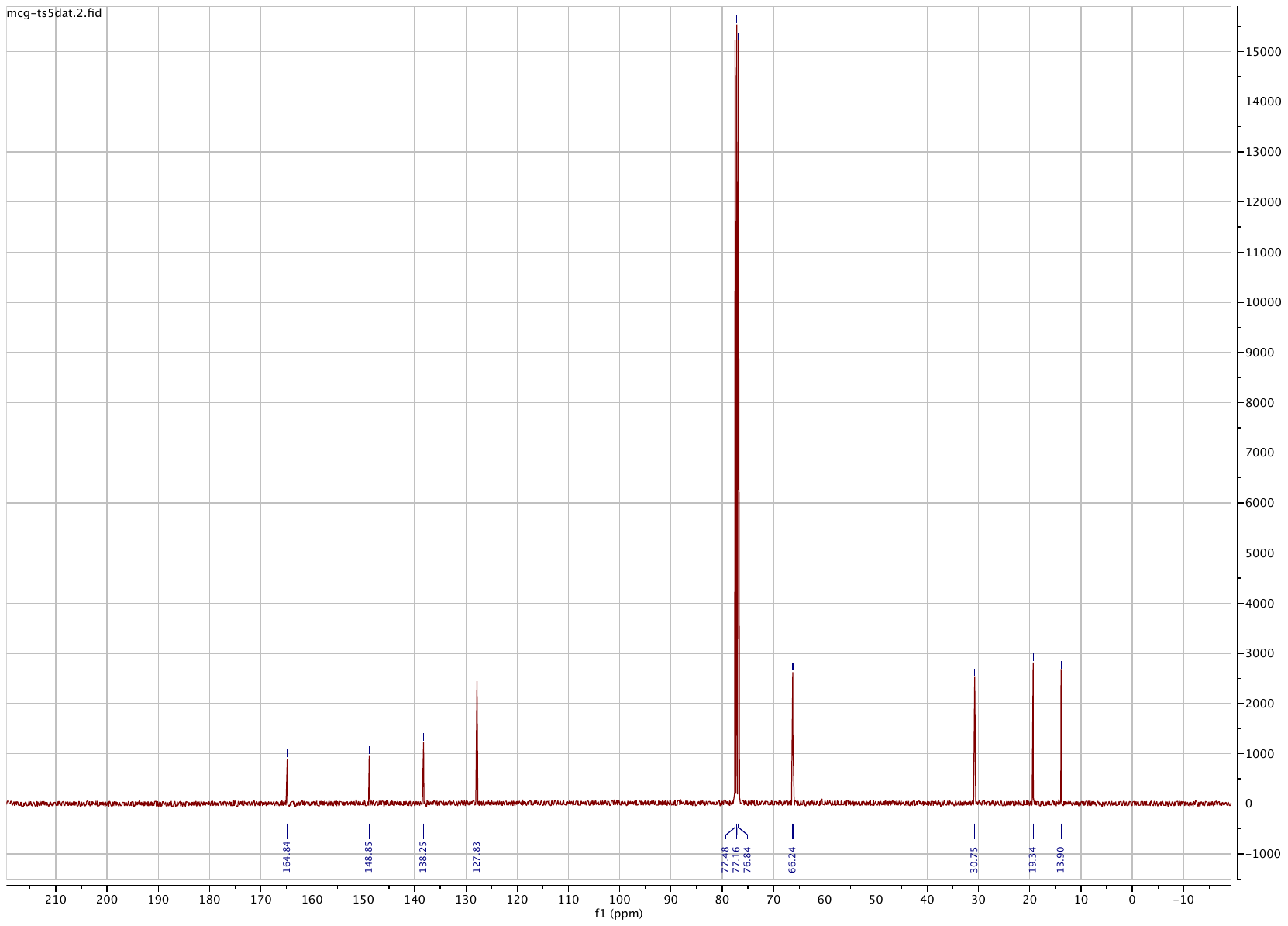

**Compound 12**

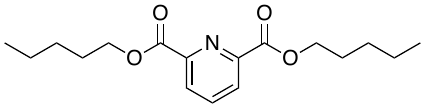

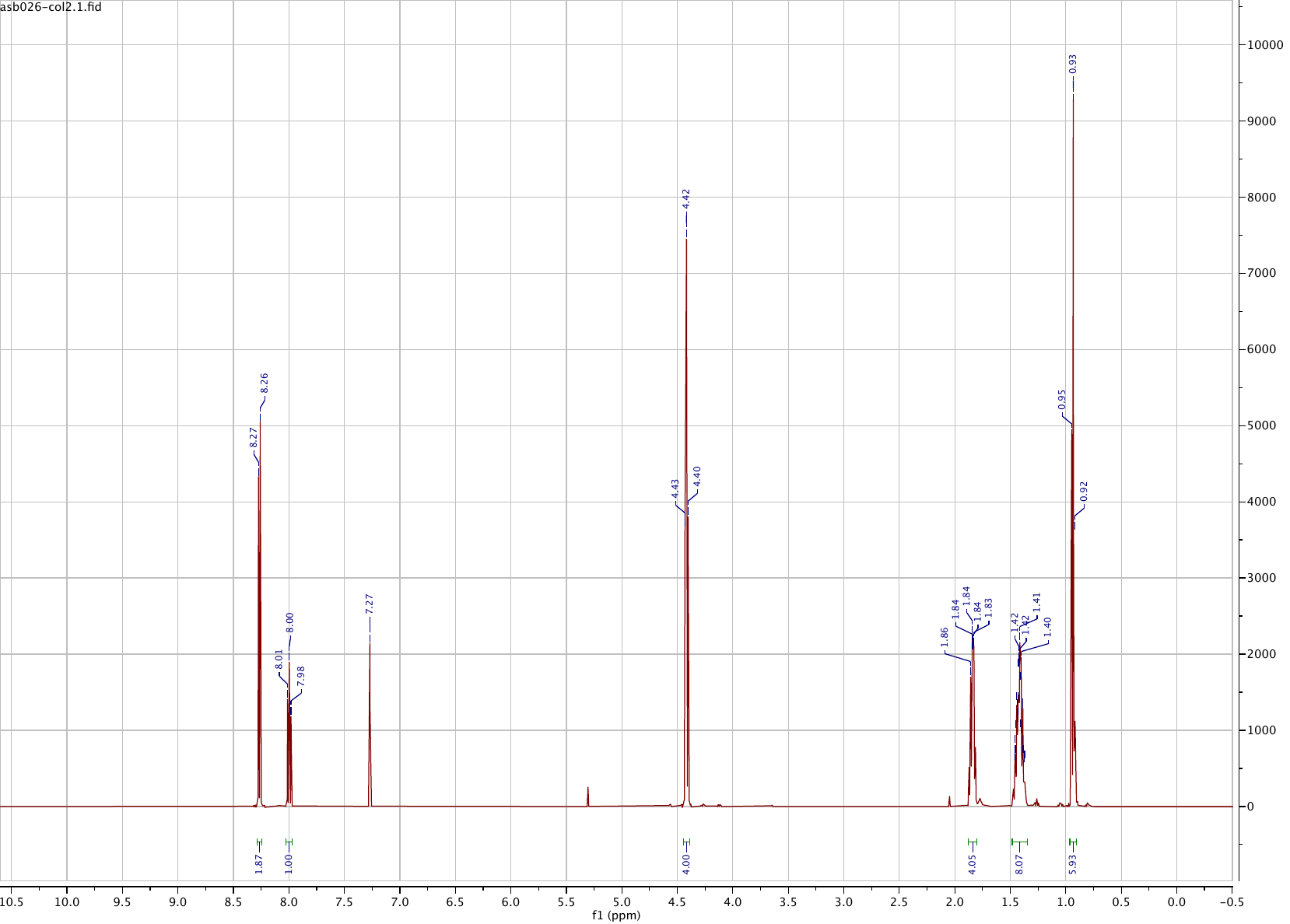

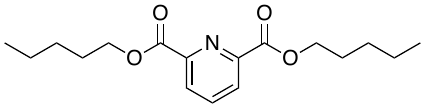

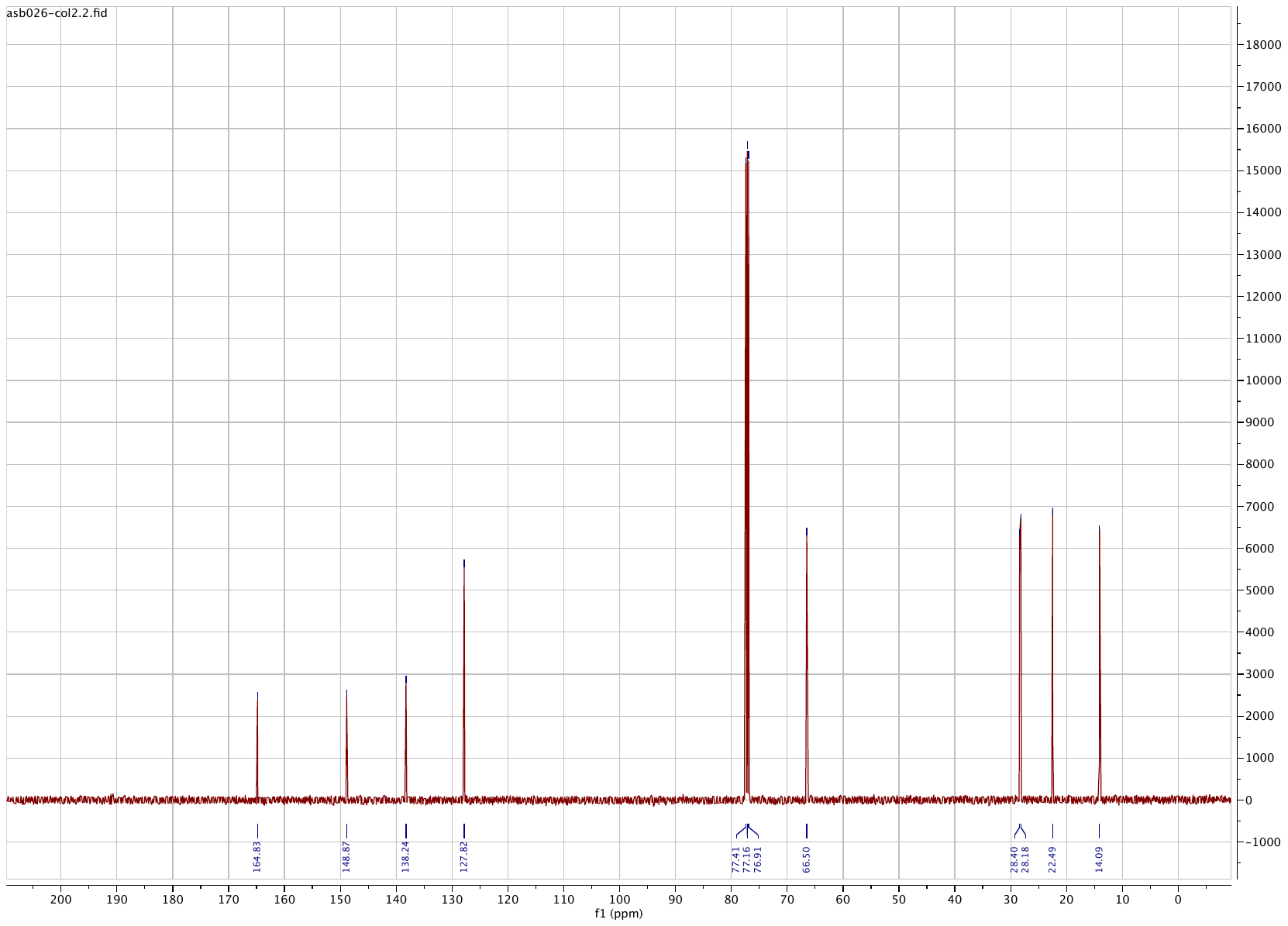

**Compound 13**

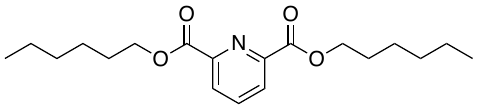

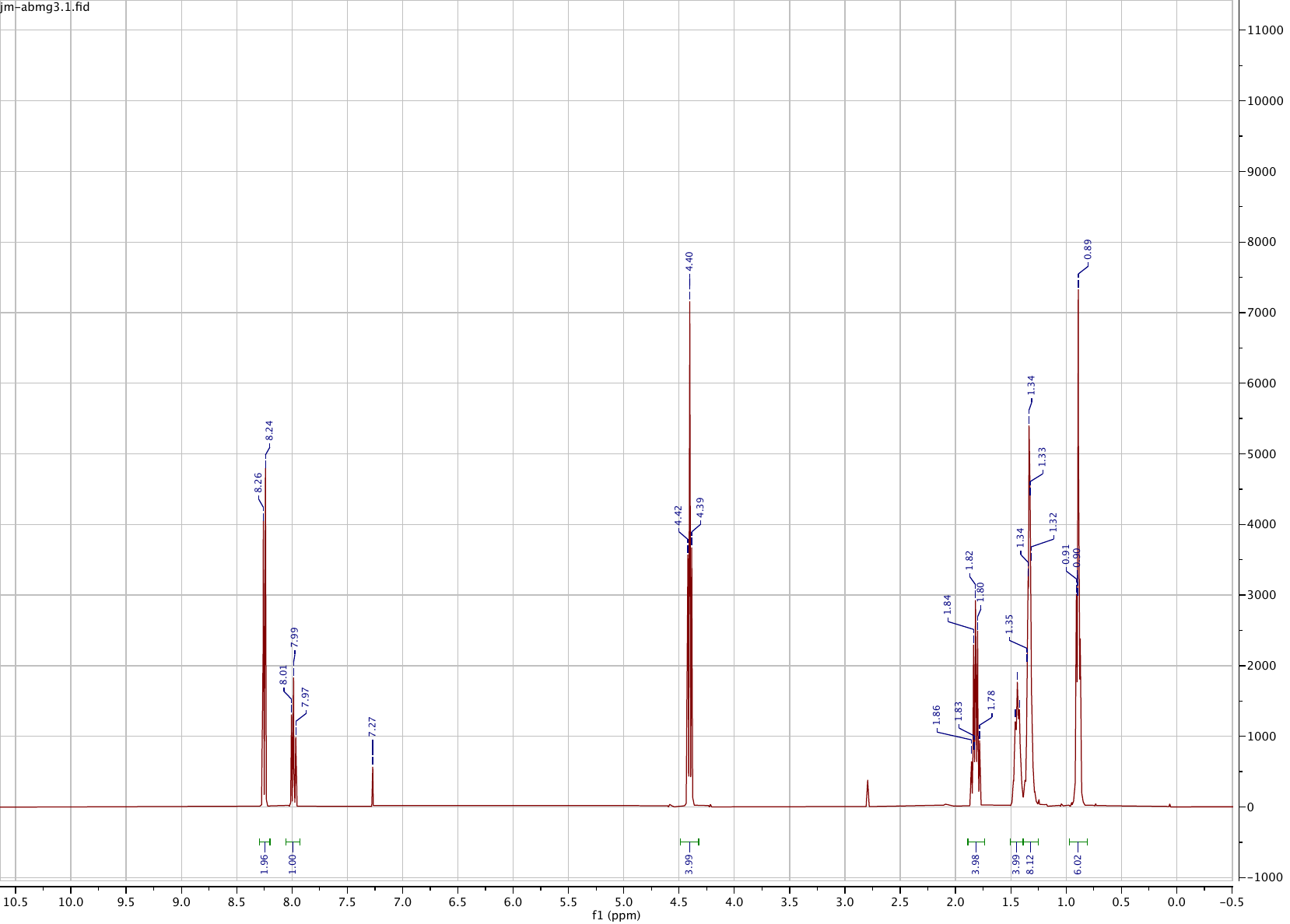

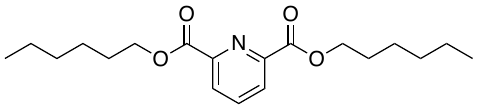

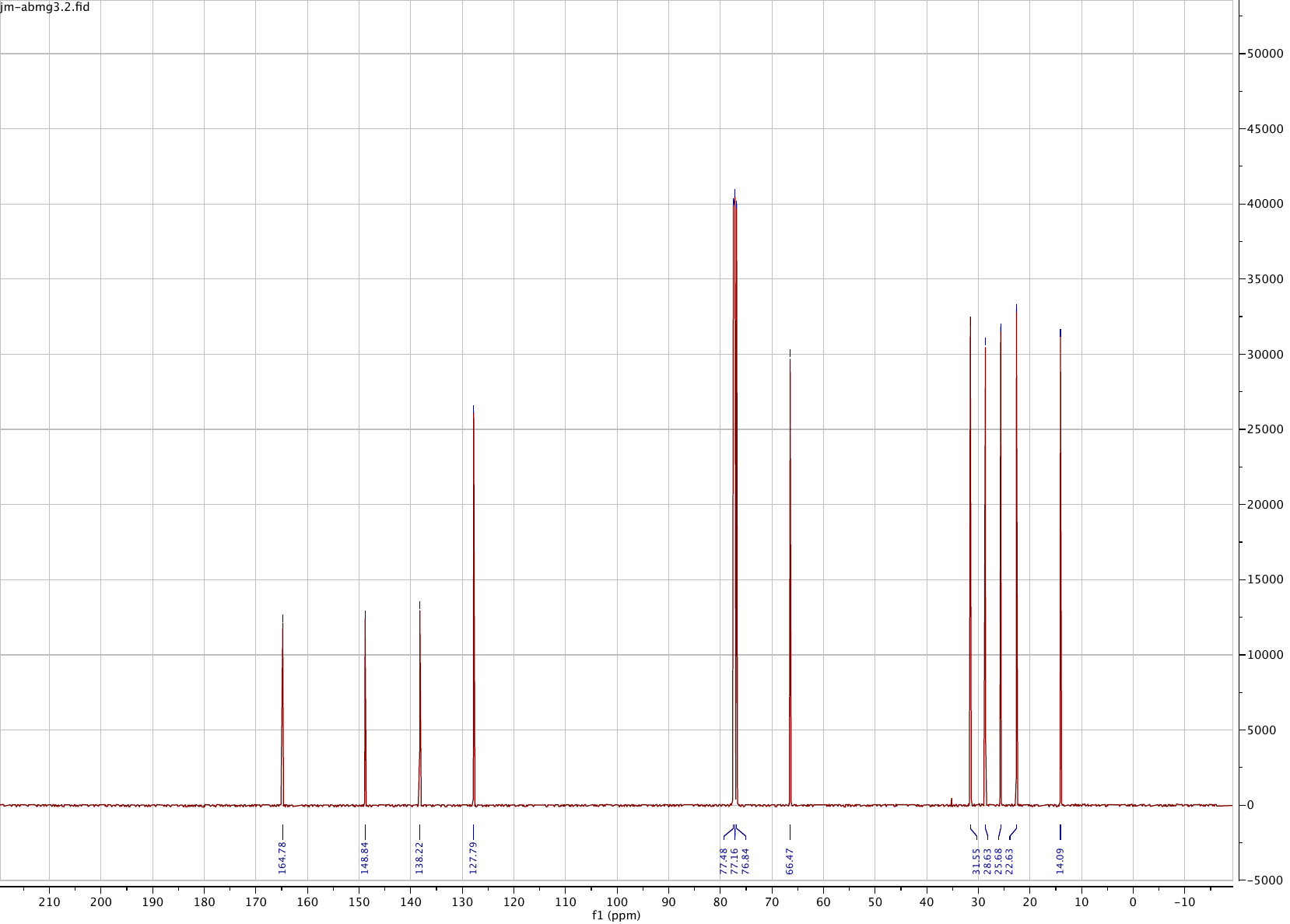

**Compound 14**

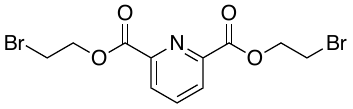

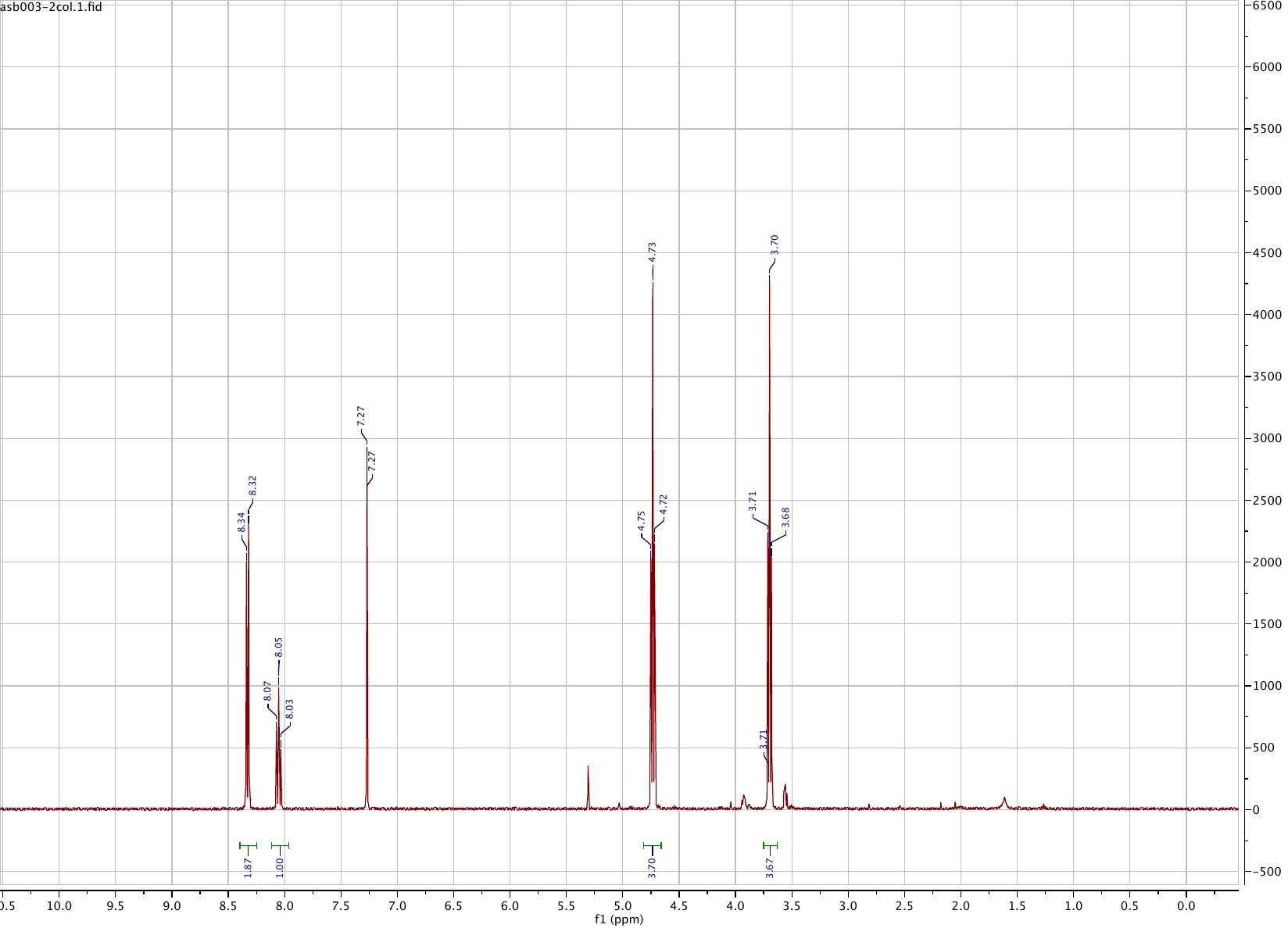

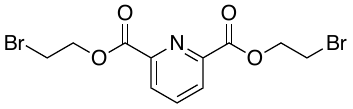

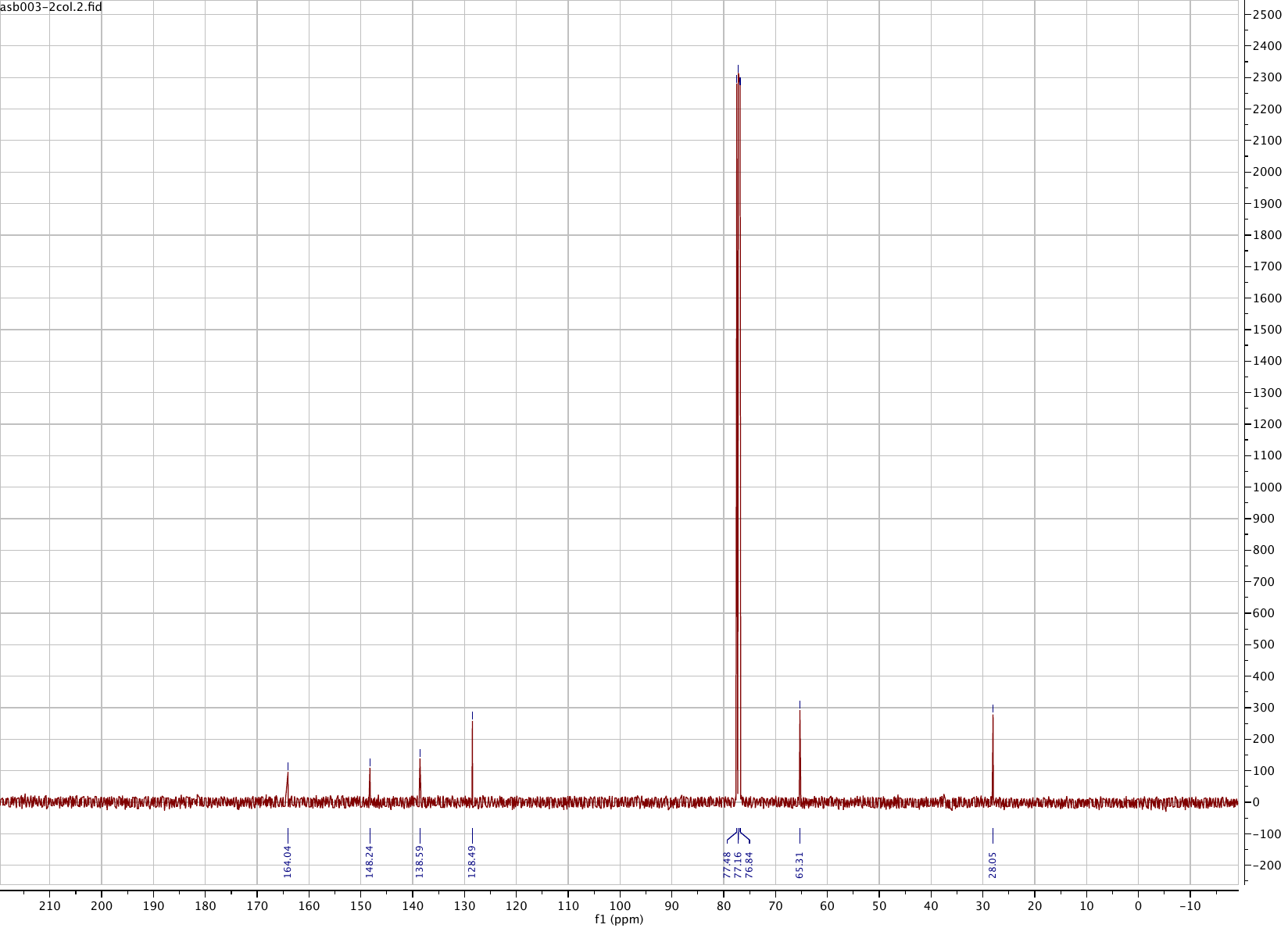

**Compound 15**

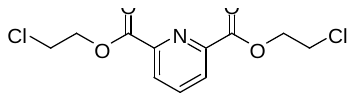

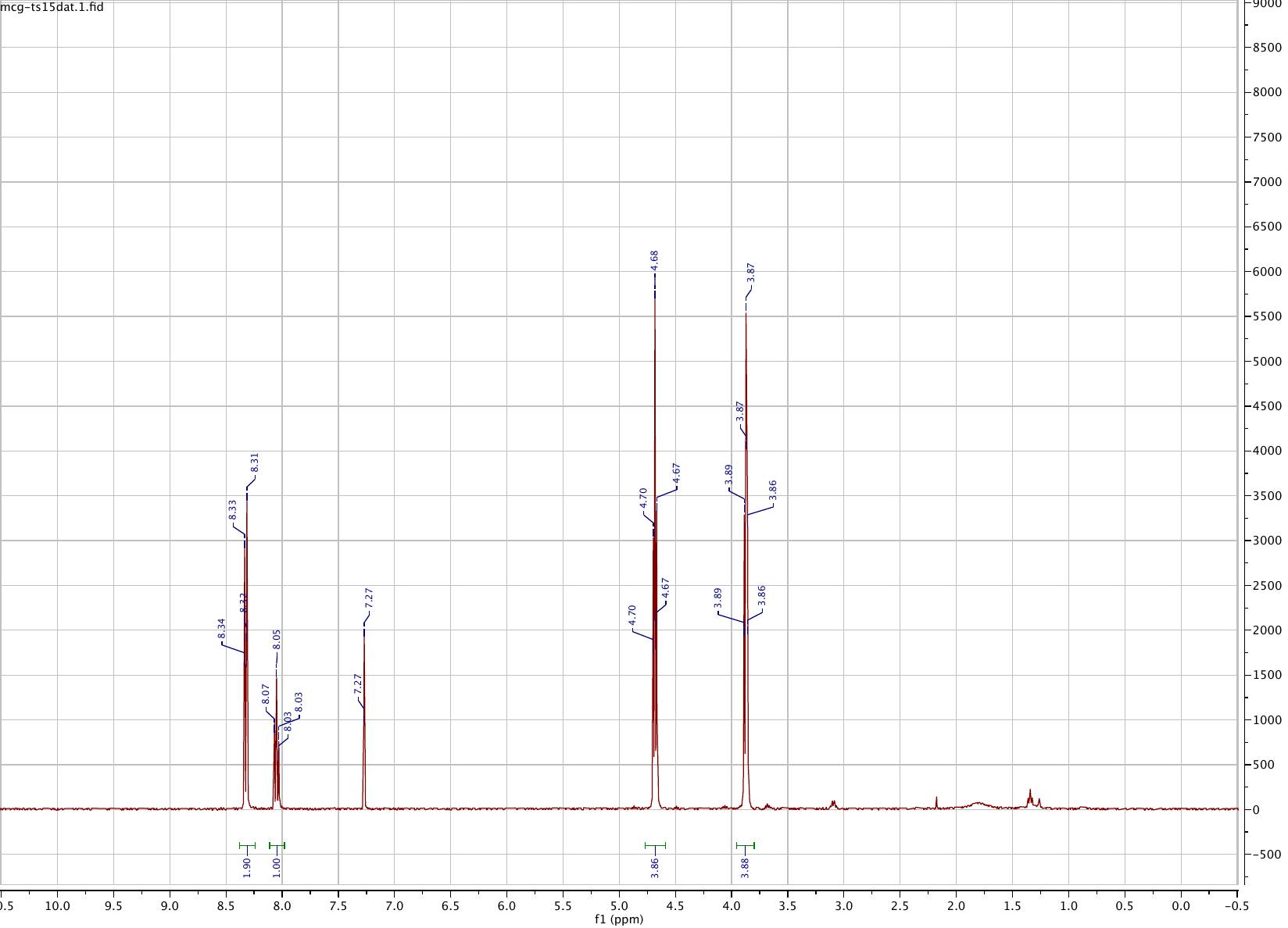

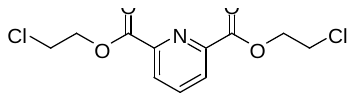

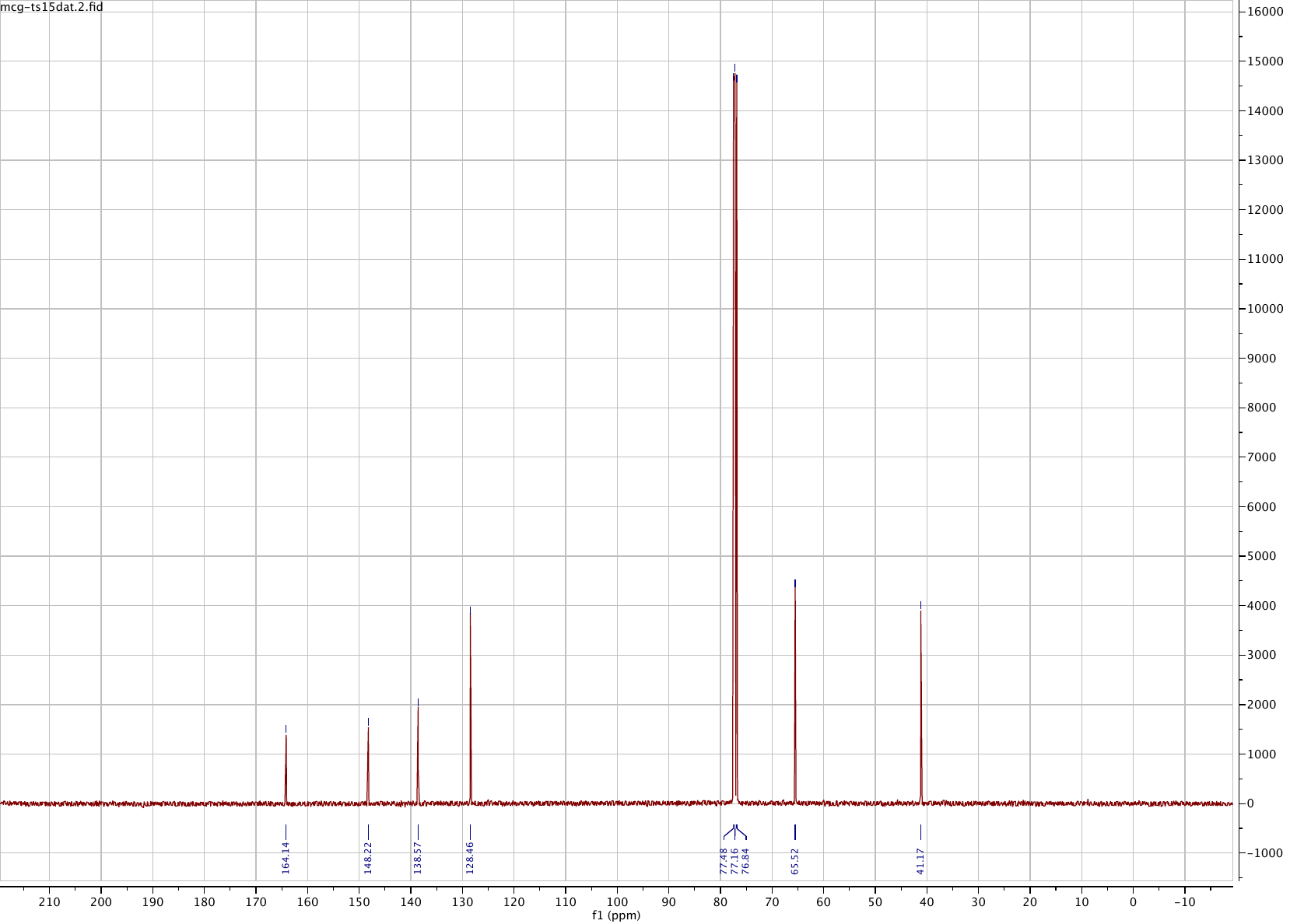

**Compound 16**

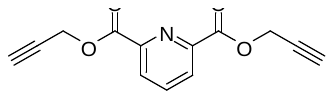

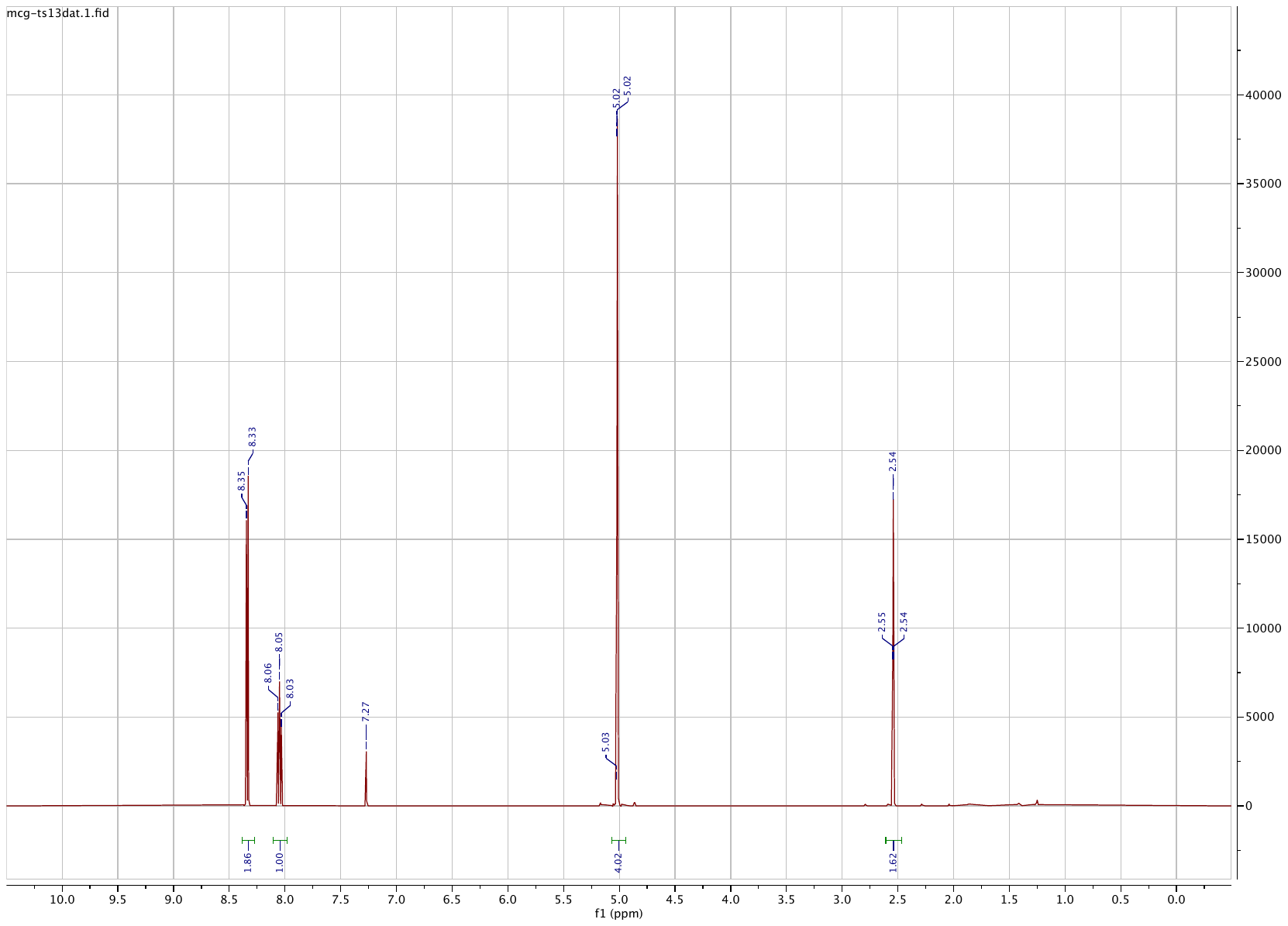

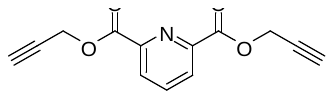

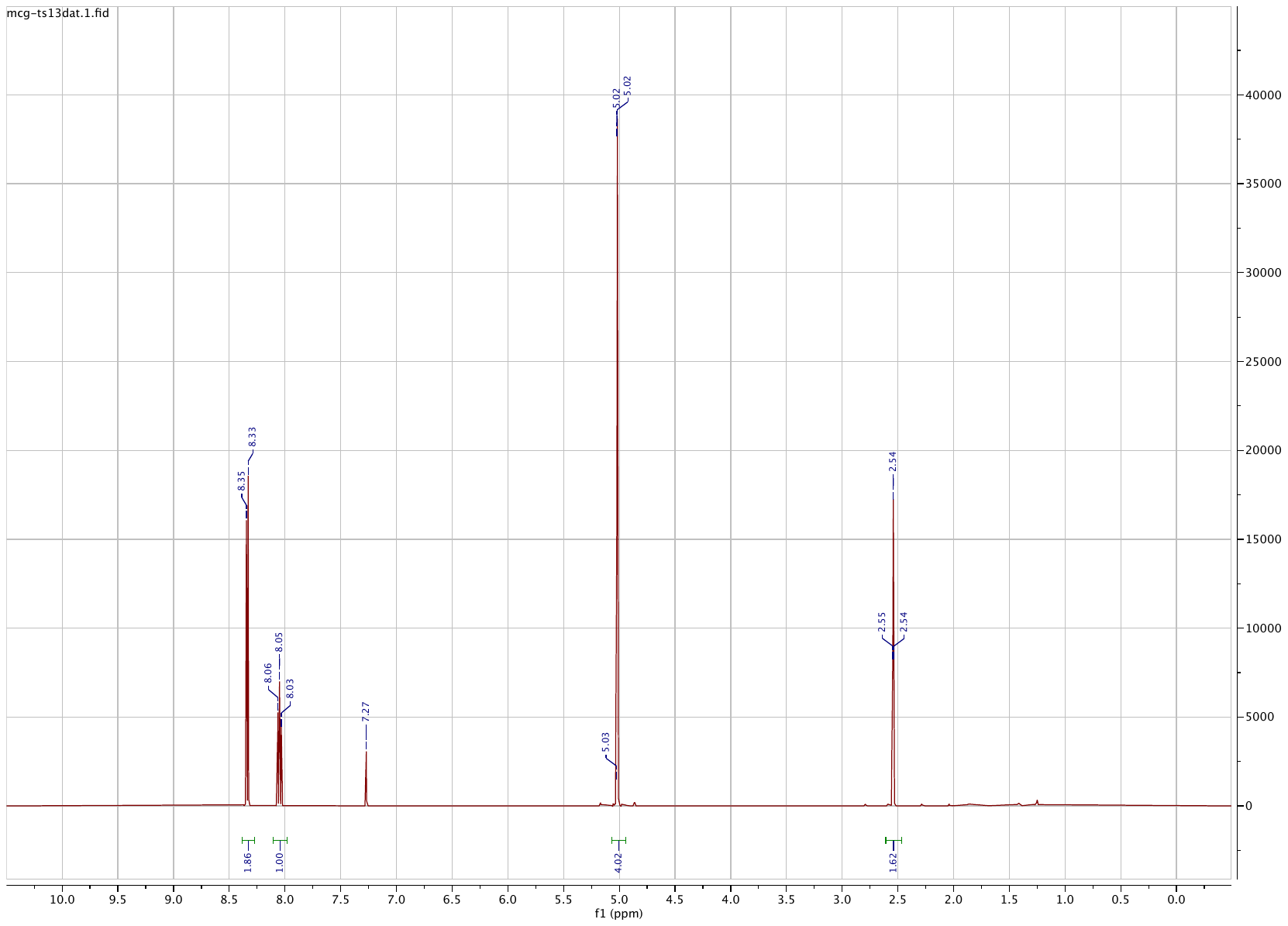

**Compound 17**

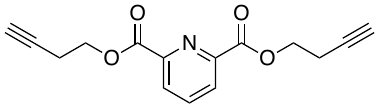

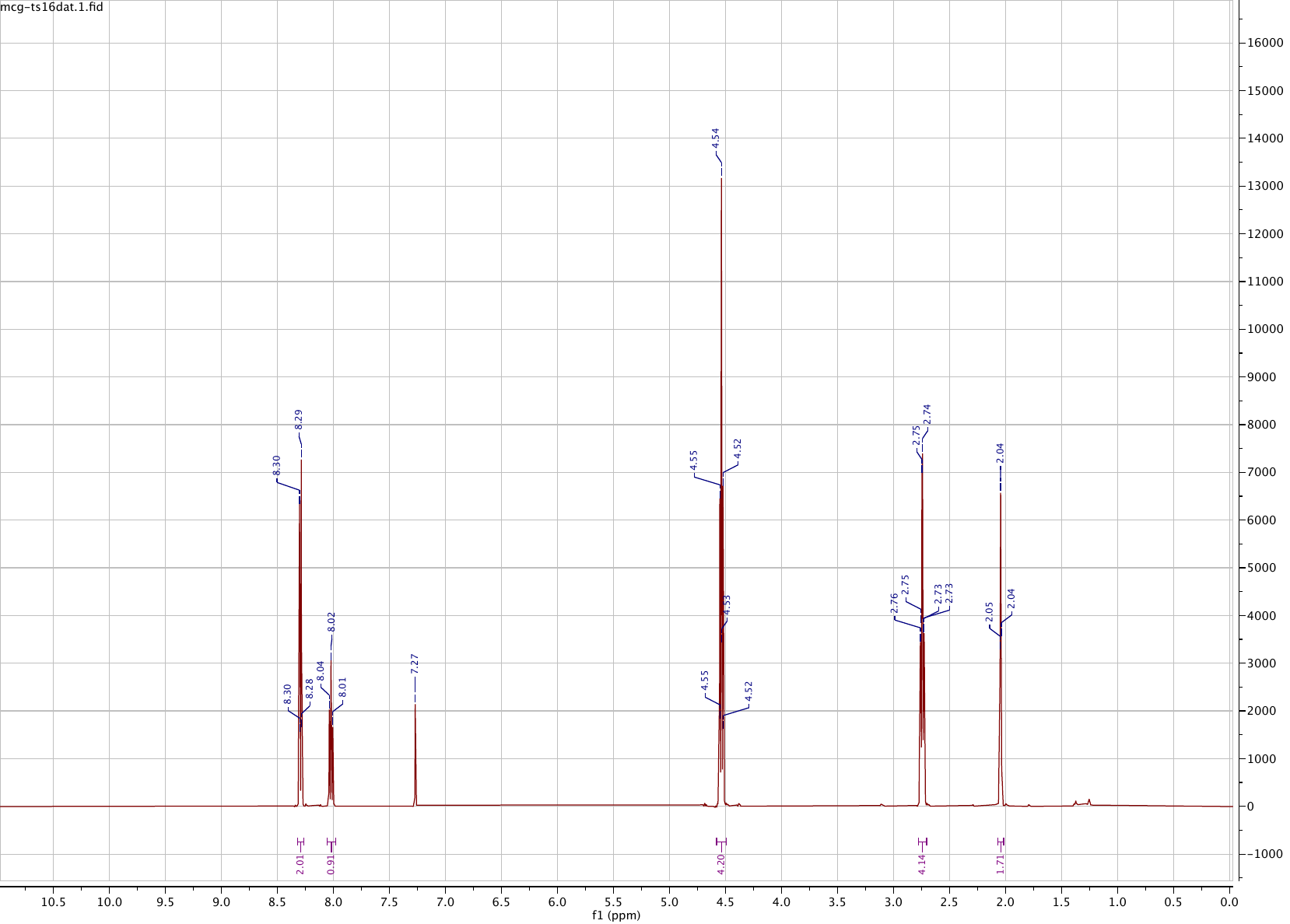

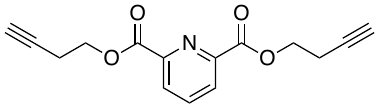

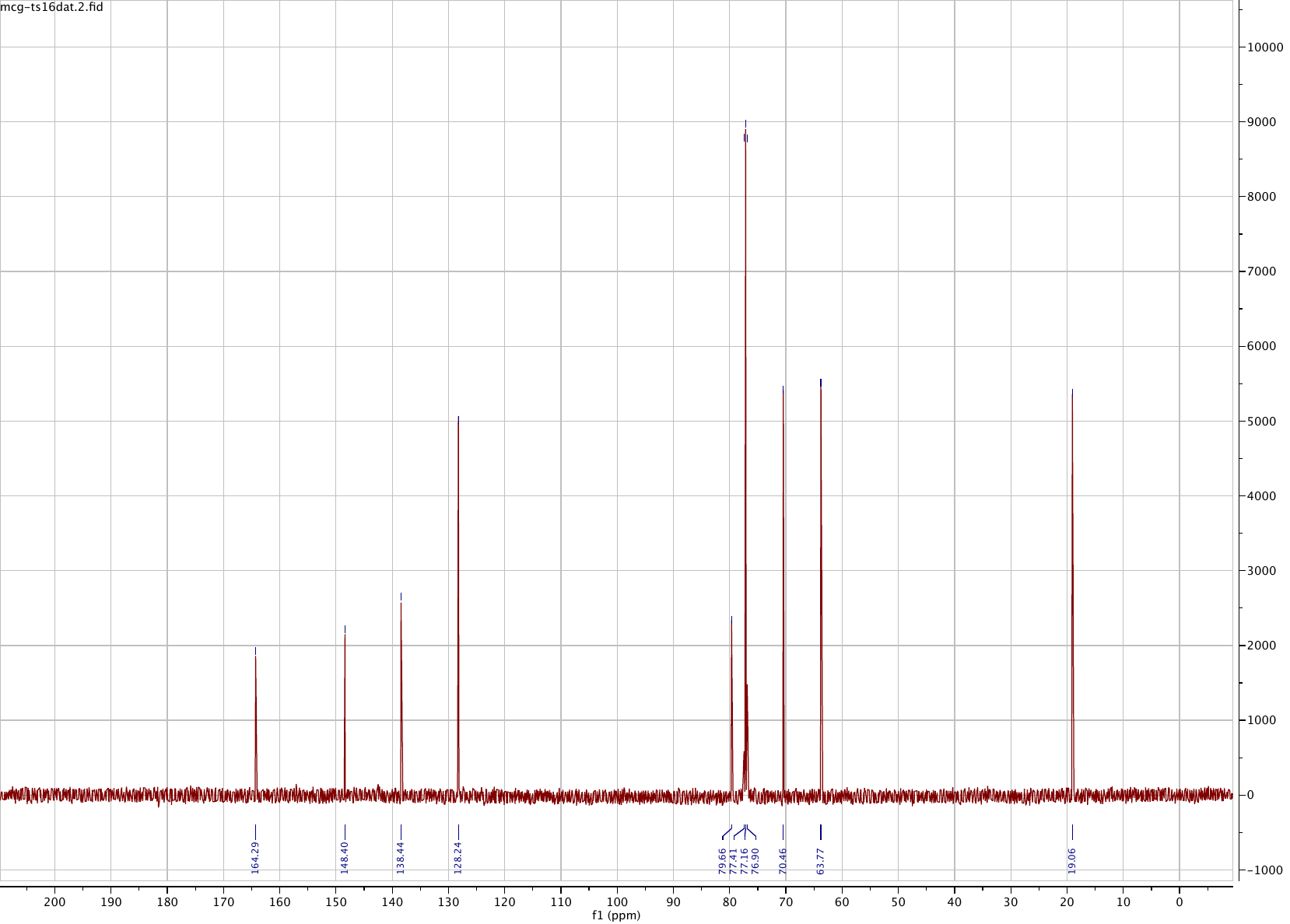

**Compound 18**

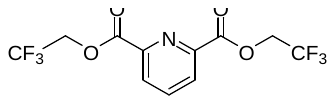

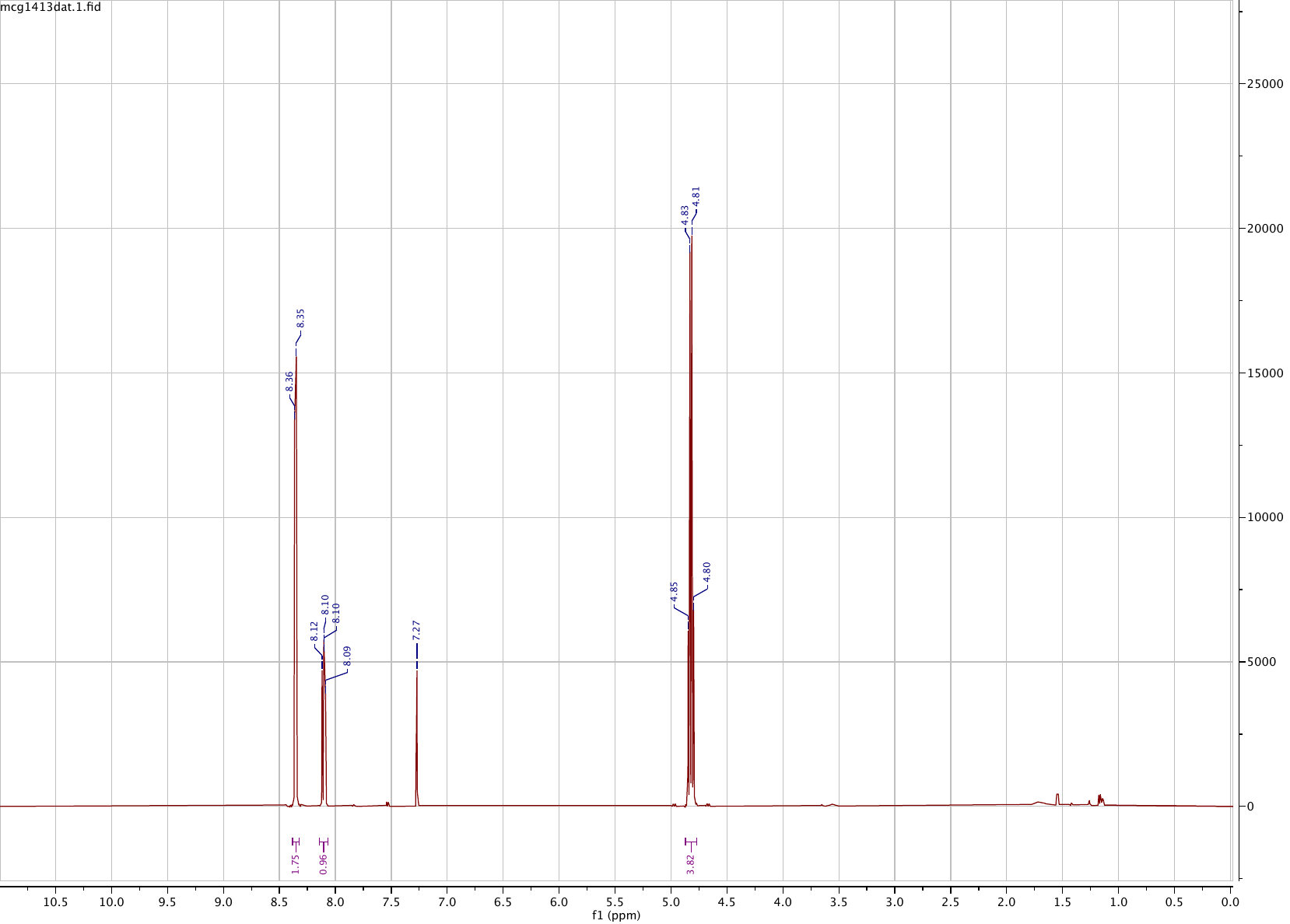

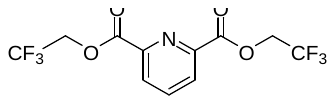

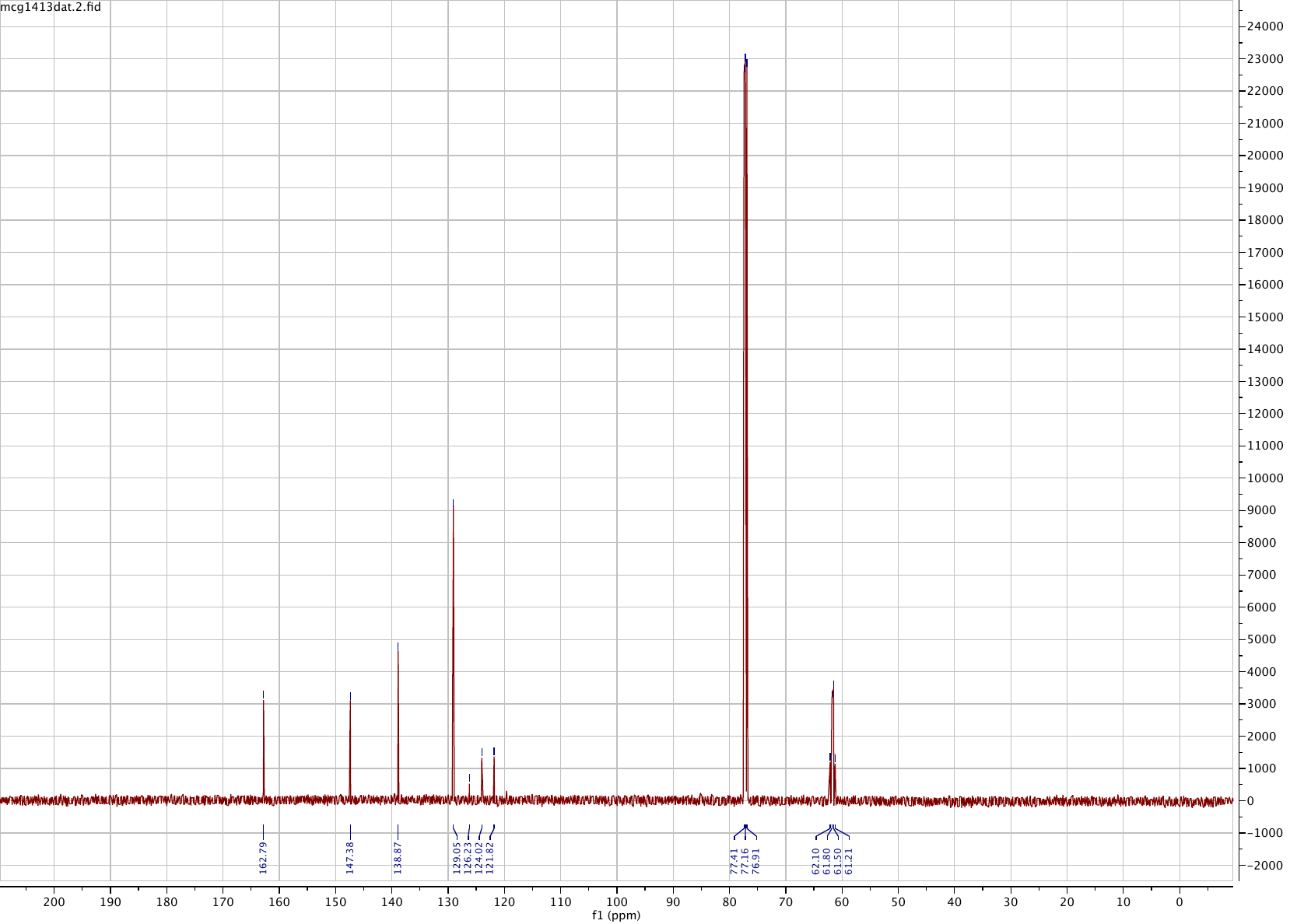

**Compound 19**

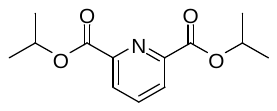

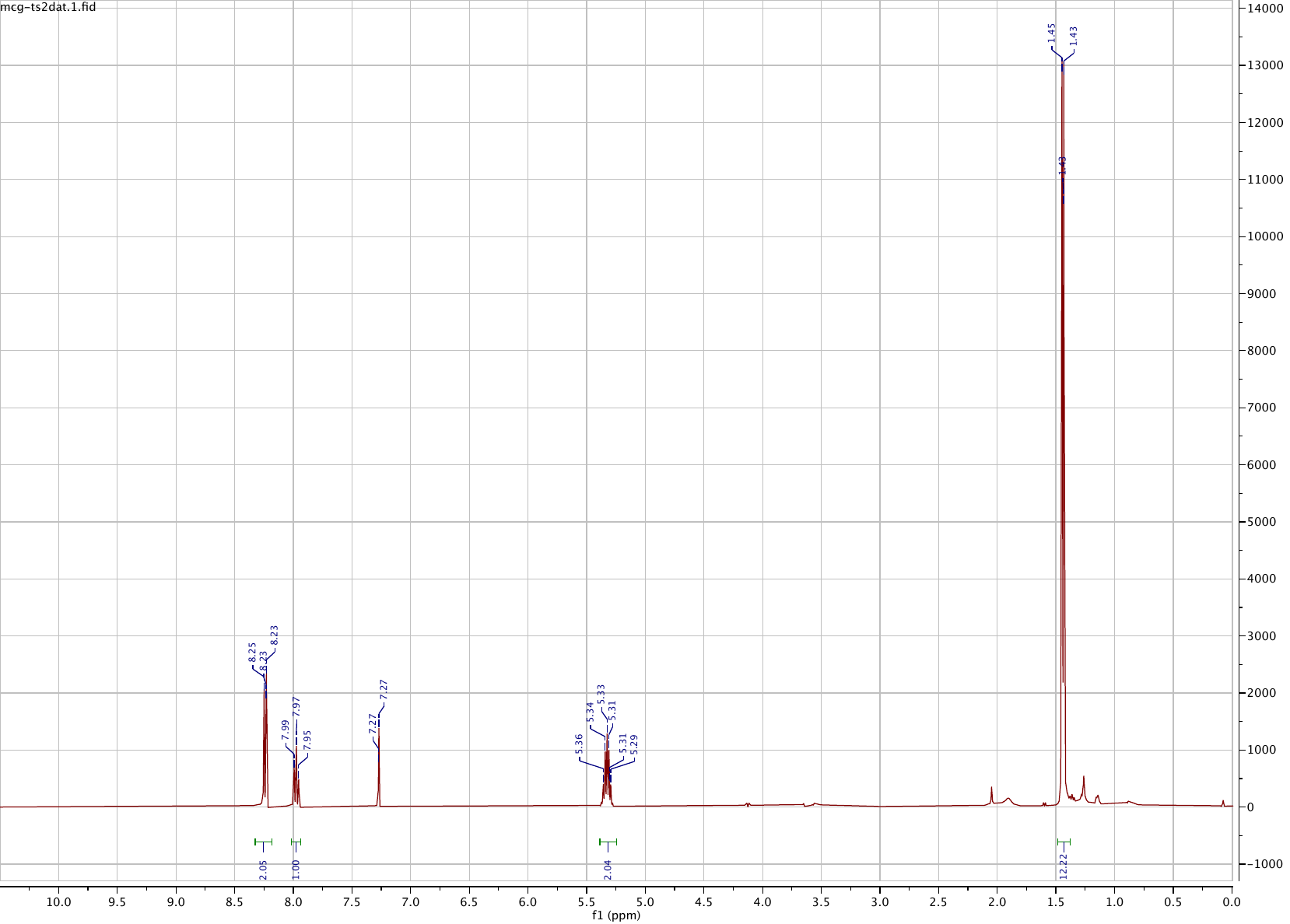

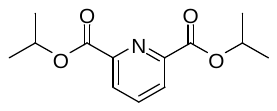

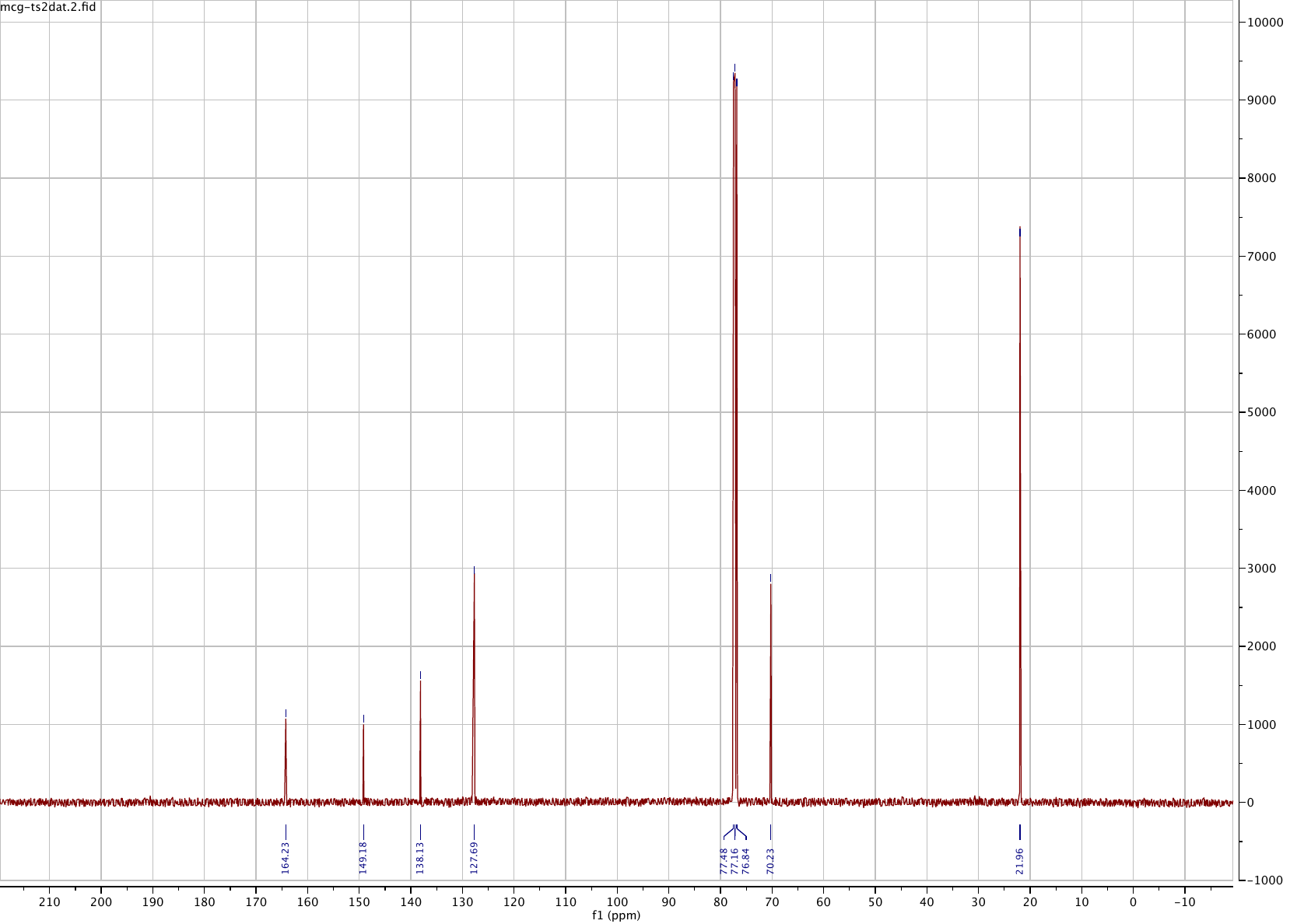

**Compound 20**

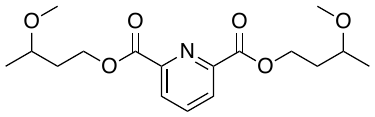

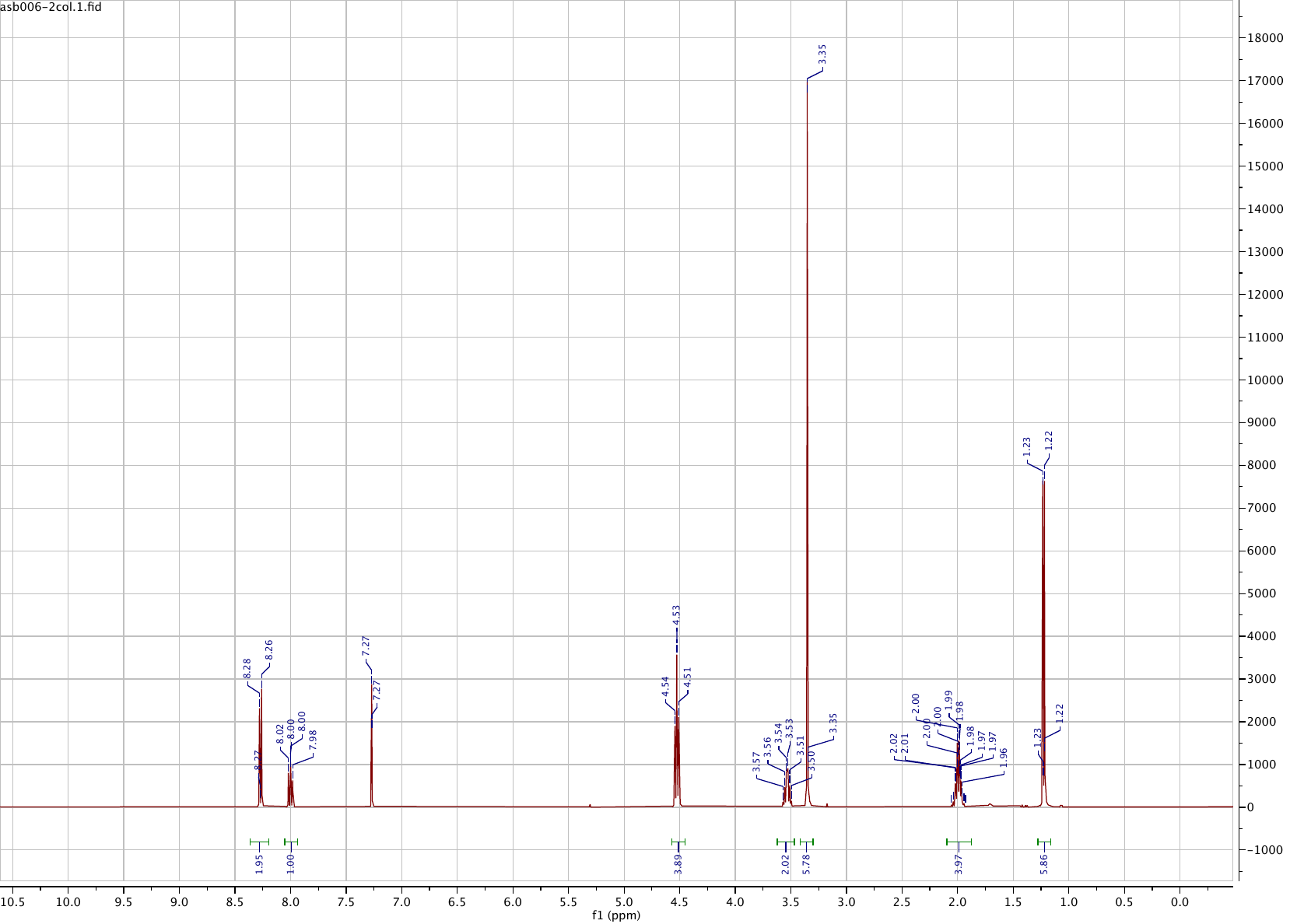

**Compound 21**
